# Criteria for effective fallow field eco schemes for farmland birds during non-breeding

**DOI:** 10.1101/2020.10.07.329847

**Authors:** Mirjam Rieger, Sarah Mailänder, Lea Stier, Julia Staggenborg, Nils Anthes

## Abstract

1. Farmland eco schemes implemented under the current Common Agricultural Policy (CAP) of the European Union are often considered ineffective in halting farmland bird declines. Fallow fields, often seeded with dedicated seed mixtures, rate among the more beneficial eco scheme types. Yet, the CAP currently defines no minimum criteria for fallow fields to qualify as eco scheme, likely jeopardizing their potential biodiversity benefits.
2. We investigated the attractiveness of four fallow field types established under CAP eco schemes and dedicated bird conservation programs in Southern Germany. Our 2-year surveys on > 100 fields focused on the non-breeding season, where food limitation can become particularly problematic. We modelled bird incidences also in response to vegetation structure and adjacent landscape features to derive minimum criteria for effective fallow field eco schemes.
3. Fallow field types varied only mildly in overall species richness but showed striking differences in the attracted species. Finches in particular tended to preferentially visit 1-year fallow fields, while buntings tended towards 2-year and older field types. 1-year CAP fallows, however, are typically removed before mid-winter, and thus potentially act as a trap to farmland birds and other wildlife.
4. The investigated species consistently preferred larger fallow fields with a more differentiated vegetation structure. Placement close to woods and hedgerows positively affected birds inhabiting woodland ecotones, while classic farmland species showed higher incidences on fallow fields embedded in open landscapes.
5. ‘*Policy implications’* Our findings call for the ongoing CAP revisions to specify minimum requirements that qualify fallow fields as eco schemes. These should include an at least biennial cycle, a diversification of seed mixtures, standards for fallow field size, and criteria for their placement in the landscape matrix.

## 1 Introduction

The diversity and abundance of farmland birds across most of Central and Western Europe has severely declined. The European common farmland bird indicator decreased by 57 percent between 1980 and 2016 (EBCC & Birdlife International 2020). This decline has been linked to the ongoing intensification of farming practices under the Common Agricultural Policy (CAP) of the European Union, whose subsidies primarily scale with farm size and productivity (Donald et al. 2002, Pe’er et al. 2014). While CAP payments also require farmers to implement dedicated eco schemes, their effectiveness to halt or even reverse farmland biodiversity declines has been questioned (Pe’er et al. 2014, 2017).

Current CAP eco schemes rest on three components. First, ‘Greening’ measures in CAP pillar I make 30 % of the direct cross compliance payments to farmers dependent on the assignment of Ecological Focus Areas (EFA) on ≥ 5% of their arable land. Second, agri-environment-climate schemes (AECS) offer complementary CAP pillar II funding to member states for farmland-related rural development programs to which farmers can voluntarily subscribe. Third, targeted species protection measures through contractual conservation agreements have been developed by most European countries, again co-funded through CAP pillar II. The third component typically includes highly efficient conservation measures but is implemented on small land fractions, only. In Germany, for example, these measures covered approx. 4,000 km^2^ or 2.4 % of the agricultural area in 2013 (Grajewski & Schmidt 2015). In contrast, the first two components receive substantial funding (approx. 27 % of the annual EU agricultural budget, Pe’er et al. 2020) and affect > 10 % of the agricultural area in Germany (Zinngrebe et al. 2017), thus offering a potential to support biodiversity at relevant spatial scales. Recent work, however, suggests that 75 % of the eco scheme area implemented under current Greening and AECS regulations employs options such as catch crops, green cover, or nitrogen fixing crops that are considered ineffective for biodiversity support (Nitsch et al. 2017, Zinngrebe et al. 2017, Pe’er et al. 2017).

One eco scheme considered generally effective for farmland bird conservation is the designation of fallow fields (here broadly including uncultivated set-aside cropland as well as fields seeded with dedicated ‘wild bird’ or wildflower seed mixtures). Yet, since farmland bird species differ in food preferences and the habitat structures they prefer for shelter, plant species composition and the resultant food supply on different fallow field categories can be expected to differently attract or deter particular bird species (e.g. Perkins et al. 2008). Hence, the conservation value of fallow field eco schemes likely varies with the chosen seed mixtures, age since establishment, vegetation structure, and the composition of the adjacent landscape matrix. This perception contrasts to current CAP eco scheme regulations, which offer only a narrow set of prescribed seed mixtures with (mostly short-term) fallow durations and lack minimum quality requirements with respect to size or placement. To enable a target-oriented revision of CAP eco schemes, our study quantified how fallow field type as well as structure and placement characteristics affect farmland bird incidences.

Our surveys focussed on the non-breeding season, when energy demands to maintain body temperature and metabolism are particularly high. Food limitation in late autumn and winter can thus substantially affect farmland bird condition and survival (Newton 2004, Siriwardena et al. 2008, Siriwardena 2010). The severity of this ‘winter food gap’ further increased with recent agricultural intensification. For example, enhanced harvesting efficiency and the replacement of seed rich winter stubble fields by winter crops and green manure reduce the availability of leftover cereal grains (Atkinson et al. 2002, Moorcroft et al. 2002, Newton 2017). Moreover, the widespread application of herbicides combined with the abolishment of set-aside requirements in the CAP 2008 reform reduced the availability of fields rich in weed seeds, and thus a prime foraging resource for granivorous birds (Marshall et al. 2003, Gillings et al. 2010).

In winter, fallow fields are well known to meet foraging and shelter requirements of many farmland birds (Gillings et al. 2010, Kasprzykowski & Goławski 2012, Joest et al. 2016, Dellwisch et al. 2019). Yet, only few studies to date differentiated bird habitat use within the ‘fallow’ category, i.e. among different types and ages of fallow fields (e.g. Buckingham et al. 1999, Orłowski 2006, Perkins et al. 2008, Birrer et al. 2018). Their findings varied substantially, likely because of differences in the investigated fallow field types, seed mixtures and surveyed regions, calling for further studies to better substantiate decisions about management strategies. Likewise, only few earlier studies investigated effects of fallow field vegetation structure and placement on bird abundance (e.g., Henderson et al. 2004, Stoate et al. 2004).

We investigated habitat use across fallow field types of different age and origin in an agricultural landscape of SW Germany. Our findings can inform the ongoing reform of CAP eco schemes for the upcoming funding period, and help conservation practitioners to select particularly suitable fallow land schemes to support their respective core target species.

## 2 Methods

We studied habitat use by 17 bird species known to use fallows for foraging or shelter (Tab. 2). These fall into two guilds containing typical inhabitants of woodland ecotones and of semi-open to open farmlands, respectively (Tab. 2). All species except chaffinch, great tit, and blue tit experienced significant population declines in Europe (EBCC & Birdlife International 2020) and / or Germany (Gerlach et al. 2019) over the past three decades, and nine species are integrated into the European farmland bird index (Tab. 2).

**Table 1.**
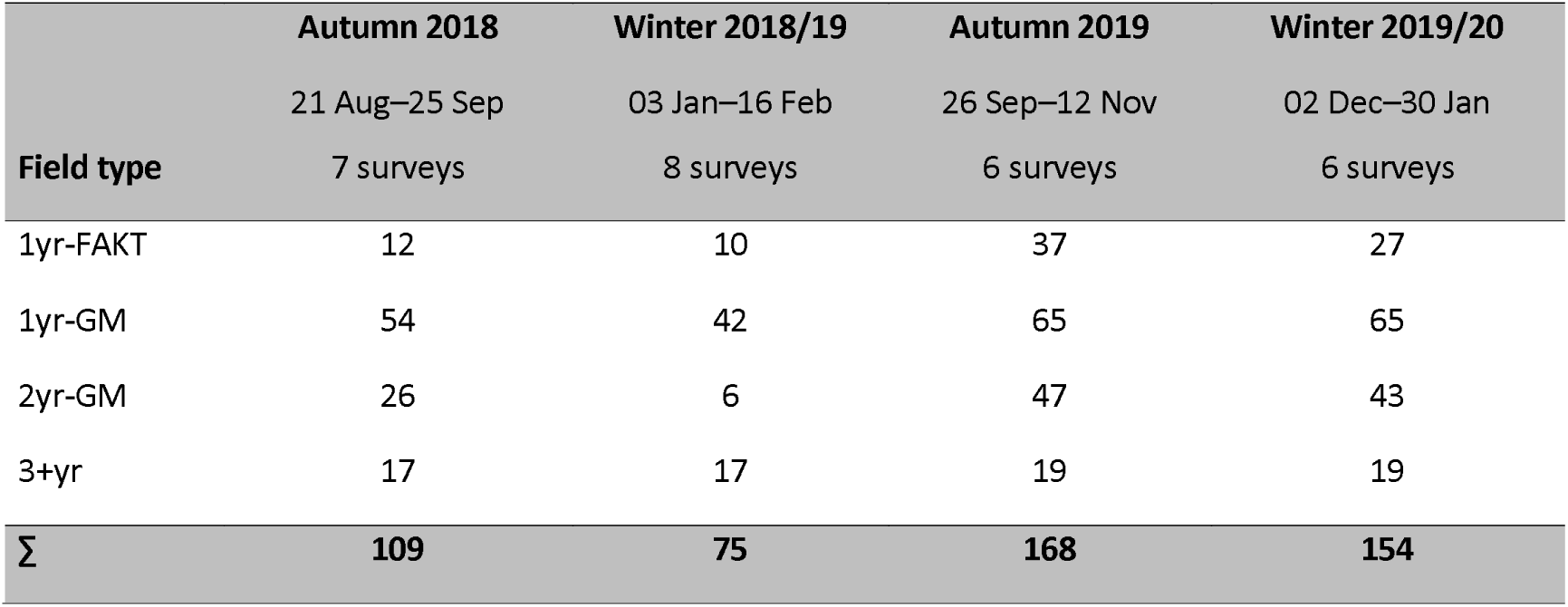
Survey periods, the number of replicate surveys, and the number of fields surveyed per fallow field type and investigated season.

|  | Autumn 2018 | Winter 2018/19 | Autumn 2019 | Winter 2019/20 |
| --- | --- | --- | --- | --- |
|  | 21 Aug–25 Sep | 03 Jan–16 Feb | 26 Sep–12 Nov | 02 Dec–30 Jan |
| Field type | 7 surveys | 8 surveys | 6 surveys | 6 surveys |
| 1yr-FAKT | 12 | 10 | 37 | 27 |
| 1yr-GM | 54 | 42 | 65 | 65 |
| 2yr-GM | 26 | 6 | 47 | 43 |
| 3+yr | 17 | 17 | 19 | 19 |
| **Σ** | **109** | **75** | **168** | **154** |

**Table 2.**
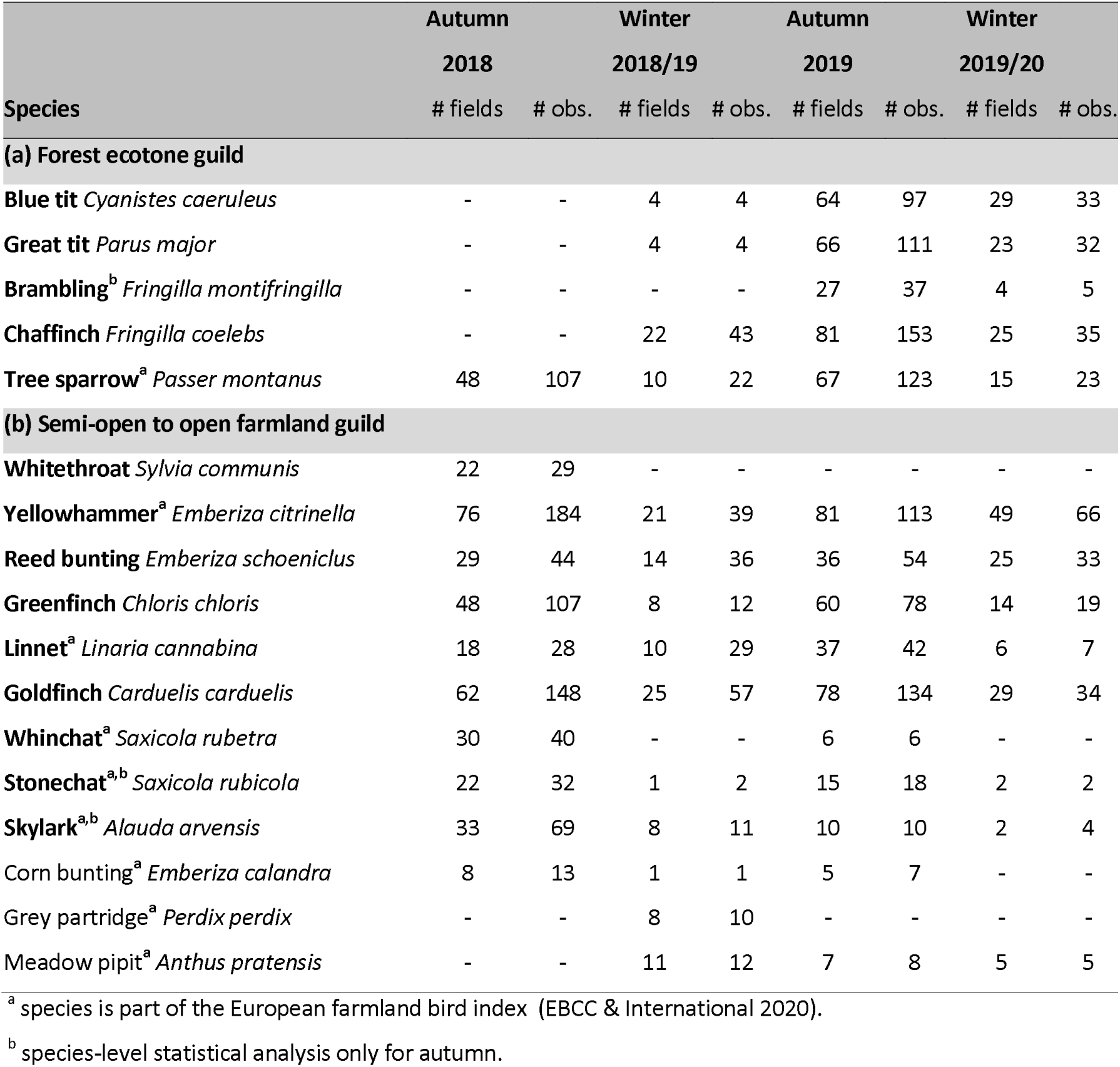
Number of sites with sightings (# fields) and total number of sightings (# obs.) per species per season. Species in bold face were used for a species-level statistical analysis.

### 2.1 Study area

We surveyed nine study areas (A to H in Fig. 1) at 330–400 m a.s.l. in an agricultural landscape in the centre of the SW German federal state of Baden-Württemberg. The study region includes floodplain soils in the valleys of the Neckar and Ammer rivers, and highly fertile loess soils on the adjacent plateaus. Most study areas are under intense agriculture, but study areas D, E and I (in Fig. 1) share exceptionally high baseline proportions of organic farming. All areas have seen an increasing establishment of annual fallows under EU CAP regulations, and perennial fallows in the context of governmental grey partridge and corn bunting conservation projects since 2014.

**Figure 1.**
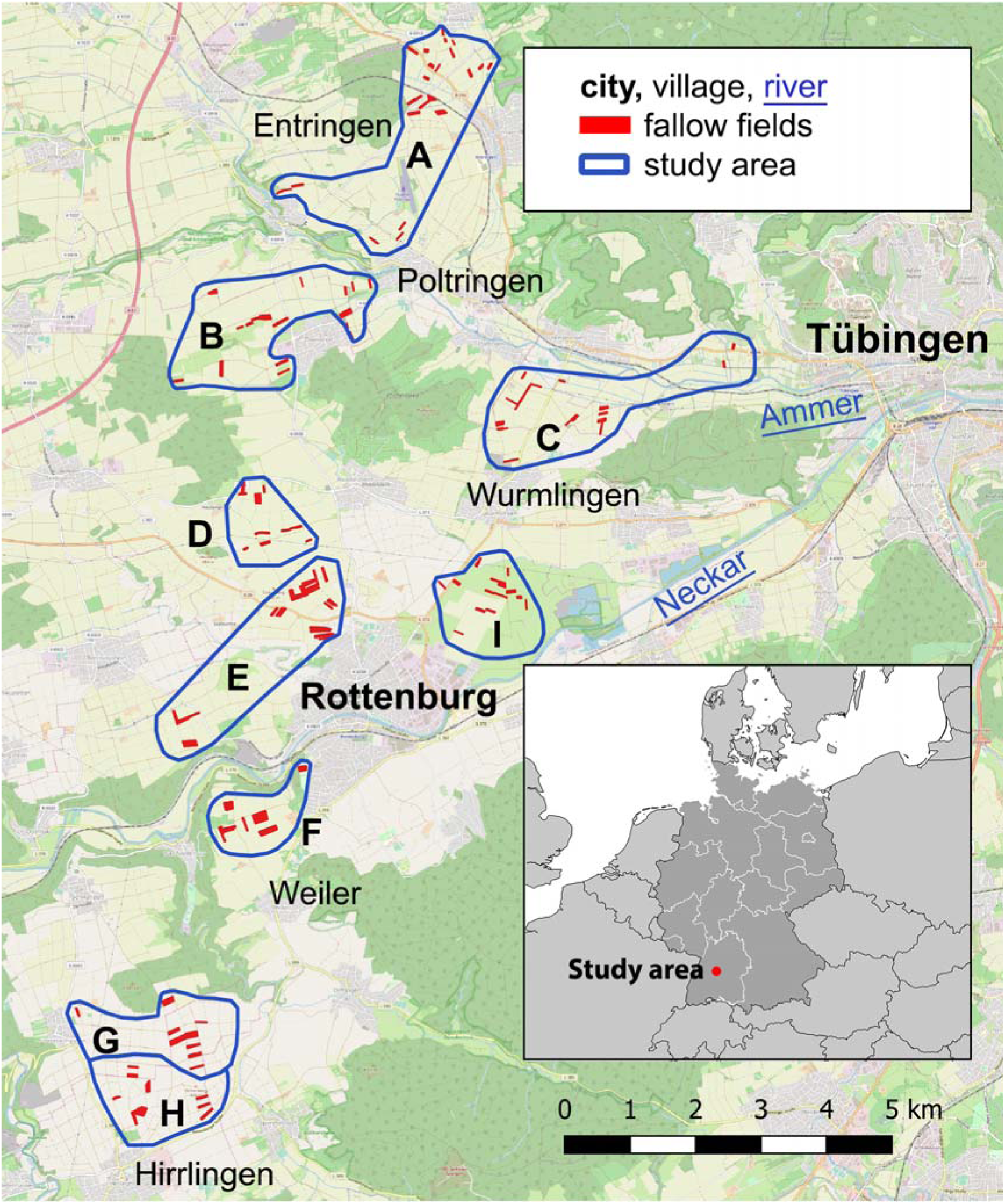
Localisation of the nine study areas and all fallow fields investigated in at least one season (sample sizes in Table 1). A – Ammerbuch, B – Oberndorf, C – Ammer Valley, D – Heuberg North, E – Heuberg South, F – Weiler, G – Hirrlingen North, H – Hirrlingen South, I – Neckar Valley. Map source: OpenStreetMap.

### 2.2 Fallow field types and their characteristics

We distinguished four fallow field types based on seed mixture (details in Online Supplement A) and fallow age, with sample sizes given in Table 1.

(i) **‘1yr-FAKT’** subsumes spring-sown annual fallows that qualify as ecological focus areas (EFA) under EU greening regulations, or as a regional eco scheme under the federal government’s AECS-program ‘FAKT’ (BMEL 2015, MLR 2017). Seed mixtures have high shares of annual plants such as sunflower *Helianthus annuus*, blue tansy *Phacelia tanacetifolia*, and ramtil *Guizotia abyssinica* (Fig. 2a). 1yr-FAKT fallows are often removed in September or November (LTZ 2020) to prepare the field for the subsequent winter culture.

**Figure 2.**
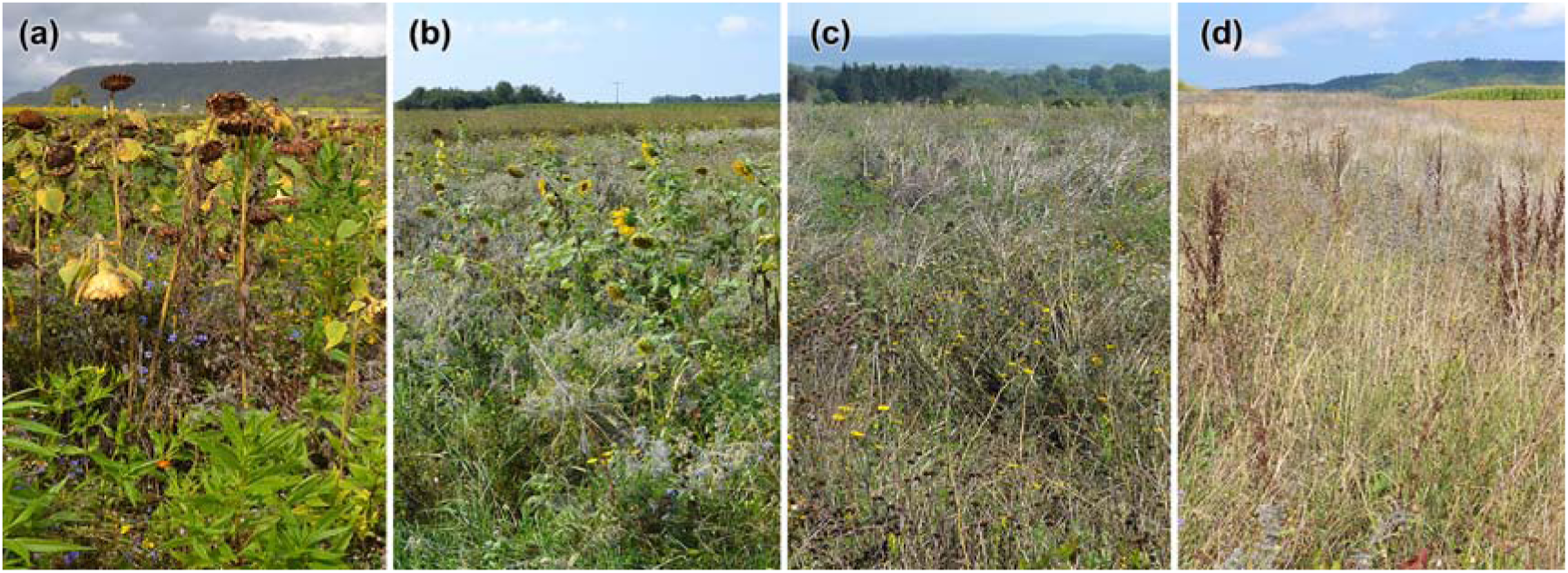
Examples of the four surveyed fallow field types: (a) 1yr-FAKT (25 Sep 2019), (b) 1yr-GM (27 Aug 2019), (c) 2yr-GM (30 Aug 2019), (d) 3+yr (11 Sep 2019). Photo © Sarah Mailänder.

(ii + iii) ‘**GM’** subsumes fallow fields sown with the so-called Göttingen mixture (GM), a perennial seed mix optimized for grey partridge conservation (Gottschalk & Beeke 2014). It has a rather balanced mix of annual and perennial plants, usually resulting in structurally rich fields. Yet, dominance may occur by mallow *Malva sylvestris* and sunflower in year 1, or yellow sweet clover *Melilotus officinalis* in year 2. GM fallows are implemented via contractual species conservation agreements under the regional Landscape Conservation Directive (Landschaftspflegerichtlinie, Az.: 73-8872.00). In a biennial alternated rotation, half the field is ploughed in winter and newly sown in spring, the other half left untouched until the subsequent winter. Hence, at any time, half the field is in its first year since sowing (**1yr-GM**, Fig. 2b), the other half in its second year (**2yr-GM**, Fig. 2c).

(iv) ‘**3+yr’** subsumes perennial fallows established under a diverse range of conservation activities (Fig. 2d) that were sown at least three years prior to our surveys. Seed mixtures cannot be reconstructed for all those sites, but about half were initially sown with ‘Lebensraum-I-Tübingen’ (Online Supplement A). 3+yr fallows vary in structure and plant composition, but often show high coverage of grasses, teasel *Dipsacus fullonum*, wild carrot *Daucus carota*, tansy *Tanacetum vulgare* and upcoming shrubs.

To understand how bird incidence varies with characteristics beyond fallow field type, we recorded the following covariates for each surveyed fallow field:

i. *Fallow field size* (in ha, range 0.06-0.70 ha, with extremes reaching 2.80 ha).
ii. *Vegetation height* (in 10 cm steps, measuring the dominant vegetation layer).
iii. *Secondary plant layer* (presence/absence), as generated by exposed bud stands of perennials such as sunflower, fennel *Foeniculum vulgare* or wild teasel, indicating structural diversity.
iv. *Dominant plant species* (presence/absence, with at least 1 species reaching a minimum coverage of 30%).
v. *Bare ground* coverage (3 categories: 0-10%, 10-30%, and >30% of accessible soil representing potential foraging habitat).
vi. *Farm track* (presence/absence) along the longer edge of the fallow field, potentially disturbing roosting birds.

Finally, we predicted bird presence to vary with landscape features directly adjacent to a fallow field. We quantified coverage of alternative roosting and foraging structures within a 100 m buffer around each surveyed fallow field from aerial images (Bing maps) using QGIS version 3.2. From this we derived proportional area coverages for other fallow fields (*Fallow coverage*) and for hedges, tree clusters and orchards (*Hedge and grove coverage*).

### 2.3 Bird surveys

Surveys covered autumn and mid-winter periods, each replicated twice in the seasons 2018/19 and 2019/20 (Table 1). We recorded the presence and individual numbers of each target species per survey round and fallow field (overpasses ignored). To exclude confounding sequence effects, we randomized starting time, walking direction and the order in which fallow fields were visited across consecutive surveys in a given study area. We spread surveys roughly uniformly across daytimes between 30 min after sunrise and 30 min before sunset and excluded periods with continuous rainfall or winds exceeding 40 km/h. We slowly passed along the long edge of each fallow field, entering it occasionally to also detect birds hiding in the vegetation. To avoid double-counts we visually traced flushed birds to their new location, recording only the fallow field of first encounter. Fields that were mulched or ploughed within a given survey period were excluded when only a single survey had been possible.

### 2.4 Statistical analysis

Statistical analyses for species-specific habitat use per season included all species with > 10 sightings that spread across at least 15 percent of the visited fallow fields (Table 2).

Separately for each bird species, but combined across years for autumn and winter season (Table 1), we modelled bird occurrences using generalized linear mixed models (GLMM). Bayesian estimation procedures used the Stan C++ library as implemented in the rstanarm package version 2.19.2 (Goodrich et al. 2020) for R version 3.6.2 (R Core Team 2017). The response variable in species-level models was derived from the number of surveys with and without recordings (= successes and failures) using the binomial family and a logit-link, resulting in ‘bird incidence’ as the probability to encounter a species on a given fallow field. We separately calculated models with the number of sighted species (‘species number’) as our response variable, now using the Poisson family and a log-link.

In all models, *fallow field type* served as our key predictor variable, with four levels as defined above. The models further contained fixed covariates to account for variation associated with fallow field characteristics beyond field type, as specified in section 2.2. Continuous covariates included *fallow field size, fallow coverage, hedge and grove coverage*, and *vegetation height*. The first three of these were log10(x + 0.01)-transformed to approach normality, and all four *z*-transformed prior to analysis. Categorical covariates included *farm track* presence, *secondary plant layer* presence, and survey *year* (2018/19, 2019/20). The analysis of winter species numbers further contained *survey completeness* (0/1) as a covariate, where ‘0’ identifies 1yr-FAKT and 2yr-GM fallow fields that were ploughed already before the end of our successive surveys. We excluded the covariates *bare ground* and *dominant plant species* because of strong covariation with fallow field type.

While the combined surveys on a given fallow field per season and year represent our raw data points, these suffer from some degree of spatial non-independence, where fields from the same study area are more likely to be visited by the same local bird population. We therefore added *study area ID* (Fig. 1) as a random intercept to each model. Study areas without any record of a given species during a particular season were excluded from the analysis. We further included an observation-level random intercept, successfully treating overdispersion (Harrison 2014).

We initiated Bayesian models with weakly informative prior distributions (means ± SD for intercepts 0 ± 10 and for coefficients 0 ± 2.5, respectively) and used posterior predictive model checking to assess model fit and an adequate representation of observed zero counts. For each model, we ran 10 MCMC chains from which we retained 3000 post-warm-up samples. We then derived covariate coefficient estimates (means ± 95% credible intervals, CrI) from the combined 30,000 posterior samples. From these, we also derived posterior probabilities (so-called Bayes *P*) that coefficient estimates – as well as pairwise differences between our four fallow field type levels – exceed 0. The explanatory power of a given model covariate increases when its posterior probability approaches 0 or 1. Most covariate estimates achieved reliable conversion indicators (Korner-Nievergelt et al. 2015), namely 5000 effective posterior samples (observed min 4344), Rhat values ≤ 1.02 (max 1.002) (Brooks & Gelman 1998), and Monte Carlo standard errors (MCSE) ≤ 2 % of the standard deviation (max 1.5%). For graphical displays, we extracted model predictions and their CrI of the predictor of interest for defined values of the remaining model predictors, i.e. continuous covariates set to their sample mean, and factors set to a single level when predicting for covariates (Korner-Nievergelt et al. 2015).

## 3 Results

Out of 17 target species, 16 and 15 were recorded at least once during the autumn and winter season, respectively (Table 2), with clearly higher average species numbers per fallow field in autumn (Fig 3).

**Figure 3.**
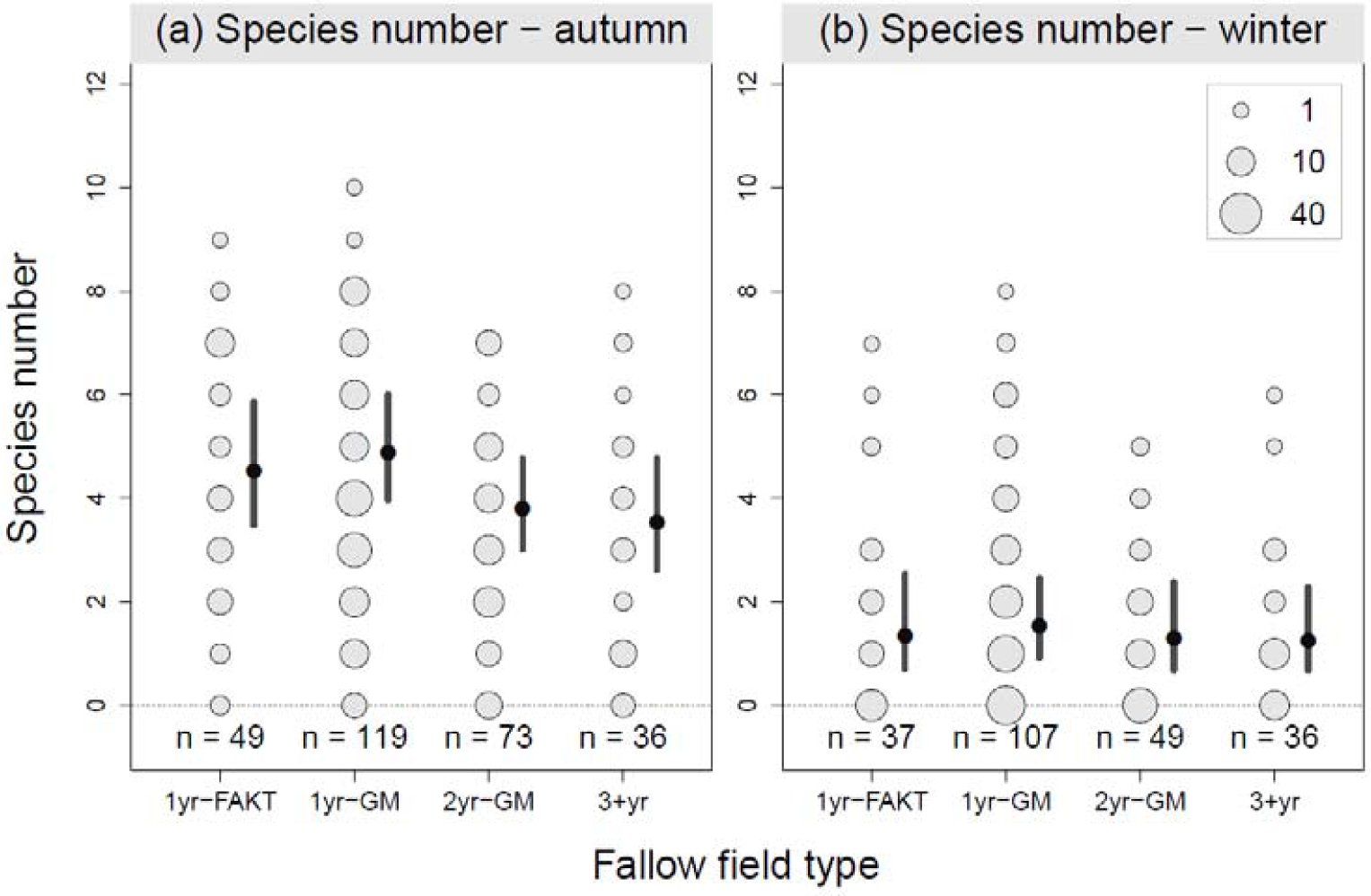
Observed variation in species number in (a) autumn and (b) winter between fallow field types. Grey dots show raw data and scale with the number of fallow fields. Black dots and flags show model-predicted means and their 95% CrI for the following co-variate levels: survey completeness = yes, year = 2019/20, farm track = no, secondary plant layer = yes. For detailed statistical results, see Online Supplements B and C.

### 3.1 Fallow field types

Average species numbers varied only little between fallow field types. In autumn, even the strongest contrasts yielded differences in mean species numbers of only 0.7 to 1.3 between 1yr-FAKT and 1yr-GM fields versus 2yr-GM and 3+yr fields, with substantial overlap in credible intervals (Fig 3a, full statistical detail in Online Supplements B and C). Mean species numbers were clearly lower in winter, and very similar in all four fallow field types (Fig 3b, Online Supplements B and C).

Individual species exhibited more differentiated patterns in their use of fallow field types. We found a particularly consistent contrast in the usage of 1-year fallow fields (1yr-FAKT and 1yr-GM) compared to later successional stages (2yr-GM and 3+yr), but no further differentiation within each of these two groups. All five surveyed finch species, great tit and tree sparrow – at least in autumn and then coupled with an undifferentiated pattern in winter – showed higher incidences in 1-year compared to older fallow fields, as exemplified in Fig 4a-c (full statistical detail and graphical displays for all study species in Online Supplements B, C, D). The reverse pattern consistently occurred in yellowhammer, reed bunting or whinchat, all showing higher incidences in older fallow fields compared to their first year counterparts (Fig 4d-f, full statistical detail in Online Supplements B, C, D).

**Figure 4.**
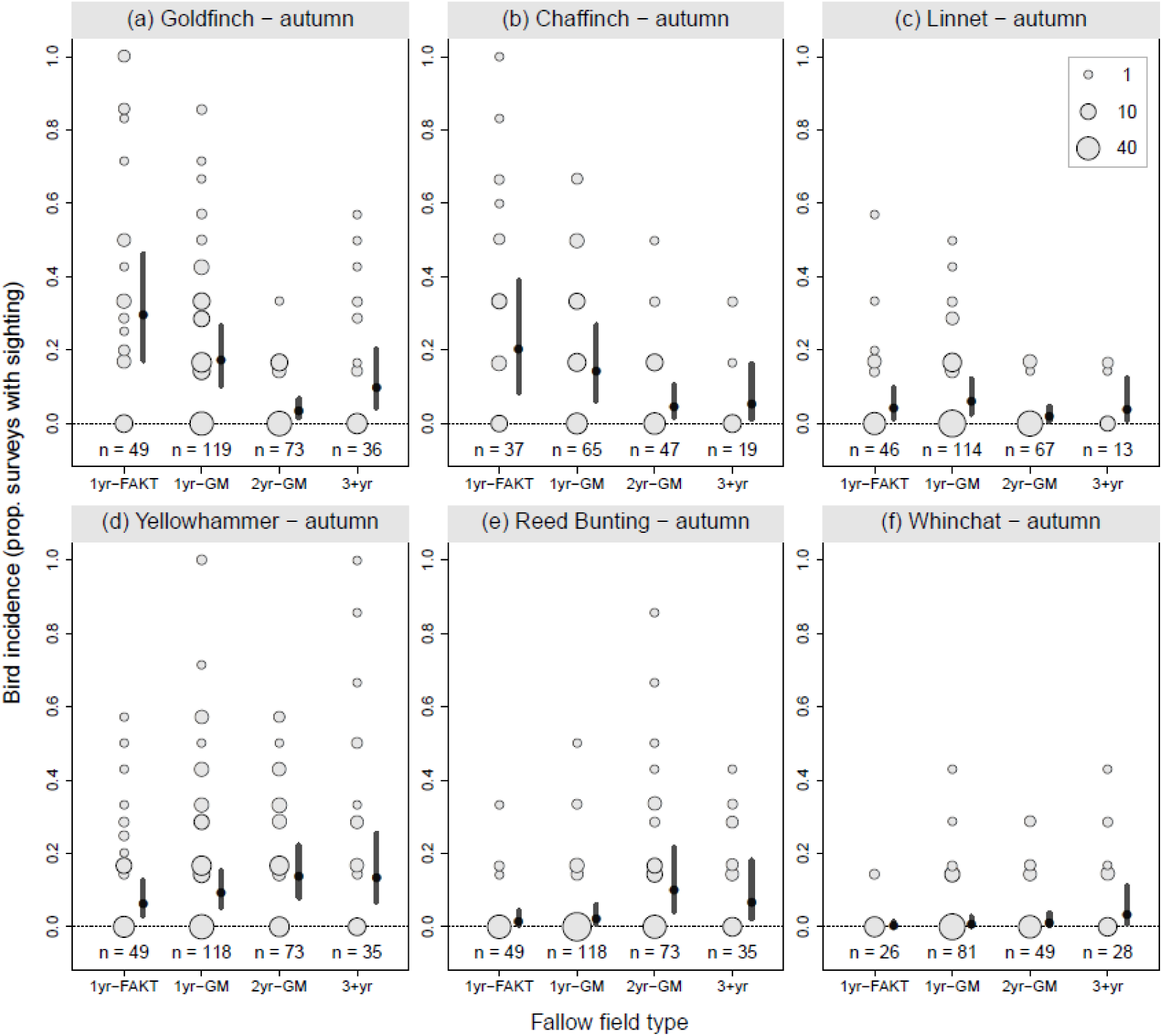
Variation in bird incidences between fallow field types. Examples illustrate strong (goldfinch, a chaffinch, b) and weak (linnet, c) preferences for 1-st year fallows, and strong (yellowhammer, d, reed bunting, e), and weak (whinchat, f) preferences for later successional stages. Grey dots show raw data and scale with the number of fallow fields. Black dots and flags show model-predicted means and their 95% CrI for the following co-variate levels: year = 2019/20, farm track = no, secondary plant layer = yes. Detailed results and displays for all study species in Online Supplements B, C, and D.

### 3.2 Fallow field characteristics

Several fallow field characteristics exhibited rather universal co-variation with the total number of recorded species per site as well as species-specific bird incidence. Fallow field size showed a consistent positive, and often strong, relationship with both response measures (Fig 5a). We found similar but slightly more variable positive relationships for fallow fields with taller vegetation (Fig 5b) and – in particular during winter – for the presence of a secondary plant layer (Fig 5e), indicating benefits to a structurally diverse fallow field vegetation.

**Figure 5.**
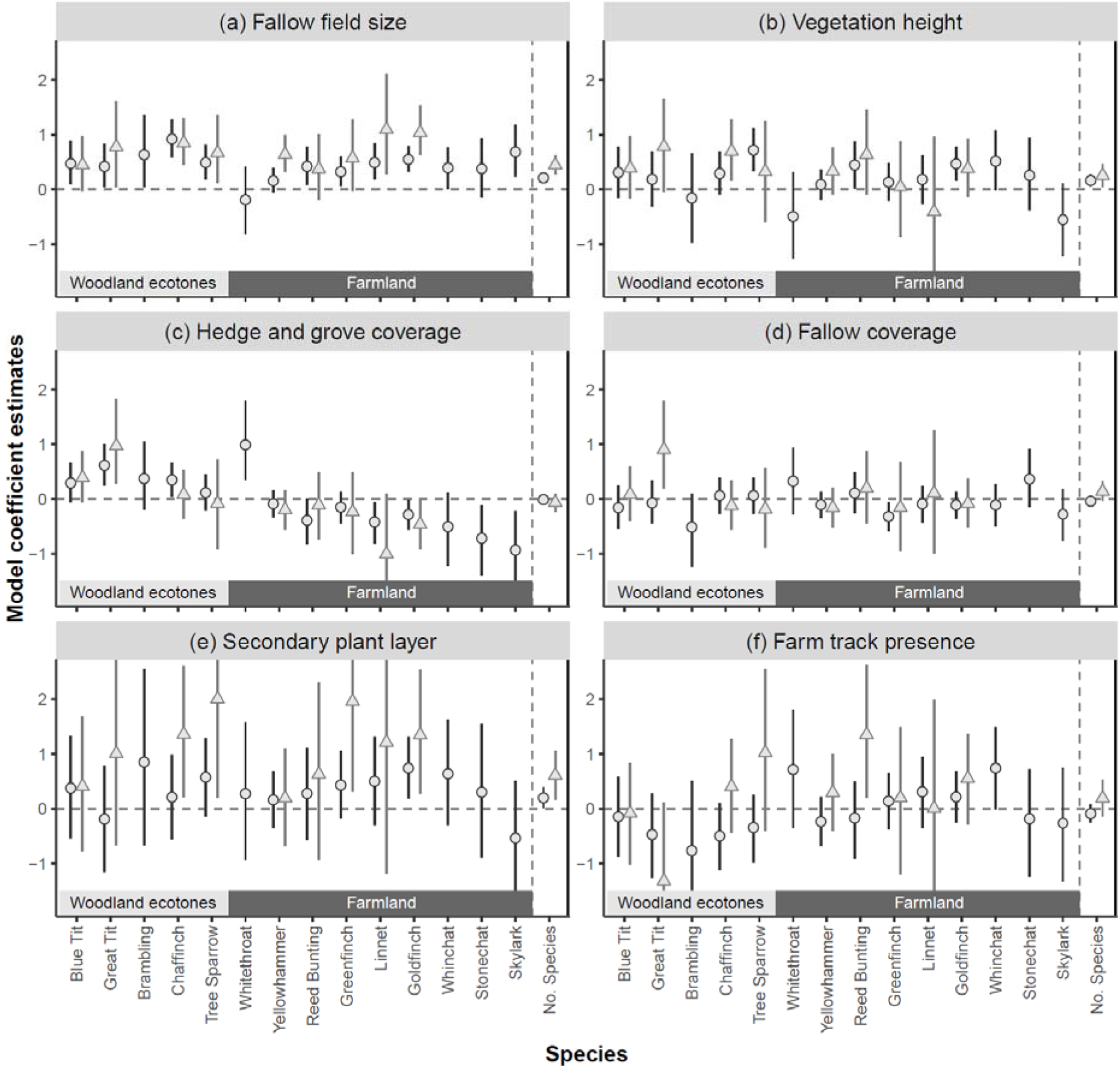
Covariates estimates with their 95% CrI for each species, separated by autumn (black circles) and winter (grey triangles) surveys, and grouped by woodland ecotone versus farmland inhabitants. Full coefficient estimates are given in Online Supplement B.

Contrary to these consistent patterns, species varied strikingly in their predicted response to the availability of hedges and groves nearby fallow fields (Fig 5c). While inhabitants of woodland ecotones such as great tit, chaffinch or blue tit tended towards positive responses, farmland birds such as goldfinch, linnet, skylark or stonechat showed higher incidences on fallow fields with little or no hedges and groves nearby, indicating avoidance reactions. Given these differential species responses, overall species richness varied largely independent of this covariate (Fig. 5c).

In the majority of cases, the availability of other fallow fields in the near surroundings unfolded no striking correlations with species presence (Fig 5d). Similarly, the presence or absence of a farm track along the long side of a fallow field yielded no consistent patterns with respect to bird presence or species richness (Fig 5f).

## 4 Discussion

Our study illustrates how fallow field eco schemes differentially support Central European farmland birds. This differentiation did not manifest at the level of overall species richness, which distributed rather uniformly across fallow field types. Instead, we document substantial variation in the degree to which individual species made use of the four investigated fallow field types. The attractiveness of fallow fields further varied strongly with several field characteristics, in particular their size, vegetation structure, and integration into the landscape matrix.

### 4.1 Fallow field types

A major shift in the supported bird community occurred between 1-year and 2-year and older fallow field types. 1-year fallow fields primarily attracted granivorous birds foraging on large seeds picked from the exposed buds of e.g. sunflowers or bare ground (Henderson et al. 2004), with finches as prime representatives of this guild. 2-year and older fallow fields rather attracted granivores foraging well hidden in the dense and highly structured vegetation on smaller seeds, with buntings as prime representatives.

We interpret this pattern to arise from three interacting factors. First, the seed mixtures used in the majority of our investigated fallow fields (Online Supplement A) contain large fractions of – mostly cultivated – annuals that provide many large seeds from, for example, sunflower, borage *Borago officinalis* or linseed *Linum usitatissimum*. In their second year, these fields tend to develop clearly lower flowering densities, resulting in lower seedset, with the latter more dominated by perennials such as yellow clover, yellow chamomile *Anthemis tinctoria* or tansy with rather small seeds. Second, our 1-year fallow fields tended to produce tall stands with little differentiation in vegetation heights and hardly any ground cover by segetal flora or grasses, thus offering only limited shelter for species that preferentially forage in the ground vegetation. This differentiation increases from 2-year onwards, providing suitable habitat structure for a wider range of bird species. Finally, immigrating weeds such as goosefoot *Chenopodium album*, mugwort *Artemisia vulgaris*, red-root amaranth *Amaranthus retroflexus* or thistle *Cirsium arvense* increase vegetation differentiation in 2-year and older fallows, and enhance the availability of small seeds.

Earlier studies also found diverging winter bird communities between younger and older fallow fields, but differences in the investigated plant compositions and age-classes make comparisons difficult. In Poland, for example, *Cirsium*-rich young fallows (< 3 years) were preferred by granivores and *Tanacetum*-rich permanent fallows (> 3 years) by insectivores (Orlowski 2006). In Great Britain, skylark, finches and yellowhammer showed more consistent preferences for 1-year than to ≥ 2-year fallows when compared to other cropland types (Buckingham et al. 1999). In Switzerland, meadow pipit and goldfinch preferred ≤ 2-year wildflower strips rich in wild teasel and wild carrot, whereas yellowhammer preferred perennial (14-year on average) fallow fields rich in *Rubus* shrubs (Birrer et al. 2018).

While our data may seem to suggest that 1-year fallows generally represent attractive foraging habitats and thus high quality eco schemes for farmland birds, only 1yr-GM-types – in stark contrast to 1yr-FAKT-types – can reliably unfold their ecological service. This contrast is rooted in current CAP regulations, which allow 1yr-FAKT type eco schemes to be mown and ploughed already between September and November of the year of sowing (e.g. LTZ 2020). As a consequence, these fallow fields are not available as foraging habitat or shelter during mid and late winter when food restriction culminates (Siriwardena et al. 2008). Such eco schemes may therefore act as a trap for overwintering birds, in particular in arable landscapes that lack local escape options at the time of ploughing. In contrast, GM-type eco schemes are maintained into at least their second year before ploughing, and are then adjacent to a 1-year field section that can act as a local refuge into spring (Gottschalk & Beeke 2014). This failure of 1yr-FAKT type eco schemes unfolds large scale consequences: Compared to more targeted (and then typically more effective) contractual conservation programs, this scheme alone covers a substantially larger land fraction, in Germany alone approx. 2330 km^2^ or 1.4 % of the agricultural area in 2019 (DBV 2019).

### 4.2 Fallow field characteristics

Beyond fallow field type, we found higher overall species richness and incidences of individual bird species on large fields with a tall and structurally diverse vegetation. This implies that eco schemes should prioritize large fallow patches, and seed those with mixtures that develop into structurally diverse and seed rich fallows (see section 4.3). We propose to derive minimum size standards from earlier suggestions that the width of fallow fields should exceed 12-20 m to minimize predation risks on breeding farmland birds such as Grey Partridge (Bro et al. 2004, Gottschalk & Beeke 2014). For a field of 150 m length, this results in minimum fallow field sizes of 0.15-0.3 ha.

The placement of fallow fields within the landscape matrix exhibited species-specific links with bird incidence. Inhabitants of woodland ecotones (Table 2) preferentially visited fallow fields adjacent to (tall) hedges, orchards or groves. This is consistent with earlier findings that many granivores preferentially forage on arable fields along hedgerows, where seed availability is enhanced and shelter nearby (e.g. Robinson & Sutherland 1999, Stoate et al. 2004, Dellwisch et al. 2019). In contrast, classic farmland birds tended to show higher incidences on fallow fields embedded in a rather open landscape matrix. Avoidance reactions towards woodland-type habitats are well-known for skylark, but have occasionally been documented also for other farmland birds such as goldfinch, linnet and buntings (Parish et al. 1995, Robinson & Sutherland 1999, Donald et al. 2001). We cannot differentiate whether our observed patterns arise from avoidance reactions or a lower likelihood to observe birds in a given fallow field when they may also just roost in a nearby grove. The latter appears plausible for yellowhammer, greenfinch, and goldfinch, which show lower incidences on fallow fields close to groves in the current study, but have been found to frequently roost in tall hedges in an earlier survey in the same study area (Dellwisch et al. 2019). In summary, fallow fields seem to represent attractive winter foraging habitat irrespective of their placement relative to hedges and groves, but with differences in the supported bird communities. Hence, to serve both requirements, effective eco schemes should assure that fallow fields are not placed solely at marginal yield locations close to forests or tall hedgerows, but also integrated into the widely open agricultural landscape.

### 4.3 Seed availability, winter food gap and seed mixtures

We did not quantify seasonal variation in seed availability in flower heads or soil samples on our study sites. Yet, haphazard surveys of seed loads in sunflowers or teasel imply substantial depletion towards mid-winter, consistent with previous research (e.g. Boatman et al. 2003, Henderson et al. 2004, Perkins et al. 2008, Geiger et al. 2014). Fallow field eco schemes may therefore fail on winter food provision as one of their key goals, with established negative effects on buntings, sparrows, or skylark (Peach et al. 1999, Siriwardena et al. 2008). Mitigation can pursue three approaches. First, fallow field eco schemes should explicitly integrate seed mixtures sown in late summer instead of spring, then enhancing chances that annuals set seed late and maintain seed provisioning until late winter.

Second, seed mixtures should increase the share of plants that retain seed into late winter. Promising candidate species include kale *Brassica oleracea*, quinoa *Chenopodium quinoa*, linseed, triticale *Tritico secale* or millet, while sunflower, borage, oat *Avena sativa* or barley *Hordeum vulgare* deplete seeds already in autumn (Boatman et al. 2003, Henderson et al. 2004, Perkins et al. 2008). This coincides with studies identifying kale, quinoa, linseed and several cereals as particularly relevant food sources for granivorous buntings and finches (Stoate et al. 2003, Henderson et al. 2004, Stoate et al. 2004, Perkins et al. 2008). We propose that these species should partially replace the substantial seed mixture shares of plants with minimal role for foraging farmland birds, such as phacelia, buckwheat or yellow sweet clover (Marshall et al. 2003, Henderson et al. 2004, Perkins et al. 2007), consistent with their representation in wild bird mixtures for the British environmental stewardship program (Boatman et al. 2003).

Third, we propose to enhance regional AECS-based funding for unharvested cereal crops (so-called ‘seeding cereals’ in Henderson et al. 2004), which represent a prime winter food for many granivorous birds. Such cereal-biased options are central to the British environmental stewardship program (e.g. Natural England 2013) and available also in some German Federal States (e.g. Joest et al. 2016), but insufficiently attractive to farmers under the current subsidy system in Baden-Württemberg. Buntings and sparrows in particular profit from the high abundance of cereals, for example on unharvested crop mixtures (Holland et al. 2006, Joest et al. 2016, Perkins et al. 2008).

### 4.4 Conclusions

This study supports the hypothesis that farmland birds substantially vary in habitat use among available fallow field types, both between species but also between seasons within single species. We suggest that the ongoing CAP reform formulates explicit minimum standards to qualify fallow fields as effective eco scheme options. A large-scale application of just a few types of fallow field seed mixtures and age as supported under current CAP regulations clearly fails to support the full diversity and demands of farmland birds in winter (Hinsley et al. 2010, Redhead et al. 2018). CAP eco schemes should therefore integrate diversification of seed mixtures (in particular to include plants retaining seed into late winter) and successional stages (e.g., 2-year-rotational and perennial) within a local context. While 1-year fallow fields provide attractive foraging habitat, CAP eco schemes need to integrate a biennial cycle as a minimal requirement, with no ploughing before April of year 2 (Siriwardena 2010). Finally, CAP eco schemes need to define minimum standards for fallow field size and criteria for their placement in the landscape matrix.

## Authors’ contributions statement

NA and JS conceived the study, and jointly with LS, MR and SM designed its methodology; MR, SM and LS collected the data; MR, SM and NA analysed the data; MR, SM and NA led manuscript drafting. All authors contributed critically to the drafts and gave final approval for publication.

## Acknowledgments

The exceptional land fractions devoted to farmland bird conservation measures in our study area originates from joined efforts of conservation practitioners (Sabine Geissler-Strobel and the IAN initiative), local nature conservation and agriculture authorities (Thorsten Teichert, Maik Klingele), and the farmers providing their land. Thanks to Matteo Santon for discussion of the statistical approaches, and to Allan Perkins for comments on an earlier manuscript draft. JS was supported by Stiftung Naturschutzfonds Baden-Württemberg (Az. 73-8831.21/54691-1749L), fieldwork by Carl ZEISS sports optics.

## Online supplements

**Online Supplement A**. Seed mixtures.

**Online Supplement B**. Bayesian model coefficients.

**Online Supplement C**. Pairwise comparisons between fallow field types.

**Online Supplement D**. Covariate raw data plots for all investigated species.

**Data available via the Dryad Digital Repository:**

DOI https://doi.org/10.5061/dryad.1rn8pk0rt

