## Supplement A for "Criteria for effective fallow field eco schemes for farmland birds during non-breeding"

**Online Supplement A**. Seed mixtures used in the investigated fallow fields, showing the proportional seed weights.

| Plant species name | Origin | Lebensraum-1-TÜ | FAKT - M1 | FAKT - M2 | GM |
|---|---|---|---|---|---|
| *Anethum graveolens* | Cultured | - | 2.0 | 2.0 | - |
| *Avena sativa* | Cultured | - | - | - | 5.0 |
| *Borago officinalis* | Cultured | 1.0 | 2.0 | 3.0 | 5.0 |
| *Brassica oleracea* | Cultured | - | - | - | 0.5 |
| *Calendula officinalis* | Cultured | - | 3.0 | 6.0 | - |
| *Centaurea cyanus* | Cultured | 0.5 | 6.0 | 6.0 | - |
| *Coriandrum sativum* | Cultured | - | 3.0 | 5.0 | - |
| *Fagopyron esculentum* | Cultured | 7.5 | 22.5 | - | 14.0 |
| *Foeniculum vulgare* | Cultured | 5.0 | 5.0 | 5.0 | 5.0 |
| *Guizotia abyssinica* | Cultured | - | 2.0 | 7.5 | - |
| *Helianthus annuus* | Cultured | 0.5 | 12.0 | 17.0 | 15.0 |
| *Lepidium sativum* | Cultured | - | - | - | 0.5 |
| *Linum usitatissimum* | Cultured | 8.0 | 4.0 | 10.0 | 15.0 |
| *Lotus corniculatus* | Cultured | 2.0 | - | - | - |
| *Medicago sativa* | Cultured | 8.0 | - | - | 7.0 |
| *Melilotus officinalis* | Cultured | - | - | - | 3.0 |
| *Onobrychis viciifolia / arena* | Cultured | 21.0 | 5.0 | 5.0 | 4.0 |
| *Petroselinum sativum / cris* | Cultured | 1.0 | - | - | - |
| *Phacelia tanacetifolia* | Cultured | - | 10.0 | 12.0 | 7.0 |
| *Raphanus sativus* | Cultured | - | 2.0 | - | 7.0 |
| *Secale multicaule* | Cultured | - | - | - | 5.0 |
| *Sinapis alba* | Cultured | - | 2.0 | - | 1.0 |
| *Trifolium incarnatum* | Cultured | - | 8.0 | 10.0 | - |
| *Trifolium pratense* | Cultured | 5.0 | - | - | 1.0 |
| *Trifolium resupinatum* | Cultured | - | 5.0 | 5.0 | - |
| *Vicia sativa* | Cultured | 3.0 | 6.0 | 6.0 | - |
| *Vicia villosa* | Cultured | 5.0 | - | - | - |
| *Achillea millefolium* | Wild | 1.0 | - | - | - |
| *Anthemis tinctoria* | Wild | 1.0 | - | - | 1.0 |
| *Artemisia vulgaris* | Wild | 0.1 | - | - | - |
| *Barbarea vulgaris* | Wild | 1.0 | - | - | - |
| *Carum carvi* | Wild | 2.5 | - | - | - |
| *Centaurea jacea* | Wild | 1.8 | - | - | - |
| *Centaurea scabiosa* | Wild | 0.1 | - | - | - |
| *Cerastium holosteoides* | Wild | 0.1 | - | - | - |
| *Clinopodium vulgare* | Wild | 0.1 | - | - | - |
| *Crepis biennis* | Wild | 1.0 | - | - | - |
| *Cychorium intybus* | Wild | 2.5 | - | - | - |
| *Daucus carota* | Wild | 2.0 | - | - | 0.5 |
| *Dipsacus sylvestris/fullonur* | Wild | 0.1 | - | - | - |
| *Echium vulgare* | Wild | 1.3 | - | - | - |
| *Galium album* | Wild | 0.5 | - | - | - |
| *Galium verum* | Wild | 0.5 | - | - | - |
| *Heracleum sphondylium* | Wild | 0.4 | - | - | - |
| *Hypericum perforatum* | Wild | 0.1 | - | - | - |

| | | | | | |
|---|---|---|---|---|---|
| *Leucanthemum ircutianum* | Wild | 0.5 | - | - | 2.0 |
| *Lychnis flos-cuculi* | Wild | 0.2 | - | - | - |
| *Malva moschata* | Wild | 0.5 | - | - | - |
| *Malva sylvestris* | Wild | 1.0 | - | - | 1.5 |
| *Medicago lupulina* | Wild | 2.0 | - | - | - |
| *Origanum vulgare* | Wild | 0.2 | - | - | - |
| *Papaver rhoeas* | Wild | - | 0.5 | 0.5 | - |
| *Plantago lanceolata* | Wild | 0.5 | - | - | - |
| *Prunella vulgaris* | Wild | 0.1 | - | - | - |
| *Reseda lutea, luteola* | Wild | 0.2 | - | - | - |
| *Salvia pratensis* | Wild | 0.5 | - | - | - |
| *Sanguisorba minor* | Wild | 5.8 | - | - | - |
| *Silene alba* | Wild | 1.5 | - | - | - |
| *Silene dioica* | Wild | 0.5 | - | - | - |
| *Silene vulgaris* | Wild | 1.8 | - | - | - |
| *Tanacetum vulgare* | Wild | 0.3 | - | - | 0.5 |
| *Verbascum lychnitis, densif* | Wild | 0.3 | - | - | - |
