## Supplement B for "Criteria for effective fallow field eco schemes for farmland birds during non-breeding"

**Online Supplement B**. Bayesian model coefficient estimates with their confidence intervals and the sample of replicate fallow fields for each species and season. Bayes *P* indicates the posterior probability of the coefficient estimate to exceed zero, with values close to zero (or one) indicating particularly robust estimates for negative (or positive) predictor coefficients.

| | | | Coefficient estimate | | | | |
|---|---|---|---|---|---|---|---|
| **Species** | **Season** | **Predictor [level]** | **fit** | **lower CrI** | **upper CrI** | **n sites** | **Bayes *P*** |
| Blue Tit | Autumn | Intercept [Fallow type 1yr-FAKT] | -3.27 | -4.40 | -2.25 | 37 | 0.000 |
| Blue Tit | Autumn | Fallow type [1yr-GM] | 0.44 | -0.47 | 1.33 | 65 | 0.834 |
| Blue Tit | Autumn | Fallow type [2yr-GM] | 0.57 | -0.42 | 1.58 | 47 | 0.877 |
| Blue Tit | Autumn | Fallow type [3+yr] | 0.21 | -1.14 | 1.49 | 19 | 0.624 |
| Blue Tit | Autumn | Fallow field size (log-z) | 0.47 | 0.10 | 0.89 | | 0.994 |
| Blue Tit | Autumn | Hedge & grove cover (log-z) | 0.29 | -0.06 | 0.66 | | 0.950 |
| Blue Tit | Autumn | Fallow cover (log-z) | -0.16 | -0.55 | 0.24 | | 0.214 |
| Blue Tit | Autumn | Vegetation height (z) | 0.31 | -0.15 | 0.78 | | 0.909 |
| Blue Tit | Autumn | Farm track presence [yes] | -0.14 | -0.88 | 0.58 | | 0.344 |
| Blue Tit | Autumn | Secondary plant layer [yes] | 0.38 | -0.54 | 1.32 | | 0.788 |
| Brambling | Autumn | Intercept [Fallow type 1yr-FAKT] | -4.48 | -6.71 | -2.88 | 32 | 0.000 |
| Brambling | Autumn | Fallow type [1yr-GM] | 0.23 | -0.98 | 1.59 | 63 | 0.645 |
| Brambling | Autumn | Fallow type [2yr-GM] | -2.30 | -4.72 | -0.36 | 45 | 0.009 |
| Brambling | Autumn | Fallow type [3+yr] | -2.19 | -5.98 | 0.72 | 7 | 0.075 |
| Brambling | Autumn | Fallow field size (log-z) | 0.63 | 0.04 | 1.36 | | 0.982 |
| Brambling | Autumn | Hedge & grove cover (log-z) | 0.37 | -0.20 | 1.05 | | 0.901 |
| Brambling | Autumn | Fallow cover (log-z) | -0.51 | -1.23 | 0.09 | | 0.049 |
| Brambling | Autumn | Vegetation height (z) | -0.15 | -0.97 | 0.65 | | 0.347 |
| Brambling | Autumn | Farm track presence [yes] | -0.76 | -2.16 | 0.50 | | 0.118 |
| Brambling | Autumn | Secondary plant layer [yes] | 0.85 | -0.66 | 2.53 | | 0.864 |
| Chaffinch | Autumn | Intercept [Fallow type 1yr-FAKT] | -1.58 | -2.55 | -0.66 | 37 | 0.001 |
| Chaffinch | Autumn | Fallow type [1yr-GM] | -0.43 | -1.09 | 0.25 | 65 | 0.105 |
| Chaffinch | Autumn | Fallow type [2yr-GM] | -1.70 | -2.59 | -0.84 | 47 | 0.000 |
| Chaffinch | Autumn | Fallow type [3+yr] | -1.53 | -2.85 | -0.33 | 19 | 0.005 |
| Chaffinch | Autumn | Fallow field size (log-z) | 0.92 | 0.59 | 1.29 | | 1.000 |
| Chaffinch | Autumn | Hedge & grove cover (log-z) | 0.35 | 0.04 | 0.66 | | 0.986 |
| Chaffinch | Autumn | Fallow cover (log-z) | 0.06 | -0.26 | 0.39 | | 0.650 |
| Chaffinch | Autumn | Vegetation height (z) | 0.29 | -0.09 | 0.69 | | 0.934 |
| Chaffinch | Autumn | Farm track presence [yes] | -0.50 | -1.12 | 0.10 | | 0.050 |
| Chaffinch | Autumn | Secondary plant layer [yes] | 0.21 | -0.55 | 0.98 | | 0.711 |
| Goldfinch | Autumn | Intercept [Fallow type 1yr-FAKT] | -1.62 | -2.41 | -0.86 | 49 | 0.000 |
| Goldfinch | Autumn | Fallow type [1yr-GM] | -0.72 | -1.30 | -0.13 | 119 | 0.008 |
| Goldfinch | Autumn | Fallow type [2yr-GM] | -2.47 | -3.28 | -1.70 | 73 | 0.000 |
| Goldfinch | Autumn | Fallow type [3+yr] | -1.36 | -2.27 | -0.53 | 36 | 0.001 |
| Goldfinch | Autumn | Fallow field size (log-z) | 0.55 | 0.33 | 0.79 | | 1.000 |
| Goldfinch | Autumn | Hedge & grove cover (log-z) | -0.29 | -0.56 | -0.02 | | 0.018 |
| Goldfinch | Autumn | Fallow cover (log-z) | -0.11 | -0.35 | 0.13 | | 0.186 |
| Goldfinch | Autumn | Vegetation height (z) | 0.47 | 0.16 | 0.78 | | 0.999 |
| Goldfinch | Autumn | Study year [19-20] | -0.03 | -0.51 | 0.47 | | 0.460 |
| Goldfinch | Autumn | Farm track presence [yes] | 0.22 | -0.25 | 0.68 | | 0.824 |
| Goldfinch | Autumn | Secondary plant layer [yes] | 0.74 | 0.19 | 1.30 | | 0.995 |
| Great Tit | Autumn | Intercept [Fallow type 1yr-FAKT] | -2.40 | -3.41 | -1.42 | 37 | 0.000 |
| Great Tit | Autumn | Fallow type [1yr-GM] | 0.08 | -0.79 | 0.95 | 65 | 0.575 |
| Great Tit | Autumn | Fallow type [2yr-GM] | -1.14 | -2.23 | -0.12 | 47 | 0.015 |
| Great Tit | Autumn | Fallow type [3+yr] | -0.74 | -2.28 | 0.63 | 19 | 0.150 |
| Great Tit | Autumn | Fallow field size (log-z) | 0.42 | 0.05 | 0.83 | | 0.987 |
| Great Tit | Autumn | Hedge & grove cover (log-z) | 0.62 | 0.24 | 1.00 | | 0.999 |
| Great Tit | Autumn | Fallow cover (log-z) | -0.07 | -0.45 | 0.33 | | 0.355 |

| Species | Season | Predictor [level] | fit | lower CrI | upper CrI | n sites | Bayes *P* |
|---|---|---|---|---|---|---|---|
| Great Tit | Autumn | Vegetation height (z) | 0.19 | -0.30 | 0.69 | | 0.775 |
| Great Tit | Autumn | Farm track presence [yes] | -0.47 | -1.25 | 0.28 | | 0.109 |
| Great Tit | Autumn | Secondary plant layer [yes] | -0.19 | -1.16 | 0.78 | | 0.351 |
| Greenfinch | Autumn | Intercept [Fallow type 1yr-FAKT] | -2.18 | -3.13 | -1.33 | 49 | 0.000 |
| Greenfinch | Autumn | Fallow type [1yr-GM] | -0.19 | -0.81 | 0.43 | 118 | 0.271 |
| Greenfinch | Autumn | Fallow type [2yr-GM] | -1.33 | -2.15 | -0.53 | 71 | 0.001 |
| Greenfinch | Autumn | Fallow type [3+yr] | -2.53 | -4.25 | -1.17 | 25 | 0.000 |
| Greenfinch | Autumn | Fallow field size (log-z) | 0.32 | 0.06 | 0.60 | | 0.992 |
| Greenfinch | Autumn | Hedge & grove cover (log-z) | -0.15 | -0.45 | 0.14 | | 0.153 |
| Greenfinch | Autumn | Fallow cover (log-z) | -0.32 | -0.59 | -0.05 | | 0.010 |
| Greenfinch | Autumn | Vegetation height (z) | 0.13 | -0.20 | 0.48 | | 0.783 |
| Greenfinch | Autumn | Study year [19-20] | -0.42 | -0.97 | 0.15 | | 0.073 |
| Greenfinch | Autumn | Farm track presence [yes] | 0.14 | -0.37 | 0.64 | | 0.707 |
| Greenfinch | Autumn | Secondary plant layer [yes] | 0.43 | -0.17 | 1.06 | | 0.918 |
| Linnet | Autumn | Intercept [Fallow type 1yr-FAKT] | -4.51 | -5.87 | -3.39 | 46 | 0.000 |
| Linnet | Autumn | Fallow type [1yr-GM] | 0.40 | -0.39 | 1.23 | 114 | 0.840 |
| Linnet | Autumn | Fallow type [2yr-GM] | -0.84 | -2.02 | 0.26 | 67 | 0.068 |
| Linnet | Autumn | Fallow type [3+yr] | -0.06 | -1.47 | 1.22 | 13 | 0.466 |
| Linnet | Autumn | Fallow field size (log-z) | 0.49 | 0.18 | 0.84 | | 0.999 |
| Linnet | Autumn | Hedge & grove cover (log-z) | -0.42 | -0.81 | -0.05 | | 0.012 |
| Linnet | Autumn | Fallow cover (log-z) | -0.09 | -0.43 | 0.23 | | 0.293 |
| Linnet | Autumn | Vegetation height (z) | 0.18 | -0.26 | 0.63 | | 0.790 |
| Linnet | Autumn | Study year [19-20] | 0.82 | 0.12 | 1.59 | | 0.989 |
| Linnet | Autumn | Farm track presence [yes] | 0.31 | -0.35 | 0.95 | | 0.828 |
| Linnet | Autumn | Secondary plant layer [yes] | 0.50 | -0.30 | 1.31 | | 0.893 |
| Reed Bunting | Autumn | Intercept [Fallow type 1yr-FAKT] | -5.18 | -6.73 | -3.84 | 49 | 0.000 |
| Reed Bunting | Autumn | Fallow type [1yr-GM] | 0.52 | -0.61 | 1.74 | 118 | 0.814 |
| Reed Bunting | Autumn | Fallow type [2yr-GM] | 2.11 | 1.00 | 3.35 | 73 | 1.000 |
| Reed Bunting | Autumn | Fallow type [3+yr] | 1.68 | 0.48 | 3.04 | 35 | 0.997 |
| Reed Bunting | Autumn | Fallow field size (log-z) | 0.42 | 0.09 | 0.77 | | 0.994 |
| Reed Bunting | Autumn | Hedge & grove cover (log-z) | -0.39 | -0.82 | 0.00 | | 0.026 |
| Reed Bunting | Autumn | Fallow cover (log-z) | 0.11 | -0.26 | 0.48 | | 0.722 |
| Reed Bunting | Autumn | Vegetation height (z) | 0.45 | 0.02 | 0.88 | | 0.980 |
| Reed Bunting | Autumn | Study year [19-20] | 0.54 | -0.10 | 1.20 | | 0.950 |
| Reed Bunting | Autumn | Farm track presence [yes] | -0.17 | -0.90 | 0.50 | | 0.311 |
| Reed Bunting | Autumn | Secondary plant layer [yes] | 0.28 | -0.57 | 1.11 | | 0.749 |
| Skylark | Autumn | Intercept [Fallow type 1yr-FAKT] | -4.33 | -6.33 | -2.68 | 31 | 0.000 |
| Skylark | Autumn | Fallow type [1yr-GM] | 0.91 | -0.52 | 2.45 | 87 | 0.893 |
| Skylark | Autumn | Fallow type [2yr-GM] | 0.73 | -0.91 | 2.40 | 51 | 0.808 |
| Skylark | Autumn | Fallow type [3+yr] | 0.49 | -1.24 | 2.26 | 33 | 0.714 |
| Skylark | Autumn | Fallow field size (log-z) | 0.69 | 0.23 | 1.18 | | 0.999 |
| Skylark | Autumn | Hedge & grove cover (log-z) | -0.93 | -1.81 | -0.21 | | 0.005 |
| Skylark | Autumn | Fallow cover (log-z) | -0.28 | -0.77 | 0.18 | | 0.115 |
| Skylark | Autumn | Vegetation height (z) | -0.55 | -1.22 | 0.11 | | 0.050 |
| Skylark | Autumn | Study year [19-20] | -1.77 | -2.93 | -0.70 | | 0.001 |
| Skylark | Autumn | Farm track presence [yes] | -0.26 | -1.31 | 0.74 | | 0.300 |
| Skylark | Autumn | Secondary plant layer [yes] | -0.53 | -1.60 | 0.51 | | 0.152 |
| Stonechat | Autumn | Intercept [Fallow type 1yr-FAKT] | -5.03 | -6.97 | -3.44 | 40 | 0.000 |
| Stonechat | Autumn | Fallow type [1yr-GM] | 0.30 | -1.14 | 1.86 | 99 | 0.651 |
| Stonechat | Autumn | Fallow type [2yr-GM] | 0.73 | -0.86 | 2.37 | 57 | 0.818 |
| Stonechat | Autumn | Fallow type [3+yr] | 0.94 | -0.77 | 2.78 | 22 | 0.858 |
| Stonechat | Autumn | Fallow field size (log-z) | 0.38 | -0.14 | 0.92 | | 0.925 |
| Stonechat | Autumn | Hedge & grove cover (log-z) | -0.72 | -1.41 | -0.11 | | 0.010 |
| Stonechat | Autumn | Fallow cover (log-z) | 0.36 | -0.15 | 0.91 | | 0.918 |

| Species | Season | Predictor [level] | fit | lower CrI | upper CrI | n sites | Bayes *P* |
|---|---|---|---|---|---|---|---|
| Stonechat | Autumn | Vegetation height (z) | 0.26 | -0.37 | 0.94 | | 0.790 |
| Stonechat | Autumn | Study year [19-20] | -0.44 | -1.48 | 0.55 | | 0.193 |
| Stonechat | Autumn | Farm track presence [yes] | -0.19 | -1.23 | 0.71 | | 0.346 |
| Stonechat | Autumn | Secondary plant layer [yes] | 0.30 | -0.90 | 1.55 | | 0.695 |
| Tree Sparrow | Autumn | Intercept [Fallow type 1yr-FAKT] | -1.82 | -2.90 | -0.78 | 49 | 0.000 |
| Tree Sparrow | Autumn | Fallow type [1yr-GM] | -0.33 | -1.07 | 0.43 | 118 | 0.197 |
| Tree Sparrow | Autumn | Fallow type [2yr-GM] | -2.12 | -3.09 | -1.20 | 73 | 0.000 |
| Tree Sparrow | Autumn | Fallow type [3+yr] | -2.08 | -3.29 | -0.91 | 35 | 0.000 |
| Tree Sparrow | Autumn | Fallow field size (log-z) | 0.49 | 0.19 | 0.81 | | 0.999 |
| Tree Sparrow | Autumn | Hedge & grove cover (log-z) | 0.12 | -0.21 | 0.45 | | 0.760 |
| Tree Sparrow | Autumn | Fallow cover (log-z) | 0.06 | -0.27 | 0.38 | | 0.645 |
| Tree Sparrow | Autumn | Vegetation height (z) | 0.72 | 0.33 | 1.12 | | 1.000 |
| Tree Sparrow | Autumn | Study year [19-20] | -0.15 | -0.79 | 0.50 | | 0.320 |
| Tree Sparrow | Autumn | Farm track presence [yes] | -0.34 | -0.98 | 0.25 | | 0.130 |
| Tree Sparrow | Autumn | Secondary plant layer [yes] | 0.57 | -0.13 | 1.29 | | 0.943 |
| Whinchat | Autumn | Intercept [Fallow type 1yr-FAKT] | -5.70 | -7.53 | -4.12 | 26 | 0.000 |
| Whinchat | Autumn | Fallow type [1yr-GM] | 1.12 | -0.24 | 2.66 | 81 | 0.946 |
| Whinchat | Autumn | Fallow type [2yr-GM] | 1.48 | 0.04 | 3.09 | 49 | 0.978 |
| Whinchat | Autumn | Fallow type [3+yr] | 2.51 | 1.05 | 4.11 | 28 | 0.999 |
| Whinchat | Autumn | Fallow field size (log-z) | 0.39 | 0.01 | 0.77 | | 0.979 |
| Whinchat | Autumn | Hedge & grove cover (log-z) | -0.51 | -1.23 | 0.11 | | 0.058 |
| Whinchat | Autumn | Fallow cover (log-z) | -0.11 | -0.50 | 0.28 | | 0.292 |
| Whinchat | Autumn | Vegetation height (z) | 0.52 | -0.01 | 1.07 | | 0.972 |
| Whinchat | Autumn | Study year [19-20] | -1.12 | -2.17 | -0.19 | | 0.008 |
| Whinchat | Autumn | Farm track presence [yes] | 0.74 | -0.01 | 1.49 | | 0.973 |
| Whinchat | Autumn | Secondary plant layer [yes] | 0.64 | -0.29 | 1.62 | | 0.912 |
| Whitethroat | Autumn | Intercept [Fallow type 1yr-FAKT] | -3.82 | -5.89 | -2.13 | 12 | 0.000 |
| Whitethroat | Autumn | Fallow type [1yr-GM] | 0.07 | -1.58 | 1.84 | 48 | 0.530 |
| Whitethroat | Autumn | Fallow type [2yr-GM] | 1.81 | -0.19 | 3.92 | 22 | 0.962 |
| Whitethroat | Autumn | Fallow type [3+yr] | -1.11 | -3.88 | 1.25 | 16 | 0.181 |
| Whitethroat | Autumn | Fallow field size (log-z) | -0.19 | -0.81 | 0.42 | | 0.267 |
| Whitethroat | Autumn | Hedge & grove cover (log-z) | 0.99 | 0.35 | 1.79 | | 0.999 |
| Whitethroat | Autumn | Fallow cover (log-z) | 0.32 | -0.28 | 0.94 | | 0.863 |
| Whitethroat | Autumn | Vegetation height (z) | -0.49 | -1.26 | 0.31 | | 0.107 |
| Whitethroat | Autumn | Farm track presence [yes] | 0.71 | -0.35 | 1.80 | | 0.907 |
| Whitethroat | Autumn | Secondary plant layer [yes] | 0.27 | -0.93 | 1.57 | | 0.674 |
| Yellowhammer | Autumn | Intercept [Fallow type 1yr-FAKT] | -1.92 | -2.72 | -1.16 | 49 | 0.000 |
| Yellowhammer | Autumn | Fallow type [1yr-GM] | 0.39 | -0.25 | 1.07 | 118 | 0.881 |
| Yellowhammer | Autumn | Fallow type [2yr-GM] | 0.85 | 0.14 | 1.58 | 73 | 0.991 |
| Yellowhammer | Autumn | Fallow type [3+yr] | 0.83 | 0.02 | 1.66 | 35 | 0.977 |
| Yellowhammer | Autumn | Fallow field size (log-z) | 0.16 | -0.05 | 0.39 | | 0.932 |
| Yellowhammer | Autumn | Hedge & grove cover (log-z) | -0.09 | -0.33 | 0.16 | | 0.241 |
| Yellowhammer | Autumn | Fallow cover (log-z) | -0.10 | -0.35 | 0.14 | | 0.198 |
| Yellowhammer | Autumn | Vegetation height (z) | 0.09 | -0.19 | 0.37 | | 0.743 |
| Yellowhammer | Autumn | Study year [19-20] | -0.94 | -1.40 | -0.49 | | 0.000 |
| Yellowhammer | Autumn | Farm track presence [yes] | -0.23 | -0.67 | 0.21 | | 0.153 |
| Yellowhammer | Autumn | Secondary plant layer [yes] | 0.16 | -0.35 | 0.67 | | 0.738 |

| | | | Coefficient estimate | | | | |
|---|---|---|---|---|---|---|---|
| Species | Season | Predictor [level] | fit | lower CrI | upper CrI | | Bayes *P* |
| Blue Tit | Winter | Intercept [Fallow type 1yr-FAKT] | -4.68 | -7.04 | -2.81 | 30 | 0.000 |
| Blue Tit | Winter | Fallow type [1yr-GM] | -0.59 | -1.72 | 0.60 | 97 | 0.154 |
| Blue Tit | Winter | Fallow type [2yr-GM] | -0.26 | -1.64 | 1.14 | 42 | 0.357 |
| Blue Tit | Winter | Fallow type [3+yr] | -1.20 | -3.68 | 0.75 | 11 | 0.116 |

| Species | Season | Predictor [level] | fit | lower CrI | upper CrI | n sites | Bayes *P* |
|---|---|---|---|---|---|---|---|
| Blue Tit | Winter | Fallow field size (log-z) | 0.45 | -0.02 | 0.98 | | 0.970 |
| Blue Tit | Winter | Hedge & grove cover (log-z) | 0.39 | -0.06 | 0.87 | | 0.954 |
| Blue Tit | Winter | Fallow cover (log-z) | 0.09 | -0.40 | 0.59 | | 0.641 |
| Blue Tit | Winter | Vegetation height (z) | 0.39 | -0.17 | 0.98 | | 0.915 |
| Blue Tit | Winter | Study year [19-20] | 1.30 | 0.04 | 2.71 | | 0.979 |
| Blue Tit | Winter | Farm track presence [yes] | -0.08 | -1.02 | 0.83 | | 0.430 |
| Blue Tit | Winter | Secondary plant layer [yes] | 0.41 | -0.78 | 1.67 | | 0.749 |
| Chaffinch | Winter | Intercept [Fallow type 1yr-FAKT] | -4.14 | -6.03 | -2.47 | 37 | 0.000 |
| Chaffinch | Winter | Fallow type [1yr-GM] | -0.52 | -1.62 | 0.55 | 106 | 0.168 |
| Chaffinch | Winter | Fallow type [2yr-GM] | -1.02 | -2.66 | 0.46 | 47 | 0.090 |
| Chaffinch | Winter | Fallow type [3+yr] | -0.94 | -2.53 | 0.51 | 26 | 0.104 |
| Chaffinch | Winter | Fallow field size (log-z) | 0.85 | 0.45 | 1.30 | | 1.000 |
| Chaffinch | Winter | Hedge & grove cover (log-z) | 0.08 | -0.36 | 0.54 | | 0.645 |
| Chaffinch | Winter | Fallow cover (log-z) | -0.12 | -0.56 | 0.33 | | 0.288 |
| Chaffinch | Winter | Vegetation height (z) | 0.70 | 0.16 | 1.28 | | 0.995 |
| Chaffinch | Winter | Study year [19-20] | -0.21 | -1.12 | 0.67 | | 0.319 |
| Chaffinch | Winter | Farm track presence [yes] | 0.40 | -0.43 | 1.28 | | 0.831 |
| Chaffinch | Winter | Secondary plant layer [yes] | 1.36 | 0.22 | 2.61 | | 0.990 |
| Goldfinch | Winter | Intercept [Fallow type 1yr-FAKT] | -4.06 | -5.89 | -2.46 | 37 | 0.000 |
| Goldfinch | Winter | Fallow type [1yr-GM] | -0.75 | -1.81 | 0.32 | 107 | 0.082 |
| Goldfinch | Winter | Fallow type [2yr-GM] | 0.19 | -1.14 | 1.56 | 49 | 0.610 |
| Goldfinch | Winter | Fallow type [3+yr] | -0.80 | -2.15 | 0.56 | 36 | 0.124 |
| Goldfinch | Winter | Fallow field size (log-z) | 1.04 | 0.64 | 1.53 | | 1.000 |
| Goldfinch | Winter | Hedge & grove cover (log-z) | -0.46 | -0.92 | -0.04 | | 0.016 |
| Goldfinch | Winter | Fallow cover (log-z) | -0.08 | -0.52 | 0.37 | | 0.359 |
| Goldfinch | Winter | Vegetation height (z) | 0.38 | -0.13 | 0.92 | | 0.928 |
| Goldfinch | Winter | Study year [19-20] | -0.46 | -1.27 | 0.34 | | 0.122 |
| Goldfinch | Winter | Farm track presence [yes] | 0.55 | -0.29 | 1.36 | | 0.906 |
| Goldfinch | Winter | Secondary plant layer [yes] | 1.34 | 0.28 | 2.53 | | 0.993 |
| Great Tit | Winter | Intercept [Fallow type 1yr-FAKT] | -5.79 | -8.77 | -3.36 | 30 | 0.000 |
| Great Tit | Winter | Fallow type [1yr-GM] | -1.22 | -2.89 | 0.34 | 82 | 0.062 |
| Great Tit | Winter | Fallow type [2yr-GM] | -1.42 | -3.48 | 0.47 | 38 | 0.072 |
| Great Tit | Winter | Fallow type [3+yr] | 0.31 | -2.03 | 2.62 | 11 | 0.607 |
| Great Tit | Winter | Fallow field size (log-z) | 0.78 | 0.05 | 1.60 | | 0.983 |
| Great Tit | Winter | Hedge & grove cover (log-z) | 0.97 | 0.28 | 1.83 | | 0.997 |
| Great Tit | Winter | Fallow cover (log-z) | 0.90 | 0.20 | 1.79 | | 0.995 |
| Great Tit | Winter | Vegetation height (z) | 0.78 | -0.04 | 1.66 | | 0.969 |
| Great Tit | Winter | Study year [19-20] | 1.52 | -0.16 | 3.29 | | 0.962 |
| Great Tit | Winter | Farm track presence [yes] | -1.32 | -3.01 | 0.11 | | 0.036 |
| Great Tit | Winter | Secondary plant layer [yes] | 1.01 | -0.67 | 2.82 | | 0.876 |
| Greenfinch | Winter | Intercept [Fallow type 1yr-FAKT] | -6.31 | -9.42 | -4.02 | 28 | 0.000 |
| Greenfinch | Winter | Fallow type [1yr-GM] | -0.45 | -1.88 | 1.04 | 68 | 0.267 |
| Greenfinch | Winter | Fallow type [2yr-GM] | -0.64 | -2.92 | 1.46 | 34 | 0.276 |
| Greenfinch | Winter | Fallow type [3+yr] | -1.13 | -3.54 | 1.04 | 12 | 0.151 |
| Greenfinch | Winter | Fallow field size (log-z) | 0.57 | -0.02 | 1.28 | | 0.971 |
| Greenfinch | Winter | Hedge & grove cover (log-z) | -0.23 | -1.00 | 0.48 | | 0.253 |
| Greenfinch | Winter | Fallow cover (log-z) | -0.16 | -0.95 | 0.67 | | 0.343 |
| Greenfinch | Winter | Vegetation height (z) | 0.05 | -0.86 | 0.88 | | 0.550 |
| Greenfinch | Winter | Study year [19-20] | 1.29 | -0.08 | 2.78 | | 0.968 |
| Greenfinch | Winter | Farm track presence [yes] | 0.20 | -1.19 | 1.49 | | 0.624 |
| Greenfinch | Winter | Secondary plant layer [yes] | 1.96 | 0.32 | 3.94 | | 0.990 |
| Linnet | Winter | Intercept [Fallow type 1yr-FAKT] | -8.00 | -12.54 | -4.50 | 21 | 0.000 |
| Linnet | Winter | Fallow type [1yr-GM] | 1.80 | -0.52 | 4.35 | 63 | 0.936 |
| Linnet | Winter | Fallow type [2yr-GM] | -1.22 | -5.44 | 2.51 | 15 | 0.268 |

| Species | Season | Predictor [level] | fit | lower CrI | upper CrI | n sites | Bayes *P* |
|---|---|---|---|---|---|---|---|
| Linnet | Winter | Fallow type [3+yr] | 0.88 | -2.12 | 3.81 | 9 | 0.724 |
| Linnet | Winter | Fallow field size (log-z) | 1.10 | 0.28 | 2.11 | | 0.996 |
| Linnet | Winter | Hedge & grove cover (log-z) | -1.00 | -2.35 | 0.10 | | 0.039 |
| Linnet | Winter | Fallow cover (log-z) | 0.11 | -0.98 | 1.25 | | 0.580 |
| Linnet | Winter | Vegetation height (z) | -0.41 | -1.91 | 0.97 | | 0.279 |
| Linnet | Winter | Study year [19-20] | 0.16 | -2.00 | 2.36 | | 0.557 |
| Linnet | Winter | Farm track presence [yes] | 0.01 | -1.96 | 1.99 | | 0.503 |
| Linnet | Winter | Secondary plant layer [yes] | 1.21 | -1.18 | 3.90 | | 0.843 |
| Reed Bunting | Winter | Intercept [Fallow type 1yr-FAKT] | -3.88 | -6.52 | -1.54 | 29 | 0.001 |
| Reed Bunting | Winter | Fallow type [1yr-GM] | -1.15 | -2.86 | 0.54 | 92 | 0.093 |
| Reed Bunting | Winter | Fallow type [2yr-GM] | -0.27 | -2.12 | 1.61 | 46 | 0.386 |
| Reed Bunting | Winter | Fallow type [3+yr] | -0.70 | -2.89 | 1.40 | 30 | 0.255 |
| Reed Bunting | Winter | Fallow field size (log-z) | 0.37 | -0.19 | 1.00 | | 0.905 |
| Reed Bunting | Winter | Hedge & grove cover (log-z) | -0.11 | -0.73 | 0.49 | | 0.354 |
| Reed Bunting | Winter | Fallow cover (log-z) | 0.19 | -0.45 | 0.88 | | 0.726 |
| Reed Bunting | Winter | Vegetation height (z) | 0.64 | -0.09 | 1.45 | | 0.958 |
| Reed Bunting | Winter | Study year [19-20] | -1.04 | -2.41 | 0.23 | | 0.052 |
| Reed Bunting | Winter | Farm track presence [yes] | 1.35 | 0.21 | 2.62 | | 0.990 |
| Reed Bunting | Winter | Secondary plant layer [yes] | 0.63 | -0.92 | 2.30 | | 0.788 |
| Tree Sparrow | Winter | Intercept [Fallow type 1yr-FAKT] | -6.21 | -9.41 | -3.57 | 27 | 0.000 |
| Tree Sparrow | Winter | Fallow type [1yr-GM] | 0.24 | -1.27 | 1.84 | 50 | 0.626 |
| Tree Sparrow | Winter | Fallow type [2yr-GM] | 0.11 | -2.49 | 2.57 | 11 | 0.534 |
| Tree Sparrow | Winter | Fallow type [3+yr] | 0.34 | -1.68 | 2.26 | 13 | 0.636 |
| Tree Sparrow | Winter | Fallow field size (log-z) | 0.67 | 0.13 | 1.36 | | 0.992 |
| Tree Sparrow | Winter | Hedge & grove cover (log-z) | -0.08 | -0.91 | 0.72 | | 0.418 |
| Tree Sparrow | Winter | Fallow cover (log-z) | -0.19 | -0.89 | 0.56 | | 0.301 |
| Tree Sparrow | Winter | Vegetation height (z) | 0.33 | -0.59 | 1.24 | | 0.767 |
| Tree Sparrow | Winter | Study year [19-20] | 0.36 | -1.26 | 1.90 | | 0.679 |
| Tree Sparrow | Winter | Farm track presence [yes] | 1.02 | -0.40 | 2.55 | | 0.923 |
| Tree Sparrow | Winter | Secondary plant layer [yes] | 2.00 | 0.20 | 3.96 | | 0.985 |
| Yellowhammer | Winter | Intercept [Fallow type 1yr-FAKT] | -4.72 | -6.29 | -3.30 | 37 | 0.000 |
| Yellowhammer | Winter | Fallow type [1yr-GM] | 0.95 | -0.07 | 2.06 | 107 | 0.965 |
| Yellowhammer | Winter | Fallow type [2yr-GM] | 1.40 | 0.20 | 2.66 | 49 | 0.990 |
| Yellowhammer | Winter | Fallow type [3+yr] | 1.57 | 0.35 | 2.83 | 36 | 0.995 |
| Yellowhammer | Winter | Fallow field size (log-z) | 0.64 | 0.32 | 0.99 | | 1.000 |
| Yellowhammer | Winter | Hedge & grove cover (log-z) | -0.20 | -0.55 | 0.16 | | 0.135 |
| Yellowhammer | Winter | Fallow cover (log-z) | -0.16 | -0.52 | 0.21 | | 0.193 |
| Yellowhammer | Winter | Vegetation height (z) | 0.33 | -0.09 | 0.77 | | 0.940 |
| Yellowhammer | Winter | Study year [19-20] | 0.54 | -0.16 | 1.28 | | 0.936 |
| Yellowhammer | Winter | Farm track presence [yes] | 0.30 | -0.41 | 0.99 | | 0.803 |
| Yellowhammer | Winter | Secondary plant layer [yes] | 0.19 | -0.67 | 1.09 | | 0.671 |

| | | | Coefficient estimate | | | | |
|---|---|---|---|---|---|---|---|
| Response | Season | Predictor [level] | fit | lower CrI | upper CrI | | Bayes *P* |
| No. Species | Winter | Intercept [Fallow type 1yr-FAKT] | -0.29 | -1.06 | 0.43 | 49 | 0.214 |
| No. Species | Winter | Fallow type [1yr-GM] | 0.13 | -0.42 | 0.66 | 119 | 0.676 |
| No. Species | Winter | Fallow type [2yr-GM] | -0.04 | -0.61 | 0.51 | 73 | 0.438 |
| No. Species | Winter | Fallow type [3+yr] | -0.08 | -0.71 | 0.55 | 36 | 0.404 |
| No. Species | Winter | Fallow field size (log-z) | 0.44 | 0.28 | 0.63 | | 1.000 |
| No. Species | Winter | Hedge & grove cover (log-z) | -0.07 | -0.23 | 0.10 | | 0.221 |
| No. Species | Winter | Fallow cover (log-z) | 0.14 | -0.04 | 0.32 | | 0.939 |
| No. Species | Winter | Vegetation height (z) | 0.25 | 0.05 | 0.46 | | 0.992 |
| No. Species | Winter | Complete survey [yes] | 0.22 | -0.29 | 0.75 | | 0.805 |
| No. Species | Winter | Study year [19-20] | -0.23 | -0.58 | 0.11 | | 0.099 |

| Species | Season | Predictor [level] | fit | lower CrI | upper CrI | n sites | Bayes *P* |
|---|---|---|---|---|---|---|---|
| No. Species | Winter | Farm track presence [yes] | 0.19 | -0.14 | 0.53 | | 0.873 |
| No. Species | Winter | Secondary plant layer [yes] | 0.61 | 0.16 | 1.05 | | 0.997 |
| No. Species | Autumn | Intercept [Fallow type 1yr-FAKT] | 1.13 | 0.86 | 1.41 | 37 | 1.000 |
| No. Species | Autumn | Fallow type [1yr-GM] | 0.08 | -0.14 | 0.29 | 107 | 0.760 |
| No. Species | Autumn | Fallow type [2yr-GM] | -0.18 | -0.43 | 0.07 | 49 | 0.084 |
| No. Species | Autumn | Fallow type [3+yr] | -0.24 | -0.56 | 0.06 | 36 | 0.060 |
| No. Species | Autumn | Fallow field size (log-z) | 0.21 | 0.13 | 0.30 | | 1.000 |
| No. Species | Autumn | Hedge & grove cover (log-z) | -0.01 | -0.10 | 0.08 | | 0.404 |
| No. Species | Autumn | Fallow cover (log-z) | -0.04 | -0.13 | 0.05 | | 0.185 |
| No. Species | Autumn | Vegetation height (z) | 0.17 | 0.06 | 0.27 | | 0.999 |
| No. Species | Autumn | Study year [19-20] | 0.17 | -0.01 | 0.34 | | 0.970 |
| No. Species | Autumn | Farm track presence [yes] | -0.09 | -0.26 | 0.07 | | 0.140 |
| No. Species | Autumn | Secondary plant layer [yes] | 0.20 | 0.01 | 0.40 | | 0.980 |
