## Supplement C for "Criteria for effective fallow field eco schemes for farmland birds during non-breeding"

**Online Supplement C**. Pairwise comparisons of species numbers and bird incidences between fallow field types, showing median Bayesian posterior differences between the assigned Base and Reference levels. Moreover, we display the posterior probabilities ('Bayes P') that bird incidence was higher on the Base compared to the Reference level. The explanatory power of fallow field types increases when these posterior probabilities approach 0 or 1. There is no objective cut-off value, but in order to make general patterns within and between species more transparent, we highlighted oarticularly robust pairwise comparisons with Bayes P < 0.02 or > 0.98 in bold blue face.

| | | Fallow field types | | Δ bird incidence (Base-Reference) | |
|---|---|---|---|---|---|
| Species | Season | Base | Reference | Median | Bayes *P* |
| Blue Tit | Autumn | 1yr-FAKT | 1yr-GM | -0.025 | 0.166 |
| Blue Tit | Autumn | 1yr-FAKT | 2yr-GM | -0.035 | 0.123 |
| Blue Tit | Autumn | 1yr-FAKT | 3+yr | -0.010 | 0.376 |
| Blue Tit | Autumn | 1yr-GM | 2yr-GM | -0.010 | 0.358 |
| Blue Tit | Autumn | 1yr-GM | 3+yr | 0.014 | 0.644 |
| Blue Tit | Autumn | 2yr-GM | 3+yr | 0.024 | 0.712 |
| Brambling | Autumn | 1yr-FAKT | 1yr-GM | -0.005 | 0.355 |
| Brambling | Autumn | 1yr-FAKT | 2yr-GM | **0.022** | **0.991** |
| Brambling | Autumn | 1yr-FAKT | 3+yr | 0.020 | 0.925 |
| Brambling | Autumn | 1yr-GM | 2yr-GM | **0.029** | **0.998** |
| Brambling | Autumn | 1yr-GM | 3+yr | 0.027 | 0.950 |
| Brambling | Autumn | 2yr-GM | 3+yr | 0.000 | 0.477 |
| Chaffinch | Autumn | 1yr-FAKT | 1yr-GM | 0.057 | 0.895 |
| Chaffinch | Autumn | 1yr-FAKT | 2yr-GM | **0.154** | **1.000** |
| Chaffinch | Autumn | 1yr-FAKT | 3+yr | **0.142** | **0.995** |
| Chaffinch | Autumn | 1yr-GM | 2yr-GM | **0.095** | **1.000** |
| Chaffinch | Autumn | 1yr-GM | 3+yr | 0.084 | 0.970 |
| Chaffinch | Autumn | 2yr-GM | 3+yr | -0.007 | 0.402 |
| Goldfinch | Autumn | 1yr-FAKT | 1yr-GM | **0.124** | **0.992** |
| Goldfinch | Autumn | 1yr-FAKT | 2yr-GM | **0.260** | **1.000** |
| Goldfinch | Autumn | 1yr-FAKT | 3+yr | **0.194** | **0.999** |
| Goldfinch | Autumn | 1yr-GM | 2yr-GM | **0.135** | **1.000** |
| Goldfinch | Autumn | 1yr-GM | 3+yr | 0.072 | 0.950 |
| Goldfinch | Autumn | 2yr-GM | 3+yr | **-0.061** | **0.011** |
| Great Tit | Autumn | 1yr-FAKT | 1yr-GM | -0.006 | 0.425 |
| Great Tit | Autumn | 1yr-FAKT | 2yr-GM | **0.050** | **0.985** |
| Great Tit | Autumn | 1yr-FAKT | 3+yr | 0.036 | 0.850 |
| Great Tit | Autumn | 1yr-GM | 2yr-GM | **0.057** | **0.998** |
| Great Tit | Autumn | 1yr-GM | 3+yr | 0.042 | 0.881 |
| Great Tit | Autumn | 2yr-GM | 3+yr | -0.012 | 0.305 |
| Greenfinch | Autumn | 1yr-FAKT | 1yr-GM | 0.016 | 0.729 |
| Greenfinch | Autumn | 1yr-FAKT | 2yr-GM | **0.073** | **0.999** |
| Greenfinch | Autumn | 1yr-FAKT | 3+yr | **0.093** | **1.000** |
| Greenfinch | Autumn | 1yr-GM | 2yr-GM | **0.057** | **1.000** |
| Greenfinch | Autumn | 1yr-GM | 3+yr | **0.077** | **1.000** |
| Greenfinch | Autumn | 2yr-GM | 3+yr | 0.019 | 0.942 |
| Linnet | Autumn | 1yr-FAKT | 1yr-GM | -0.018 | 0.160 |
| Linnet | Autumn | 1yr-FAKT | 2yr-GM | 0.022 | 0.932 |
| Linnet | Autumn | 1yr-FAKT | 3+yr | 0.002 | 0.534 |

| | | | | | |
|---|---|---|---|---|---|
| Linnet | Autumn | 1yr-GM | 2yr-GM | **0.040** | **0.997** |
| Linnet | Autumn | 1yr-GM | 3+yr | 0.020 | 0.775 |
| Linnet | Autumn | 2yr-GM | 3+yr | -0.019 | 0.157 |
| Reed Bunting | Autumn | 1yr-FAKT | 1yr-GM | -0.008 | 0.186 |
| Reed Bunting | Autumn | 1yr-FAKT | 2yr-GM | **-0.085** | **0.000** |
| Reed Bunting | Autumn | 1yr-FAKT | 3+yr | **-0.053** | **0.003** |
| Reed Bunting | Autumn | 1yr-GM | 2yr-GM | **-0.076** | **0.000** |
| Reed Bunting | Autumn | 1yr-GM | 3+yr | **-0.044** | **0.014** |
| Reed Bunting | Autumn | 2yr-GM | 3+yr | 0.030 | 0.802 |
| Skylark | Autumn | 1yr-FAKT | 1yr-GM | -0.002 | 0.107 |
| Skylark | Autumn | 1yr-FAKT | 2yr-GM | -0.001 | 0.192 |
| Skylark | Autumn | 1yr-FAKT | 3+yr | -0.001 | 0.286 |
| Skylark | Autumn | 1yr-GM | 2yr-GM | 0.001 | 0.604 |
| Skylark | Autumn | 1yr-GM | 3+yr | 0.001 | 0.753 |
| Skylark | Autumn | 2yr-GM | 3+yr | 0.001 | 0.622 |
| Stonechat | Autumn | 1yr-FAKT | 1yr-GM | -0.002 | 0.349 |
| Stonechat | Autumn | 1yr-FAKT | 2yr-GM | -0.006 | 0.182 |
| Stonechat | Autumn | 1yr-FAKT | 3+yr | -0.008 | 0.142 |
| Stonechat | Autumn | 1yr-GM | 2yr-GM | -0.004 | 0.235 |
| Stonechat | Autumn | 1yr-GM | 3+yr | -0.007 | 0.183 |
| Stonechat | Autumn | 2yr-GM | 3+yr | -0.003 | 0.392 |
| Tree Sparrow | Autumn | 1yr-FAKT | 1yr-GM | 0.046 | 0.803 |
| Tree Sparrow | Autumn | 1yr-FAKT | 2yr-GM | **0.171** | **1.000** |
| Tree Sparrow | Autumn | 1yr-FAKT | 3+yr | **0.167** | **1.000** |
| Tree Sparrow | Autumn | 1yr-GM | 2yr-GM | **0.123** | **1.000** |
| Tree Sparrow | Autumn | 1yr-GM | 3+yr | **0.120** | **0.999** |
| Tree Sparrow | Autumn | 2yr-GM | 3+yr | -0.001 | 0.474 |
| Whinchat | Autumn | 1yr-FAKT | 1yr-GM | -0.005 | 0.054 |
| Whinchat | Autumn | 1yr-FAKT | 2yr-GM | -0.008 | 0.022 |
| Whinchat | Autumn | 1yr-FAKT | 3+yr | **-0.029** | **0.001** |
| Whinchat | Autumn | 1yr-GM | 2yr-GM | -0.003 | 0.253 |
| Whinchat | Autumn | 1yr-GM | 3+yr | **-0.024** | **0.004** |
| Whinchat | Autumn | 2yr-GM | 3+yr | -0.020 | 0.035 |
| Whitethroat | Autumn | 1yr-FAKT | 1yr-GM | -0.001 | 0.470 |
| Whitethroat | Autumn | 1yr-FAKT | 2yr-GM | -0.056 | 0.038 |
| Whitethroat | Autumn | 1yr-FAKT | 3+yr | 0.007 | 0.819 |
| Whitethroat | Autumn | 1yr-GM | 2yr-GM | **-0.057** | **0.016** |
| Whitethroat | Autumn | 1yr-GM | 3+yr | 0.008 | 0.870 |
| Whitethroat | Autumn | 2yr-GM | 3+yr | **0.067** | **0.995** |
| Yellowhammer | Autumn | 1yr-FAKT | 1yr-GM | -0.027 | 0.119 |
| Yellowhammer | Autumn | 1yr-FAKT | 2yr-GM | **-0.071** | **0.009** |
| Yellowhammer | Autumn | 1yr-FAKT | 3+yr | -0.070 | 0.023 |
| Yellowhammer | Autumn | 1yr-GM | 2yr-GM | -0.044 | 0.043 |
| Yellowhammer | Autumn | 1yr-GM | 3+yr | -0.043 | 0.107 |
| Yellowhammer | Autumn | 2yr-GM | 3+yr | 0.001 | 0.511 |

| | | Fallow field types | | Δ bird incidence (Base-Reference) | |
|---|---|---|---|---|---|
| **Species** | **Season** | **Base** | **Reference** | **Median** | **Bayes *P*** |
| Blue Tit | Winter | 1yr-FAKT | 1yr-GM | 0.021 | 0.846 |
| Blue Tit | Winter | 1yr-FAKT | 2yr-GM | 0.010 | 0.643 |

| | | | | | |
|---|---|---|---|---|---|
| Blue Tit | Winter | 1yr-FAKT | 3+yr | 0.032 | 0.884 |
| Blue Tit | Winter | 1yr-GM | 2yr-GM | -0.011 | 0.267 |
| Blue Tit | Winter | 1yr-GM | 3+yr | 0.011 | 0.722 |
| Blue Tit | Winter | 2yr-GM | 3+yr | 0.022 | 0.808 |
| Chaffinch | Winter | 1yr-FAKT | 1yr-GM | 0.018 | 0.832 |
| Chaffinch | Winter | 1yr-FAKT | 2yr-GM | 0.028 | 0.910 |
| Chaffinch | Winter | 1yr-FAKT | 3+yr | 0.027 | 0.896 |
| Chaffinch | Winter | 1yr-GM | 2yr-GM | 0.010 | 0.779 |
| Chaffinch | Winter | 1yr-GM | 3+yr | 0.009 | 0.724 |
| Chaffinch | Winter | 2yr-GM | 3+yr | -0.001 | 0.468 |
| Goldfinch | Winter | 1yr-FAKT | 1yr-GM | 0.020 | 0.918 |
| Goldfinch | Winter | 1yr-FAKT | 2yr-GM | -0.007 | 0.390 |
| Goldfinch | Winter | 1yr-FAKT | 3+yr | 0.020 | 0.876 |
| Goldfinch | Winter | 1yr-GM | 2yr-GM | -0.027 | 0.045 |
| Goldfinch | Winter | 1yr-GM | 3+yr | 0.001 | 0.532 |
| Goldfinch | Winter | 2yr-GM | 3+yr | 0.028 | 0.923 |
| Great Tit | Winter | 1yr-FAKT | 1yr-GM | 0.032 | 0.938 |
| Great Tit | Winter | 1yr-FAKT | 2yr-GM | 0.034 | 0.928 |
| Great Tit | Winter | 1yr-FAKT | 3+yr | -0.013 | 0.393 |
| Great Tit | Winter | 1yr-GM | 2yr-GM | 0.002 | 0.588 |
| Great Tit | Winter | 1yr-GM | 3+yr | -0.049 | 0.104 |
| Great Tit | Winter | 2yr-GM | 3+yr | -0.050 | 0.108 |
| Greenfinch | Winter | 1yr-FAKT | 1yr-GM | 0.012 | 0.733 |
| Greenfinch | Winter | 1yr-FAKT | 2yr-GM | 0.015 | 0.724 |
| Greenfinch | Winter | 1yr-FAKT | 3+yr | 0.023 | 0.849 |
| Greenfinch | Winter | 1yr-GM | 2yr-GM | 0.003 | 0.577 |
| Greenfinch | Winter | 1yr-GM | 3+yr | 0.010 | 0.742 |
| Greenfinch | Winter | 2yr-GM | 3+yr | 0.006 | 0.642 |
| Linnet | Winter | 1yr-FAKT | 1yr-GM | -0.004 | 0.064 |
| Linnet | Winter | 1yr-FAKT | 2yr-GM | 0.000 | 0.732 |
| Linnet | Winter | 1yr-FAKT | 3+yr | -0.001 | 0.276 |
| Linnet | Winter | 1yr-GM | 2yr-GM | 0.004 | 0.934 |
| Linnet | Winter | 1yr-GM | 3+yr | 0.002 | 0.758 |
| Linnet | Winter | 2yr-GM | 3+yr | -0.001 | 0.179 |
| Reed Bunting | Winter | 1yr-FAKT | 1yr-GM | 0.008 | 0.907 |
| Reed Bunting | Winter | 1yr-FAKT | 2yr-GM | 0.002 | 0.614 |
| Reed Bunting | Winter | 1yr-FAKT | 3+yr | 0.005 | 0.745 |
| Reed Bunting | Winter | 1yr-GM | 2yr-GM | -0.005 | 0.109 |
| Reed Bunting | Winter | 1yr-GM | 3+yr | -0.002 | 0.334 |
| Reed Bunting | Winter | 2yr-GM | 3+yr | 0.003 | 0.661 |
| Tree Sparrow | Winter | 1yr-FAKT | 1yr-GM | -0.006 | 0.374 |
| Tree Sparrow | Winter | 1yr-FAKT | 2yr-GM | -0.002 | 0.466 |
| Tree Sparrow | Winter | 1yr-FAKT | 3+yr | -0.008 | 0.364 |
| Tree Sparrow | Winter | 1yr-GM | 2yr-GM | 0.003 | 0.546 |
| Tree Sparrow | Winter | 1yr-GM | 3+yr | -0.002 | 0.462 |
| Tree Sparrow | Winter | 2yr-GM | 3+yr | -0.005 | 0.433 |
| Yellowhammer | Winter | 1yr-FAKT | 1yr-GM | -0.026 | 0.035 |
| Yellowhammer | Winter | 1yr-FAKT | 2yr-GM | **-0.050** | **0.010** |
| Yellowhammer | Winter | 1yr-FAKT | 3+yr | **-0.062** | **0.005** |
| Yellowhammer | Winter | 1yr-GM | 2yr-GM | -0.023 | 0.146 |
| Yellowhammer | Winter | 1yr-GM | 3+yr | -0.034 | 0.099 |

| | | | | | |
|---|---|---|---|---|---|
| Yellowhammer | Winter | 2yr-GM | 3+yr | -0.011 | 0.381 |

| | | Fallow field types | | Δ bird incidence (Base-Reference) | |
|---|---|---|---|---|---|
| **Species** | **Season** | **Base** | **Reference** | **Median** | **Bayes *P*** |
| NSpecies | Autumn | 1yr-FAKT | 1yr-GM | -0.353 | 0.240 |
| NSpecies | Autumn | 1yr-FAKT | 2yr-GM | 0.730 | 0.916 |
| NSpecies | Autumn | 1yr-FAKT | 3+yr | 0.971 | 0.940 |
| NSpecies | Autumn | 1yr-GM | 2yr-GM | **1.087** | **0.994** |
| NSpecies | Autumn | 1yr-GM | 3+yr | **1.330** | **0.989** |
| NSpecies | Autumn | 2yr-GM | 3+yr | 0.246 | 0.672 |
| NSpecies | Winter | 1yr-FAKT | 1yr-GM | -0.207 | 0.296 |
| NSpecies | Winter | 1yr-FAKT | 2yr-GM | 0.044 | 0.547 |
| NSpecies | Winter | 1yr-FAKT | 3+yr | 0.104 | 0.606 |
| NSpecies | Winter | 1yr-GM | 2yr-GM | 0.250 | 0.764 |
| NSpecies | Winter | 1yr-GM | 3+yr | 0.311 | 0.808 |
| NSpecies | Winter | 2yr-GM | 3+yr | 0.065 | 0.566 |
