## Supplement D for "Criteria for effective fallow field eco schemes for farmland birds during non-breeding"

**Online Supplement 4**. Covariate raw data plots for all investigated species, including model predictions and their 95% credible intervals.

Grey dots show raw data, black dots (lines) with flags (grey shading) the model-predicted means and their 95% CrI for the following co-variate levels: fallow field type = 1yr-GM, year = 2019/20, farm track = no, secondary plant layer = yes, survey completeness = yes.

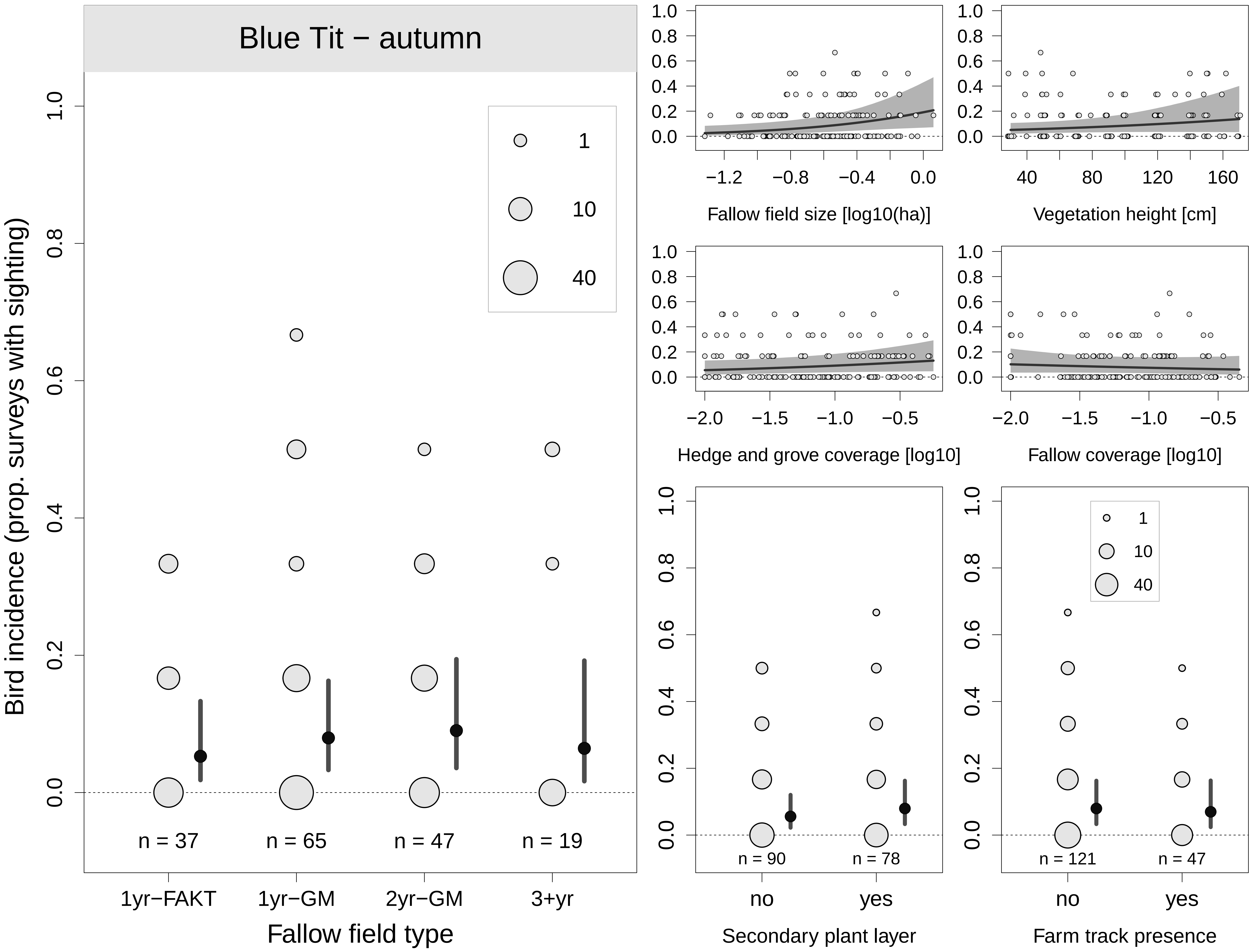
Blue Tit − autumn
Bird incidence (prop. surveys with sighting)
0.0
0.2
0.4
0.6
0.8
1.0
1
10
40
n = 37
n = 65
n = 47
n = 19
1yr−FAKT
1yr−GM
2yr−GM
3+yr
Fallow field type
−1.2
−0.8
−0.4
0.0
Fallow field size [log10(ha)]
40
80
120
160
Vegetation height [cm]
−2.0
−1.5
−1.0
−0.5
Hedge and grove coverage [log10]
Fallow coverage [log10]
n = 90
n = 78
no
yes
Secondary plant layer
n = 121
n = 47
Farm track presence

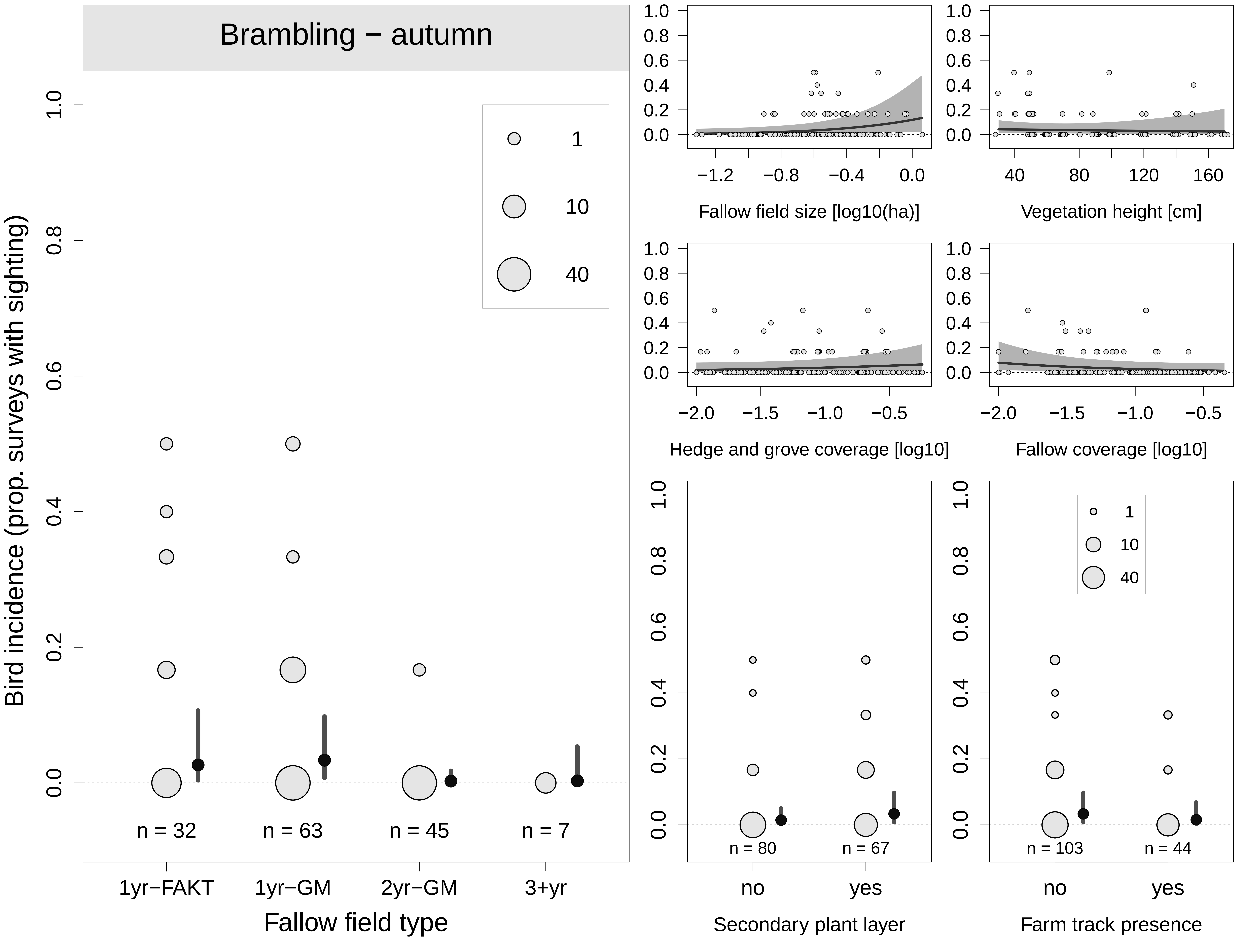
Brambling − autumn
Bird incidence (prop. surveys with sighting)
Fallow field type
1yr−FAKT
1yr−GM
2yr−GM
3+yr
n = 32
n = 63
n = 45
n = 7
1
10
40
Fallow field size [log10(ha)]
Vegetation height [cm]
Hedge and grove coverage [log10]
Fallow coverage [log10]
Secondary plant layer
Farm track presence
no
yes
n = 80
n = 67
n = 103
n = 44

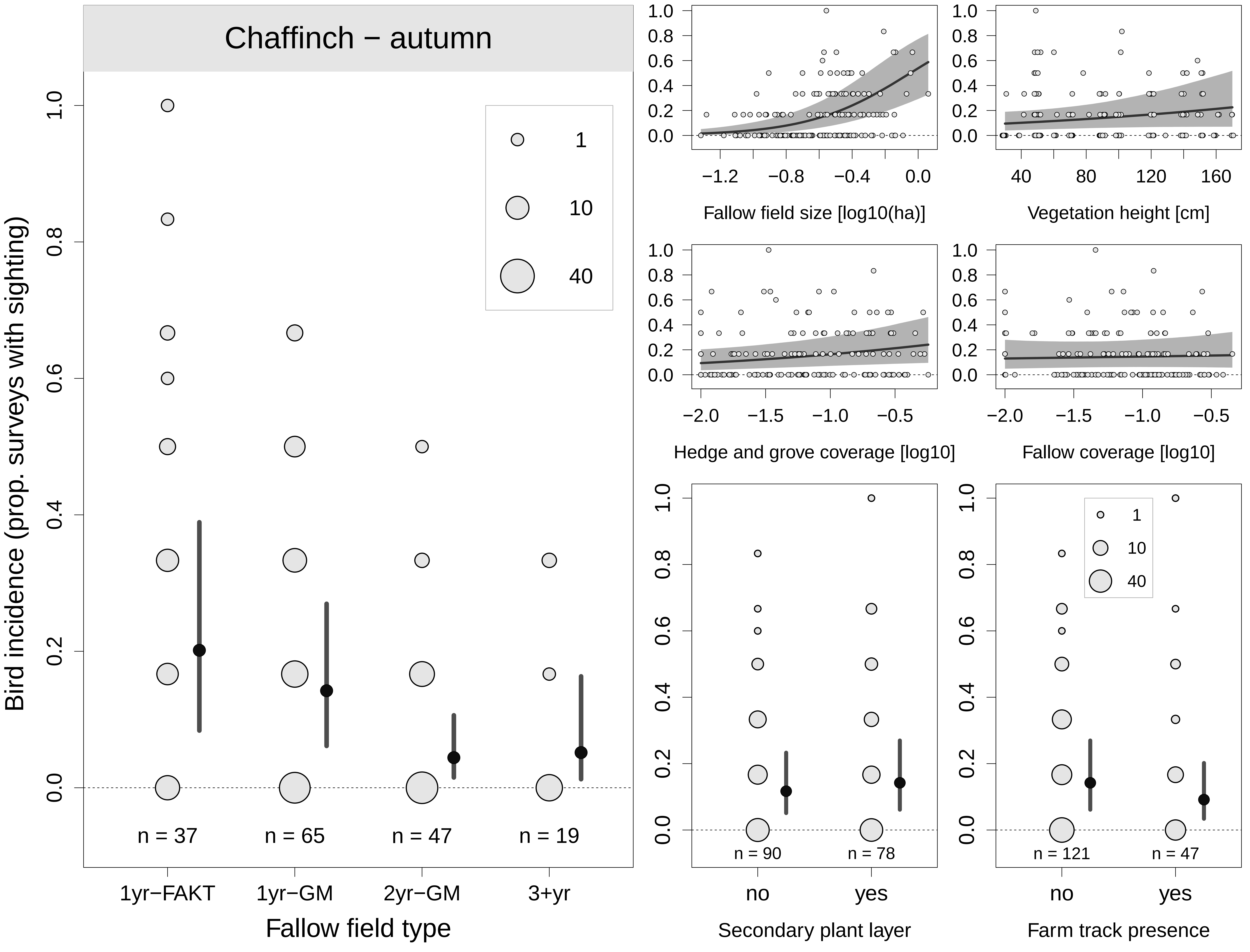
Chaffinch − autumn
Bird incidence (prop. surveys with sighting)
Fallow field type
1yr−FAKT
1yr−GM
2yr−GM
3+yr
n = 37
n = 65
n = 47
n = 19
1
10
40
Fallow field size [log10(ha)]
Vegetation height [cm]
Hedge and grove coverage [log10]
Fallow coverage [log10]
Secondary plant layer
Farm track presence
no
yes
n = 90
n = 78
n = 121
n = 47

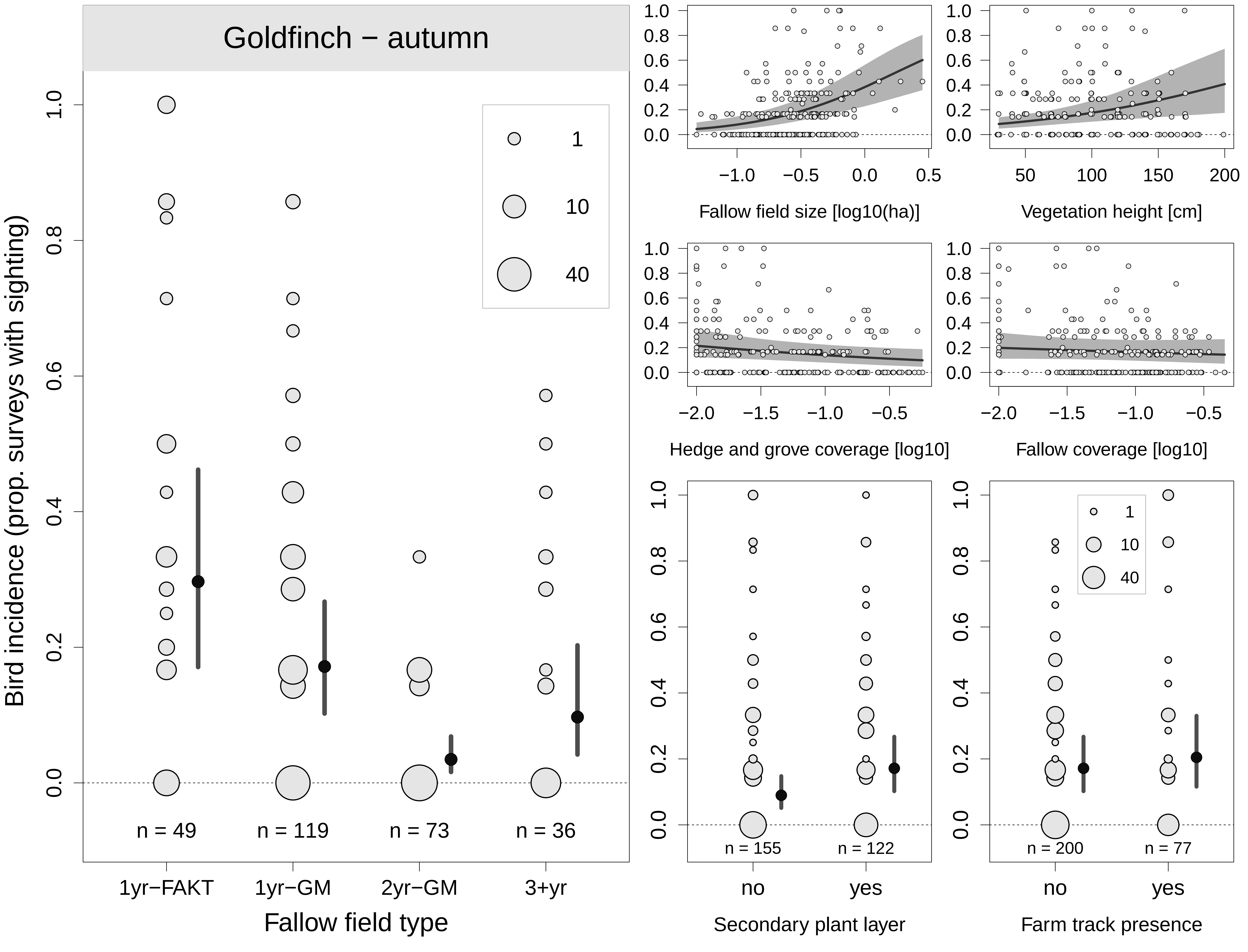

Goldfinch − autumn
Bird incidence (prop. surveys with sighting)
1
10
40
n = 49
n = 119
n = 73
n = 36
1yr−FAKT
1yr−GM
2yr−GM
3+yr
Fallow field type
Fallow field size [log10(ha)]
Vegetation height [cm]
Hedge and grove coverage [log10]
Fallow coverage [log10]
n = 155
n = 122
no
yes
Secondary plant layer
n = 200
n = 77
no
yes
Farm track presence

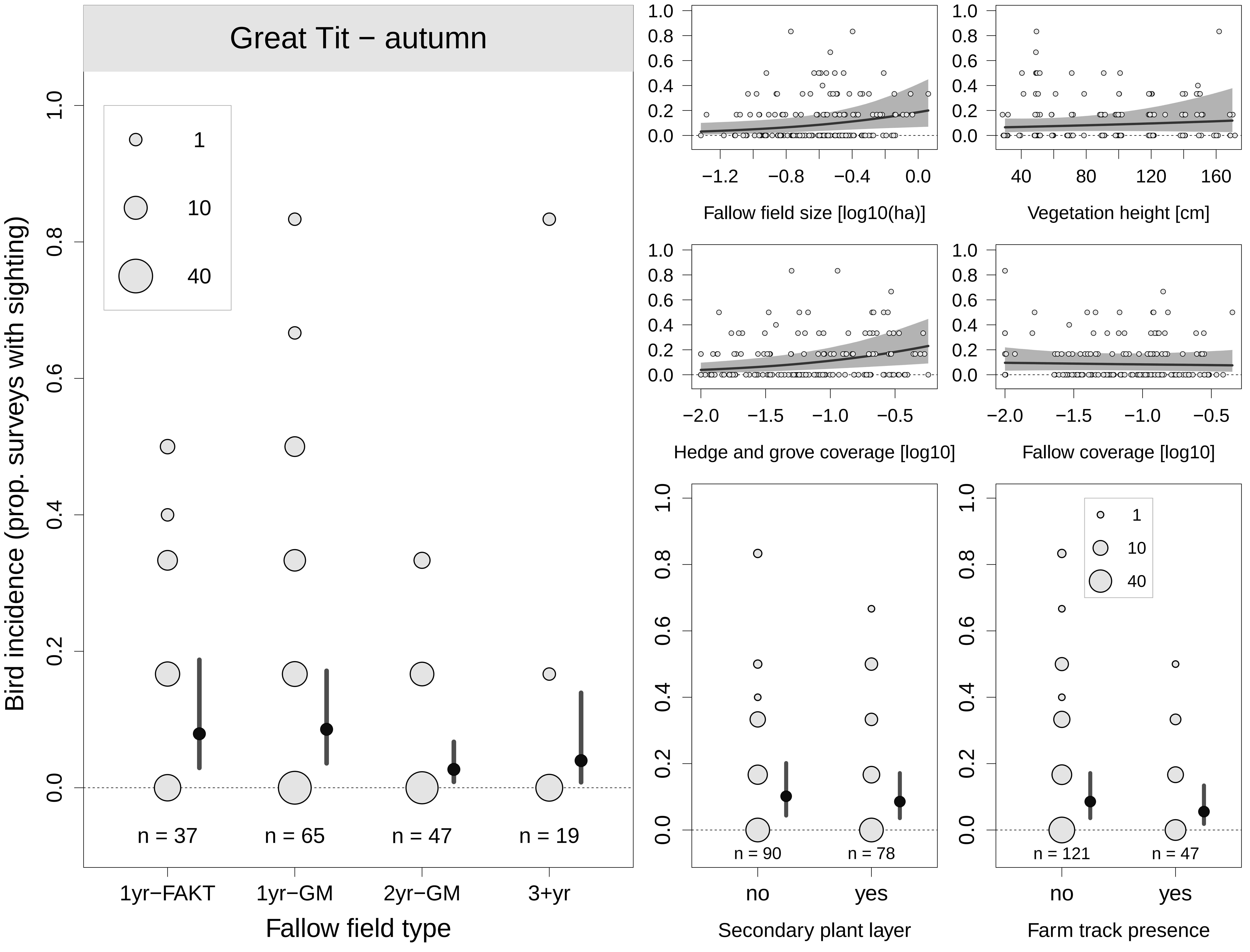

Great Tit − autumn
Bird incidence (prop. surveys with sighting)
1
10
40
n = 37
n = 65
n = 47
n = 19
1yr−FAKT
1yr−GM
2yr−GM
3+yr
Fallow field type
Fallow field size [log10(ha)]
Vegetation height [cm]
Hedge and grove coverage [log10]
Fallow coverage [log10]
n = 90
n = 78
no
yes
Secondary plant layer
n = 121
n = 47
no
yes
Farm track presence

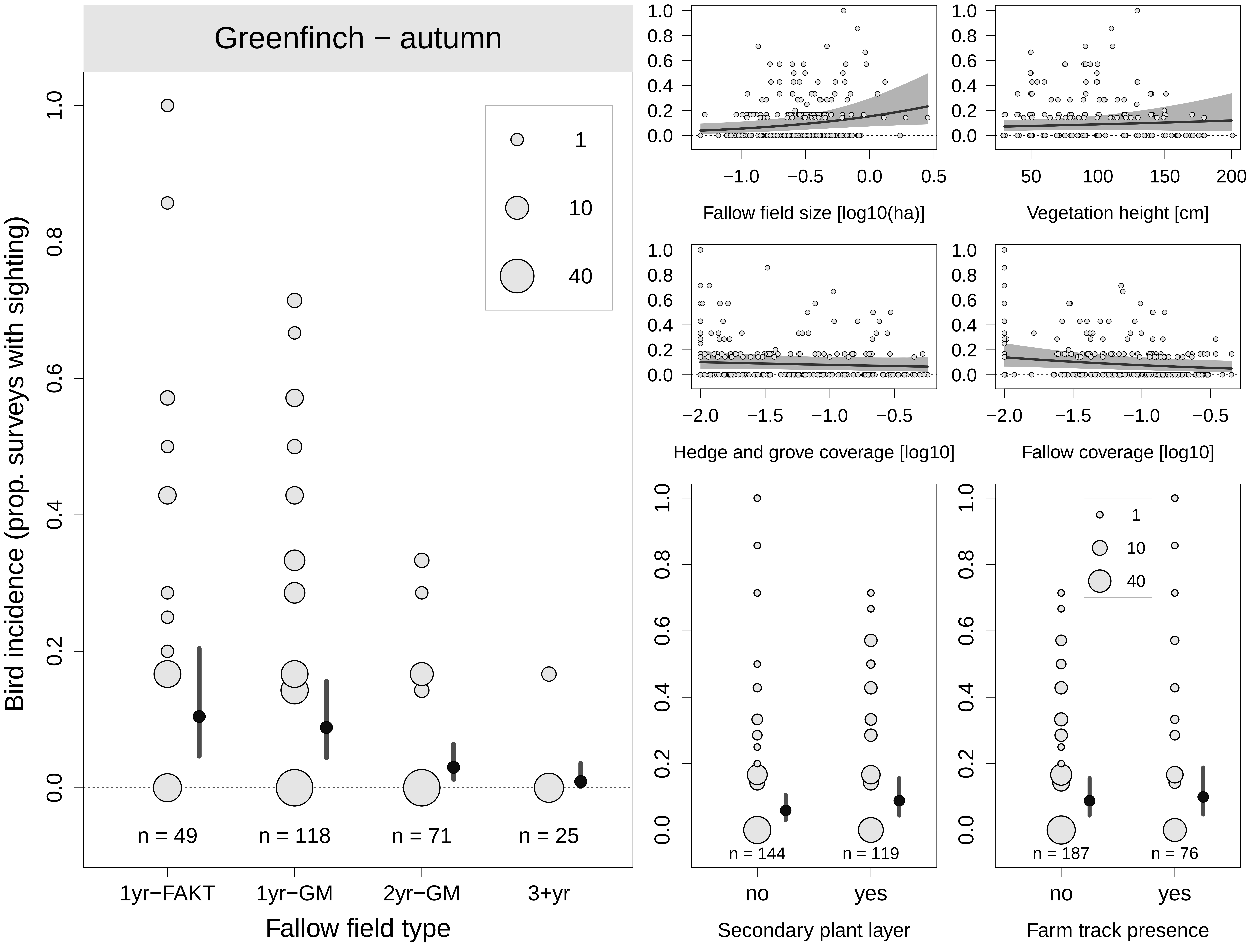

Greenfinch − autumn
Bird incidence (prop. surveys with sighting)
0.0
0.2
0.4
0.6
0.8
1.0
1
10
40
n = 49
n = 118
n = 71
n = 25
1yr−FAKT
1yr−GM
2yr−GM
3+yr
Fallow field type
−1.0
−0.5
0.0
0.5
Fallow field size [log10(ha)]
50
100
150
200
Vegetation height [cm]
−2.0
−1.5
−1.0
−0.5
Hedge and grove coverage [log10]
Fallow coverage [log10]
n = 144
n = 119
no
yes
Secondary plant layer
n = 187
n = 76
Farm track presence

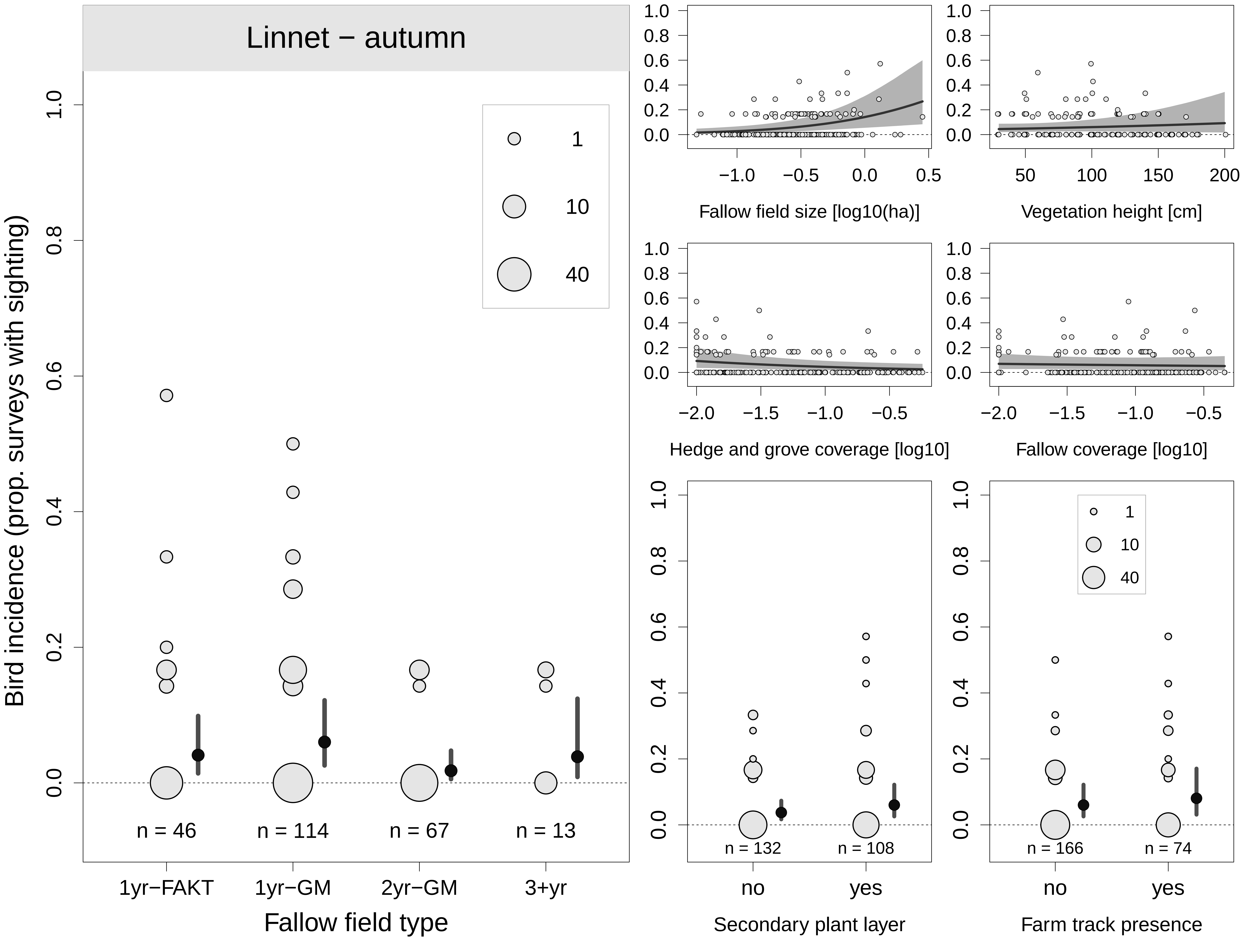

Linnet − autumn
Bird incidence (prop. surveys with sighting)
1
10
40
n = 46
n = 114
n = 67
n = 13
1yr−FAKT
1yr−GM
2yr−GM
3+yr
Fallow field type
Fallow field size [log10(ha)]
Vegetation height [cm]
Hedge and grove coverage [log10]
Fallow coverage [log10]
n = 132
n = 108
no
yes
Secondary plant layer
n = 166
n = 74
no
yes
Farm track presence

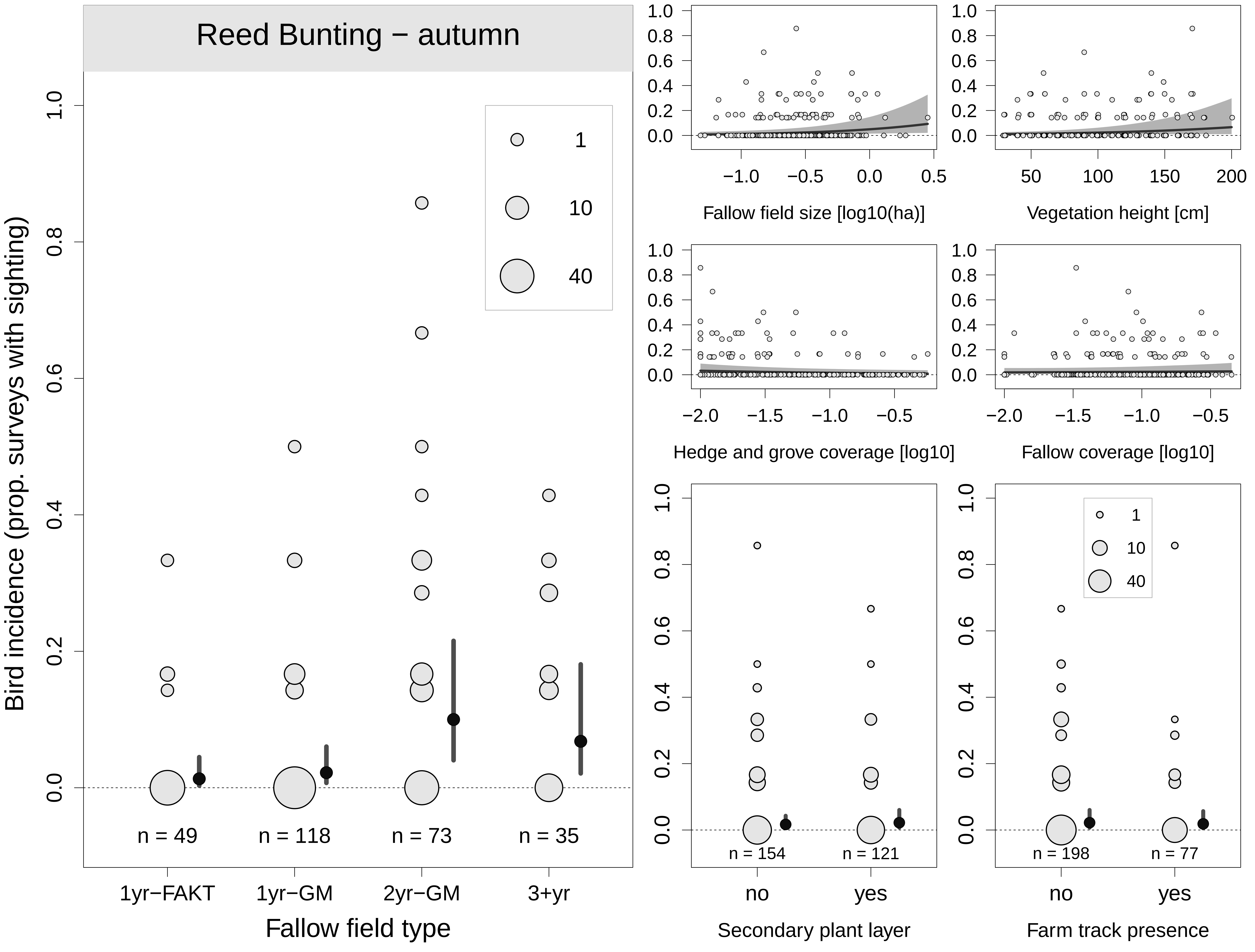

Reed Bunting − autumn
Bird incidence (prop. surveys with sighting)
1
10
40
n = 49
n = 118
n = 73
n = 35
1yr−FAKT
1yr−GM
2yr−GM
3+yr
Fallow field type
Fallow field size [log10(ha)]
Vegetation height [cm]
Hedge and grove coverage [log10]
Fallow coverage [log10]
n = 154
n = 121
no
yes
Secondary plant layer
n = 198
n = 77
Farm track presence

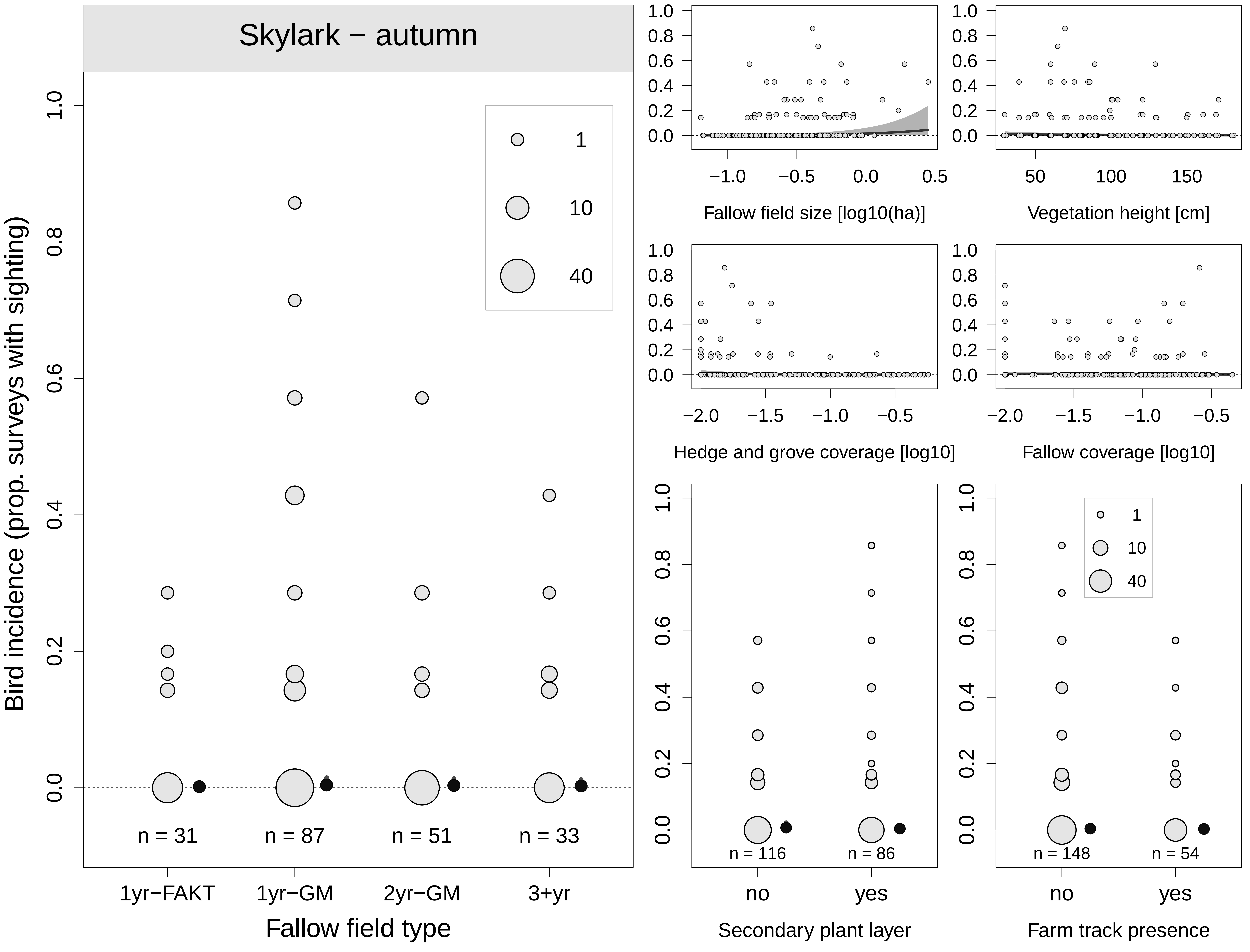

Skylark − autumn
Bird incidence (prop. surveys with sighting)
1
10
40
n = 31
n = 87
n = 51
n = 33
1yr−FAKT
1yr−GM
2yr−GM
3+yr
Fallow field type
Fallow field size [log10(ha)]
Vegetation height [cm]
Hedge and grove coverage [log10]
Fallow coverage [log10]
n = 116
n = 86
no
yes
Secondary plant layer
n = 148
n = 54
no
yes
Farm track presence

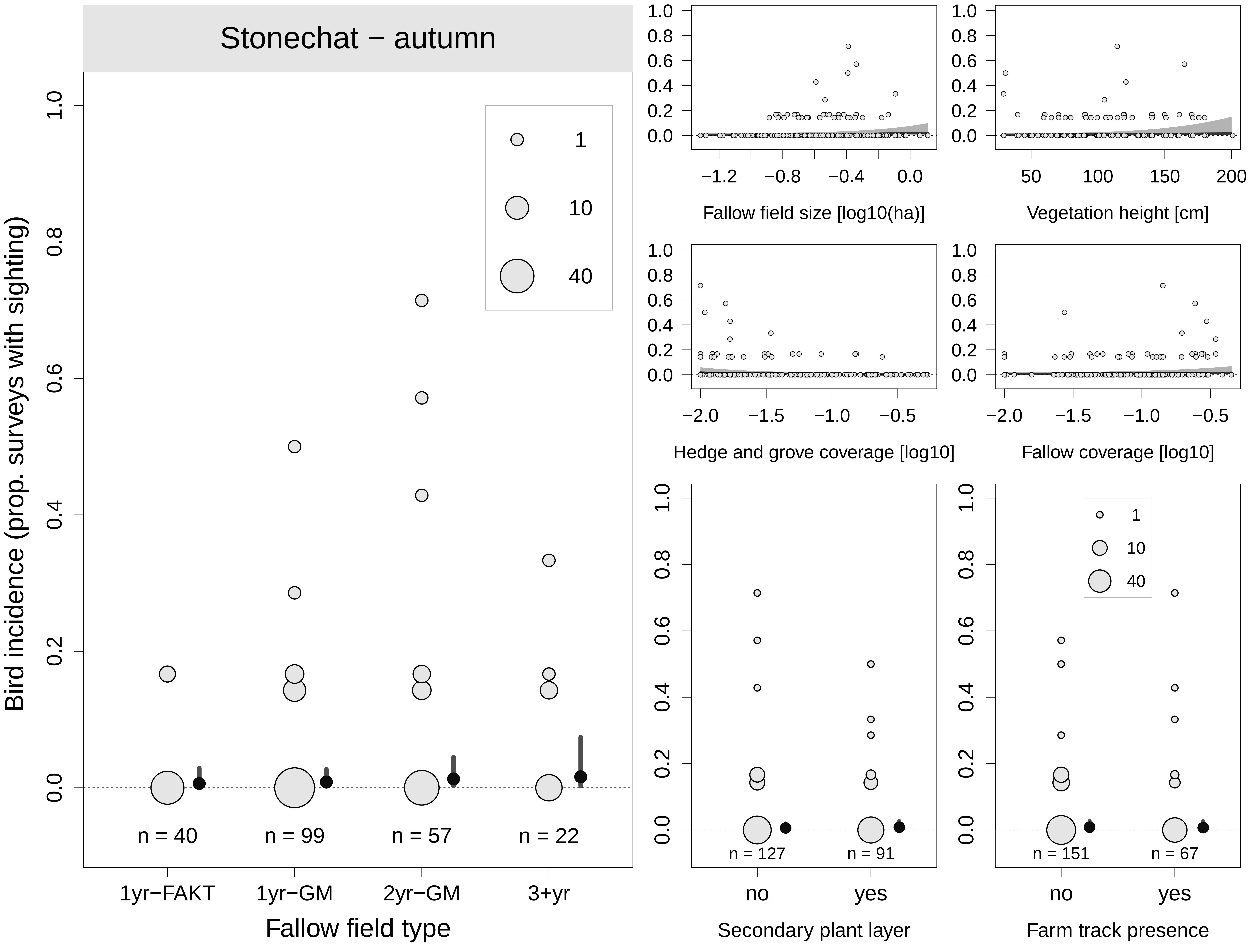

Stonechat − autumn
Bird incidence (prop. surveys with sighting)
1
10
40
n = 40
n = 99
n = 57
n = 22
1yr−FAKT
1yr−GM
2yr−GM
3+yr
Fallow field type
Fallow field size [log10(ha)]
Vegetation height [cm]
Hedge and grove coverage [log10]
Fallow coverage [log10]
n = 127
n = 91
no
yes
Secondary plant layer
n = 151
n = 67
no
yes
Farm track presence

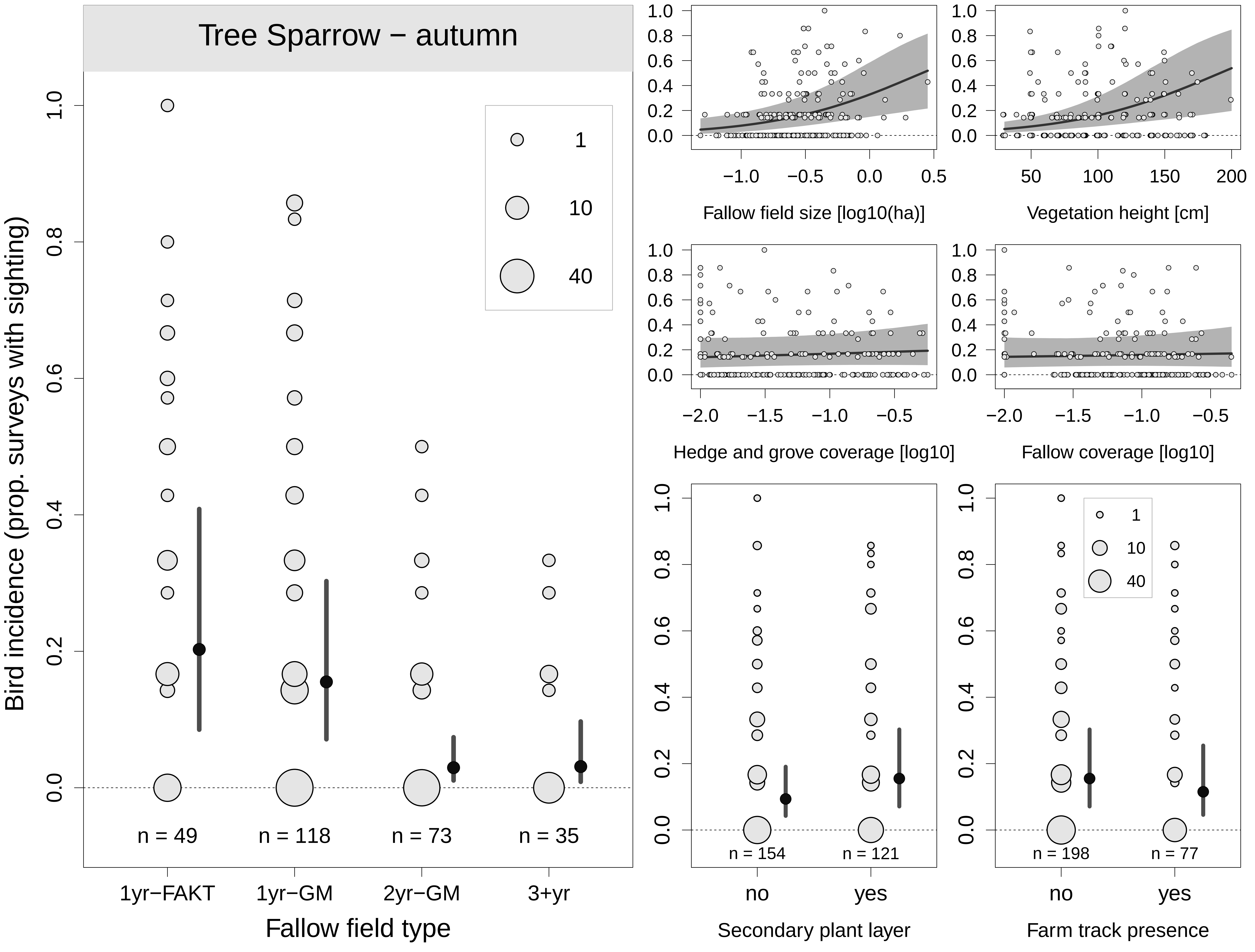

Tree Sparrow − autumn
Bird incidence (prop. surveys with sighting)
1
10
40
n = 49
n = 118
n = 73
n = 35
1yr−FAKT
1yr−GM
2yr−GM
3+yr
Fallow field type
Fallow field size [log10(ha)]
Vegetation height [cm]
Hedge and grove coverage [log10]
Fallow coverage [log10]
n = 154
n = 121
no
yes
Secondary plant layer
n = 198
n = 77
no
yes
Farm track presence

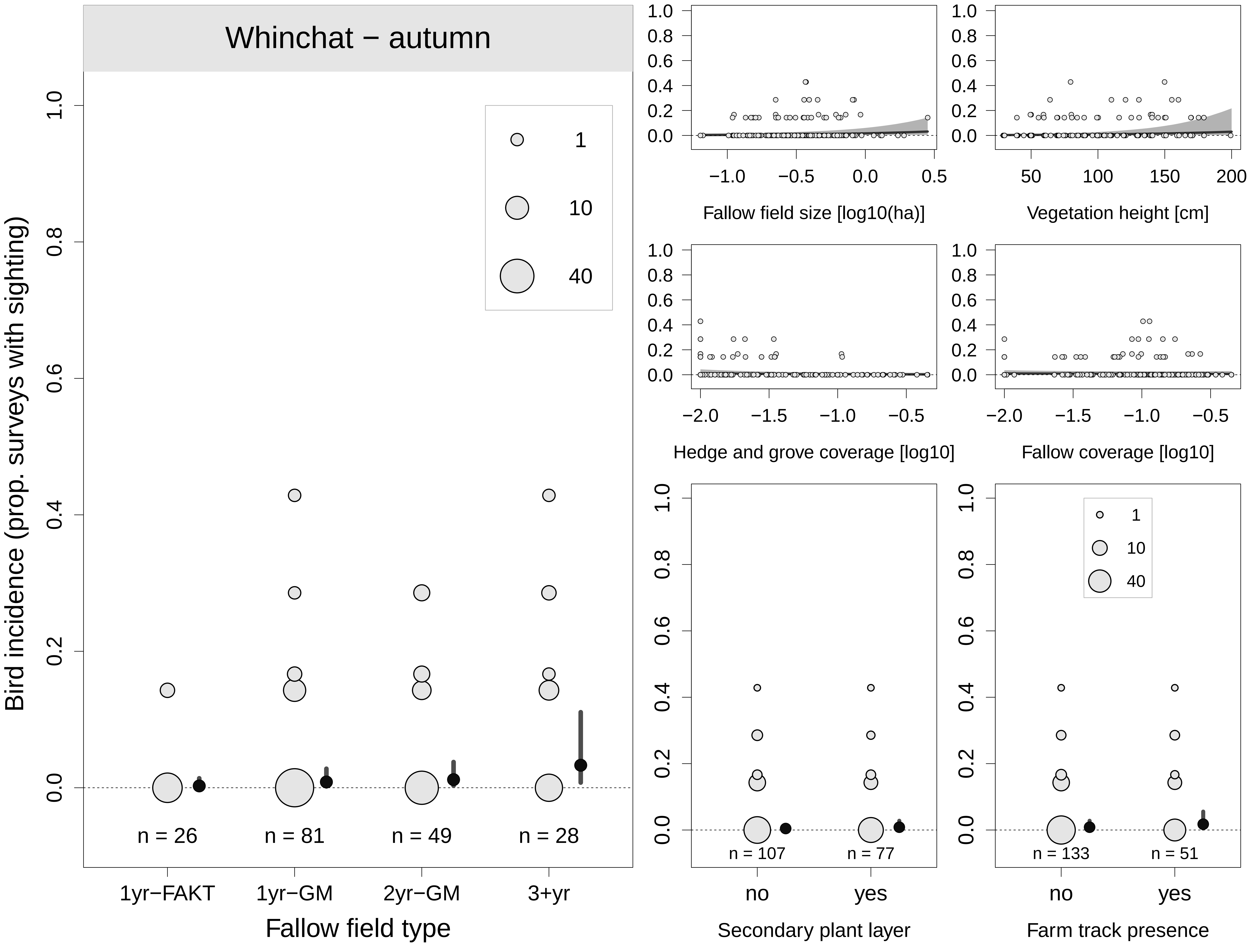
Whinchat − autumn
Bird incidence (prop. surveys with sighting)
1.0
0.8
0.6
0.4
0.2
0.0
1
10
40
n = 26
n = 81
n = 49
n = 28
1yr−FAKT
1yr−GM
2yr−GM
3+yr
Fallow field type
−1.0
−0.5
0.0
0.5
Fallow field size [log10(ha)]
50
100
150
200
Vegetation height [cm]
−2.0
−1.5
−1.0
−0.5
Hedge and grove coverage [log10]
Fallow coverage [log10]
n = 107
n = 77
no
yes
Secondary plant layer
n = 133
n = 51
Farm track presence

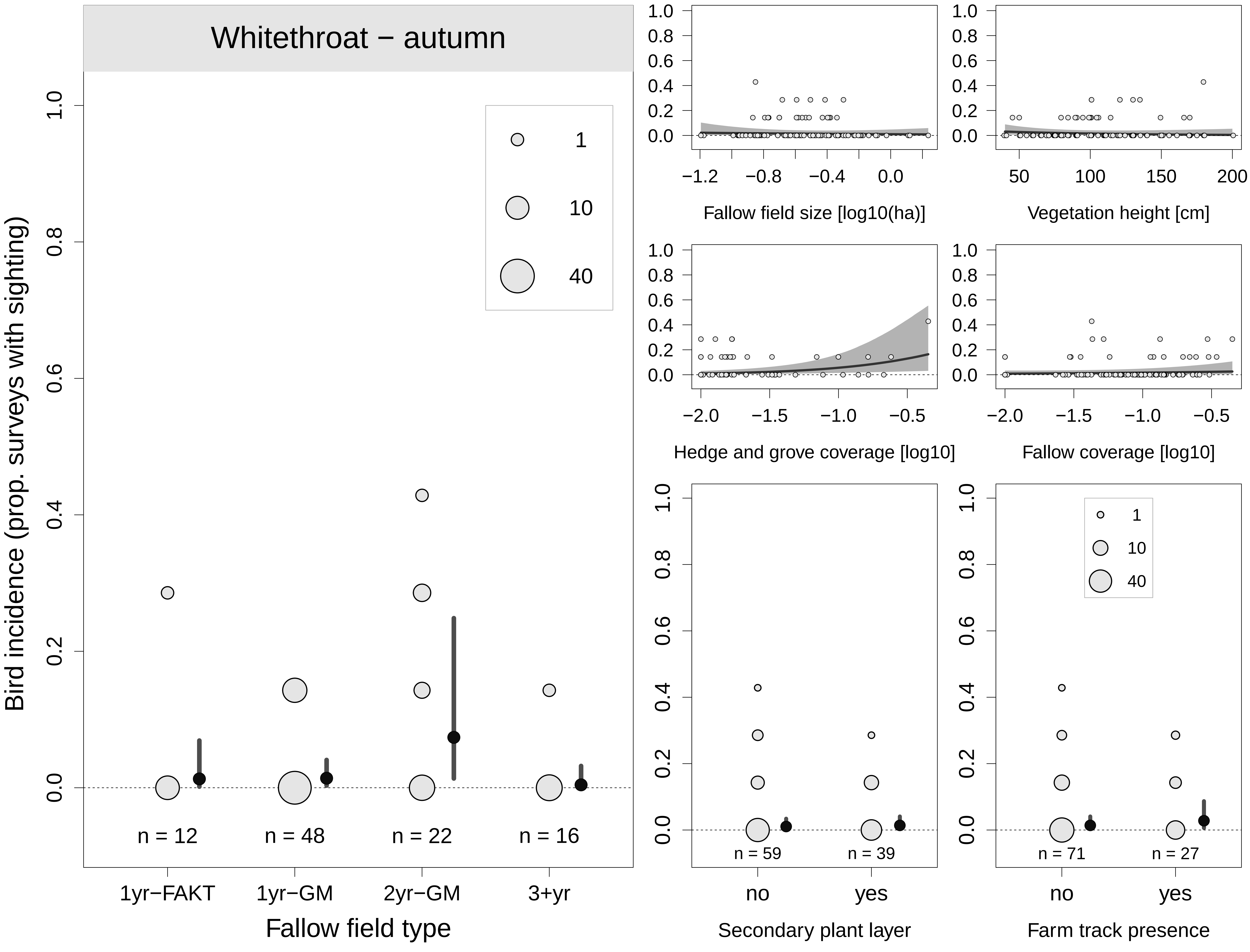

Whitethroat − autumn
Bird incidence (prop. surveys with sighting)
1.0
0.8
0.6
0.4
0.2
0.0
1
10
40
n = 12
n = 48
n = 22
n = 16
1yr−FAKT
1yr−GM
2yr−GM
3+yr
Fallow field type
Fallow field size [log10(ha)]
Vegetation height [cm]
Hedge and grove coverage [log10]
Fallow coverage [log10]
n = 59
n = 39
no
yes
Secondary plant layer
n = 71
n = 27
no
yes
Farm track presence

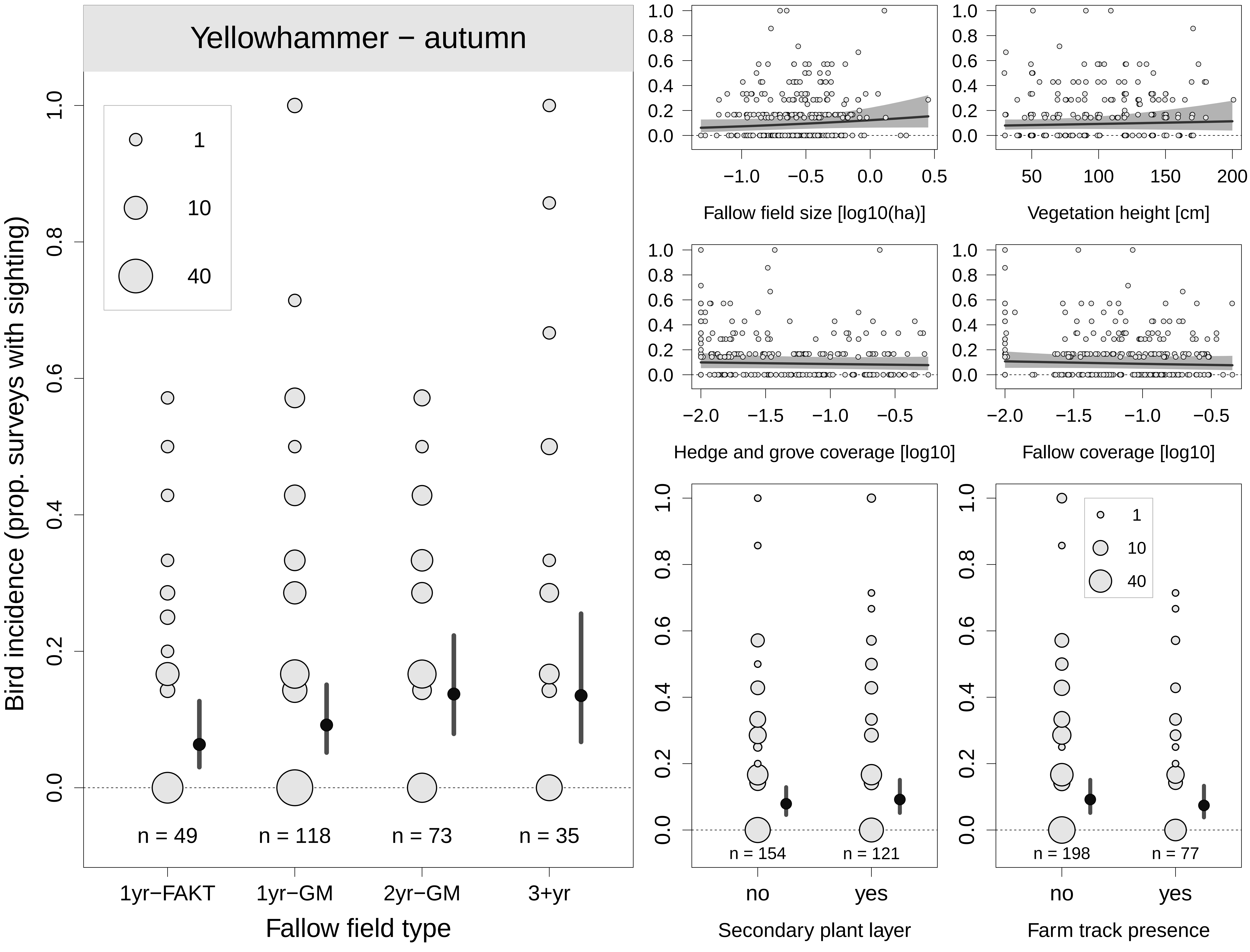

Yellowhammer − autumn
Bird incidence (prop. surveys with sighting)
1
10
40
n = 49
n = 118
n = 73
n = 35
1yr−FAKT
1yr−GM
2yr−GM
3+yr
Fallow field type
Fallow field size [log10(ha)]
Vegetation height [cm]
Hedge and grove coverage [log10]
Fallow coverage [log10]
n = 154
n = 121
no
yes
Secondary plant layer
n = 198
n = 77
no
yes
Farm track presence

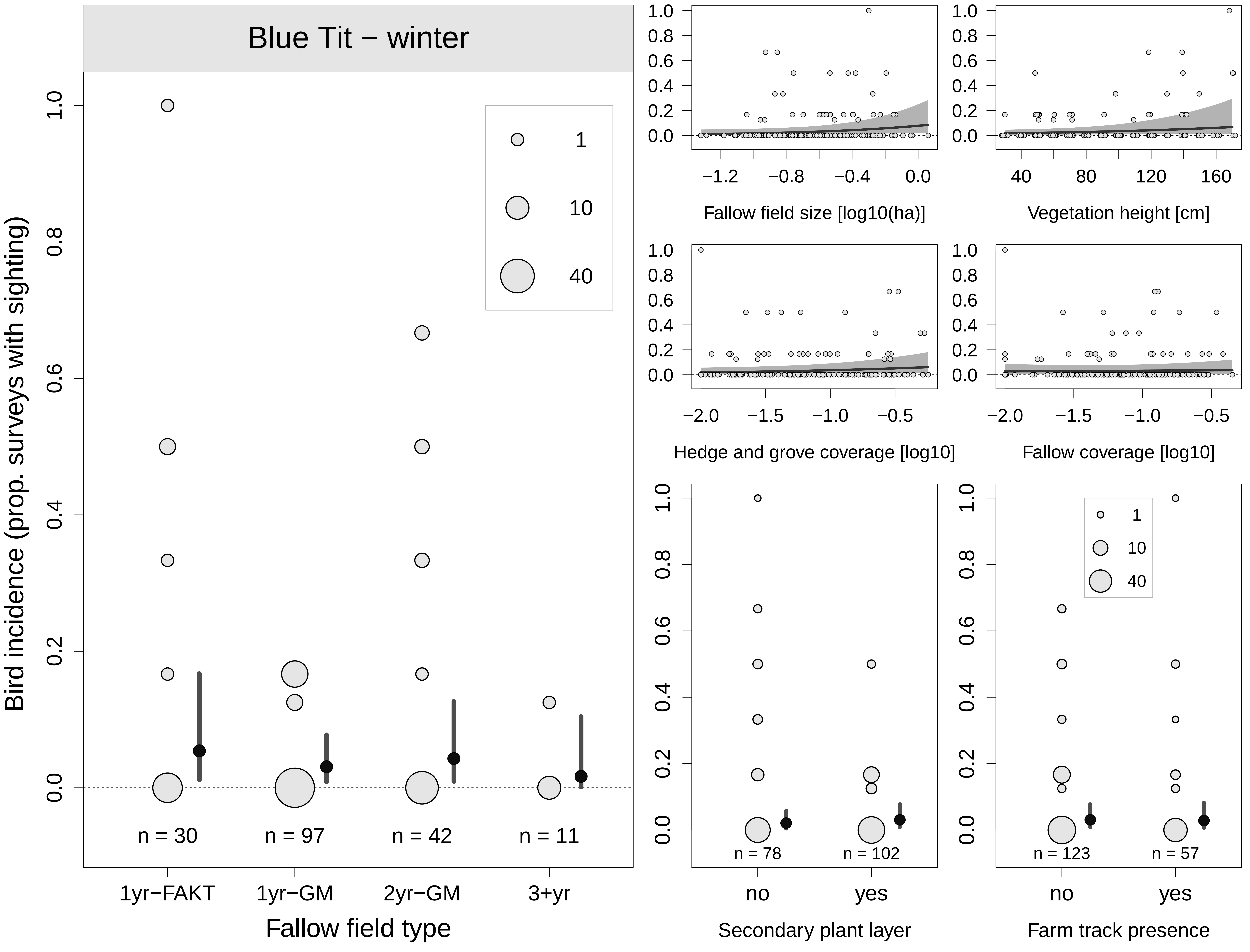
Blue Tit − winter
Bird incidence (prop. surveys with sighting)
Fallow field type
1yr−FAKT
1yr−GM
2yr−GM
3+yr
n = 30
n = 97
n = 42
n = 11
Fallow field size [log10(ha)]
Vegetation height [cm]
Hedge and grove coverage [log10]
Fallow coverage [log10]
Secondary plant layer
Farm track presence
no
yes
n = 78
n = 102
n = 123
n = 57

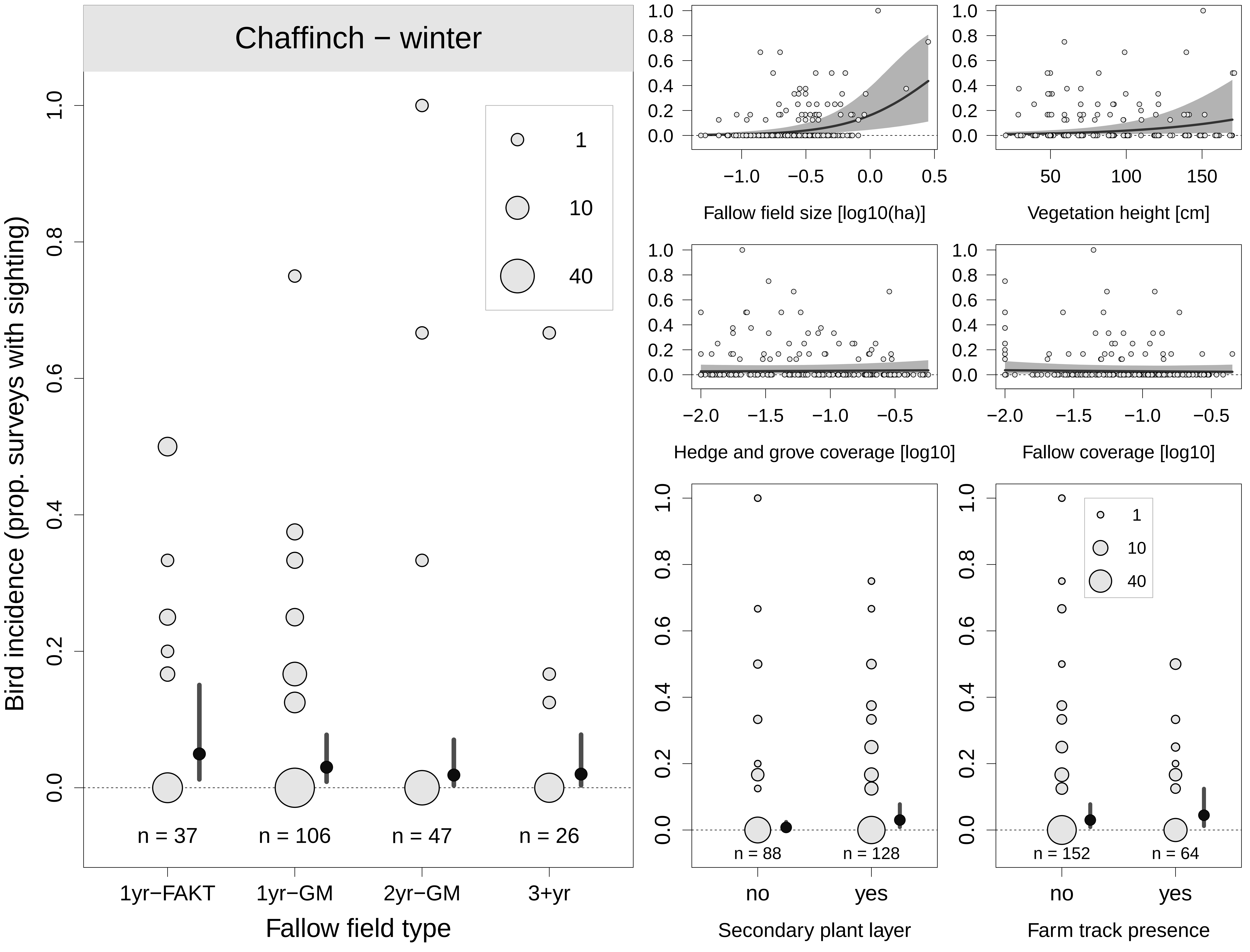

Chaffinch − winter
Bird incidence (prop. surveys with sighting)
1
10
40
n = 37
n = 106
n = 47
n = 26
1yr−FAKT
1yr−GM
2yr−GM
3+yr
Fallow field type
Fallow field size [log10(ha)]
Vegetation height [cm]
Hedge and grove coverage [log10]
Fallow coverage [log10]
n = 88
n = 128
no
yes
Secondary plant layer
n = 152
n = 64
no
yes
Farm track presence

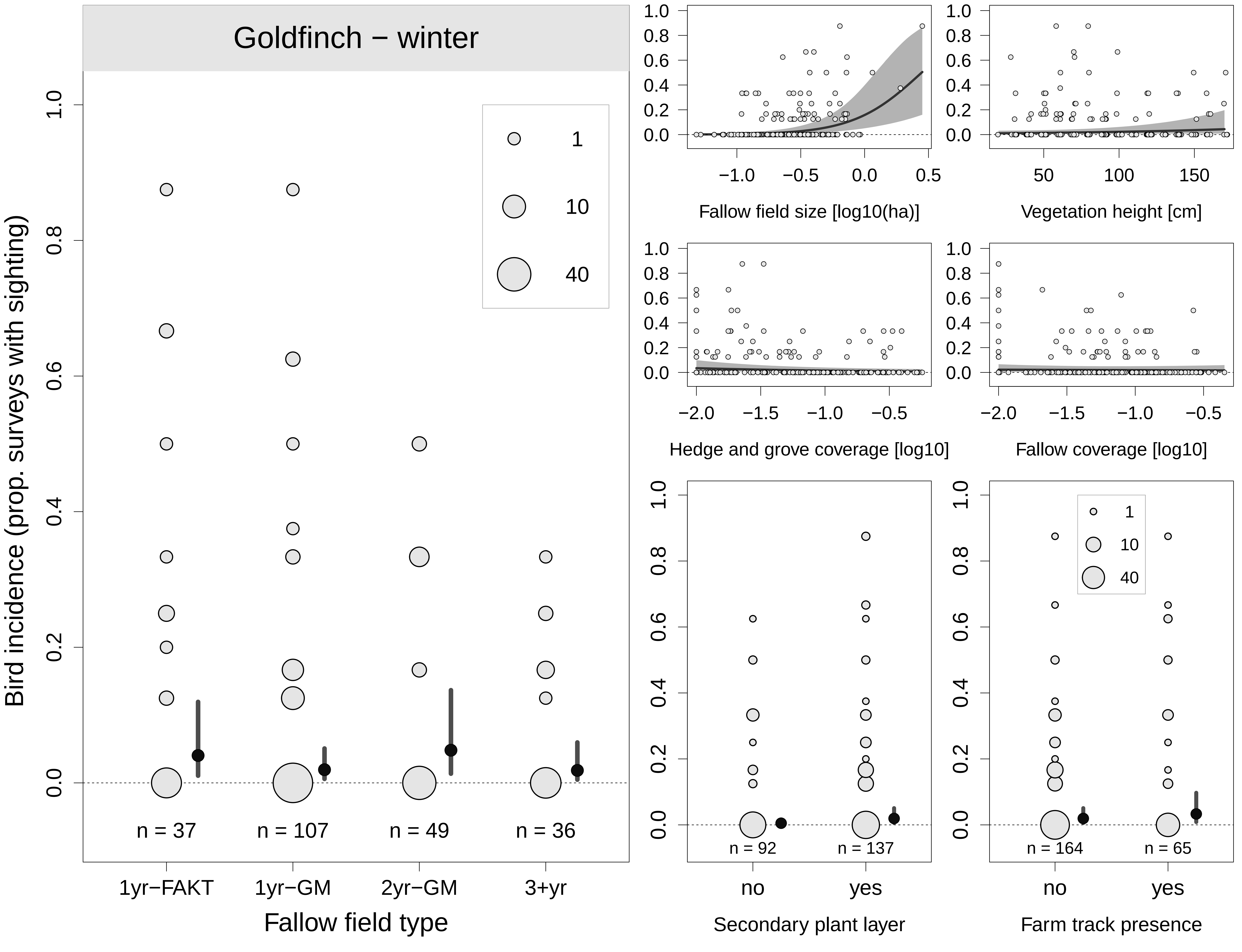

Goldfinch − winter
Bird incidence (prop. surveys with sighting)
1.0
0.8
0.6
0.4
0.2
0.0
1
10
40
n = 37
n = 107
n = 49
n = 36
1yr−FAKT
1yr−GM
2yr−GM
3+yr
Fallow field type
Fallow field size [log10(ha)]
Vegetation height [cm]
Hedge and grove coverage [log10]
Fallow coverage [log10]
n = 92
n = 137
no
yes
Secondary plant layer
n = 164
n = 65
Farm track presence

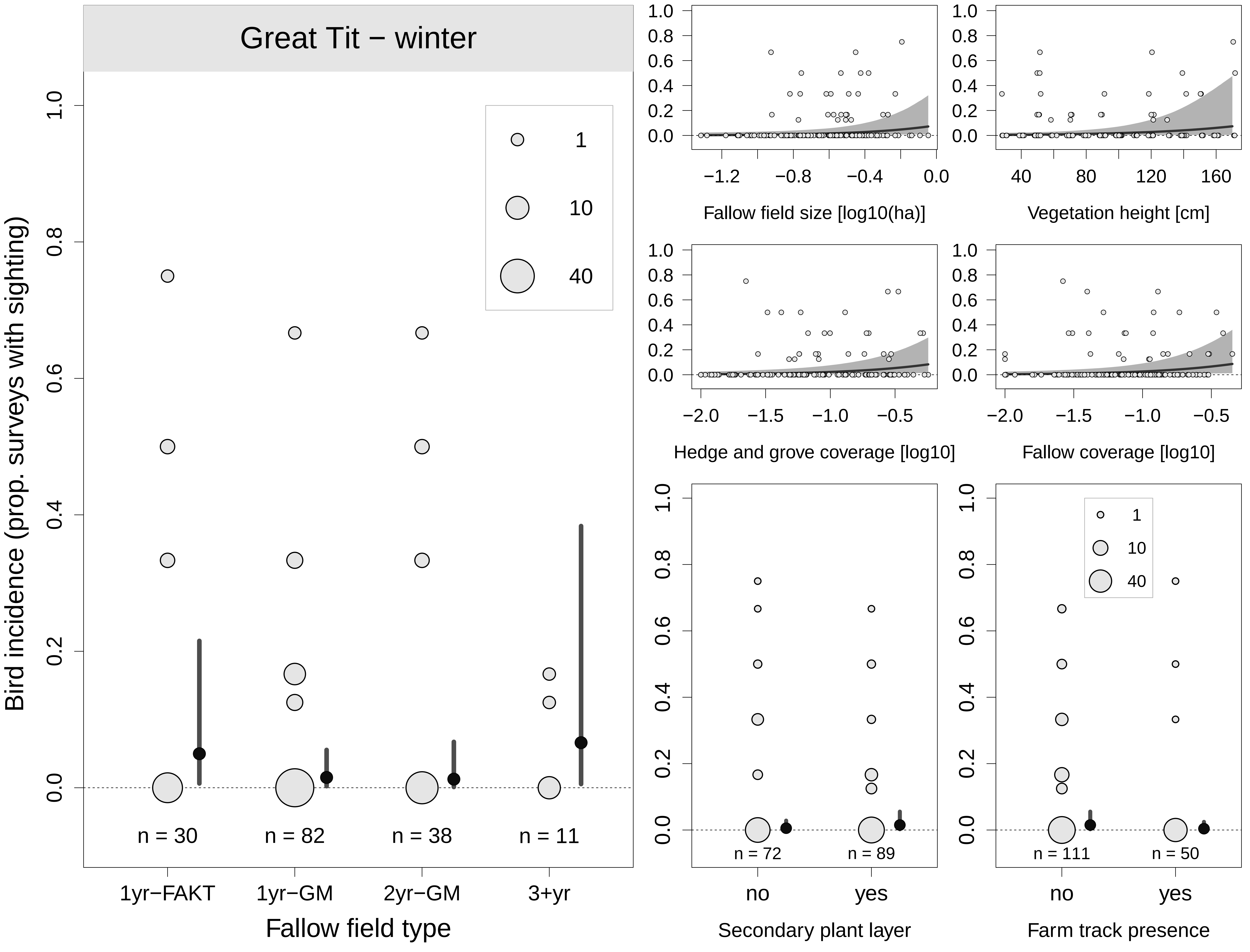

Great Tit − winter
Bird incidence (prop. surveys with sighting)
1
10
40
n = 30
n = 82
n = 38
n = 11
1yr−FAKT
1yr−GM
2yr−GM
3+yr
Fallow field type
Fallow field size [log10(ha)]
Vegetation height [cm]
Hedge and grove coverage [log10]
Fallow coverage [log10]
n = 72
n = 89
no
yes
Secondary plant layer
n = 111
n = 50
no
yes
Farm track presence

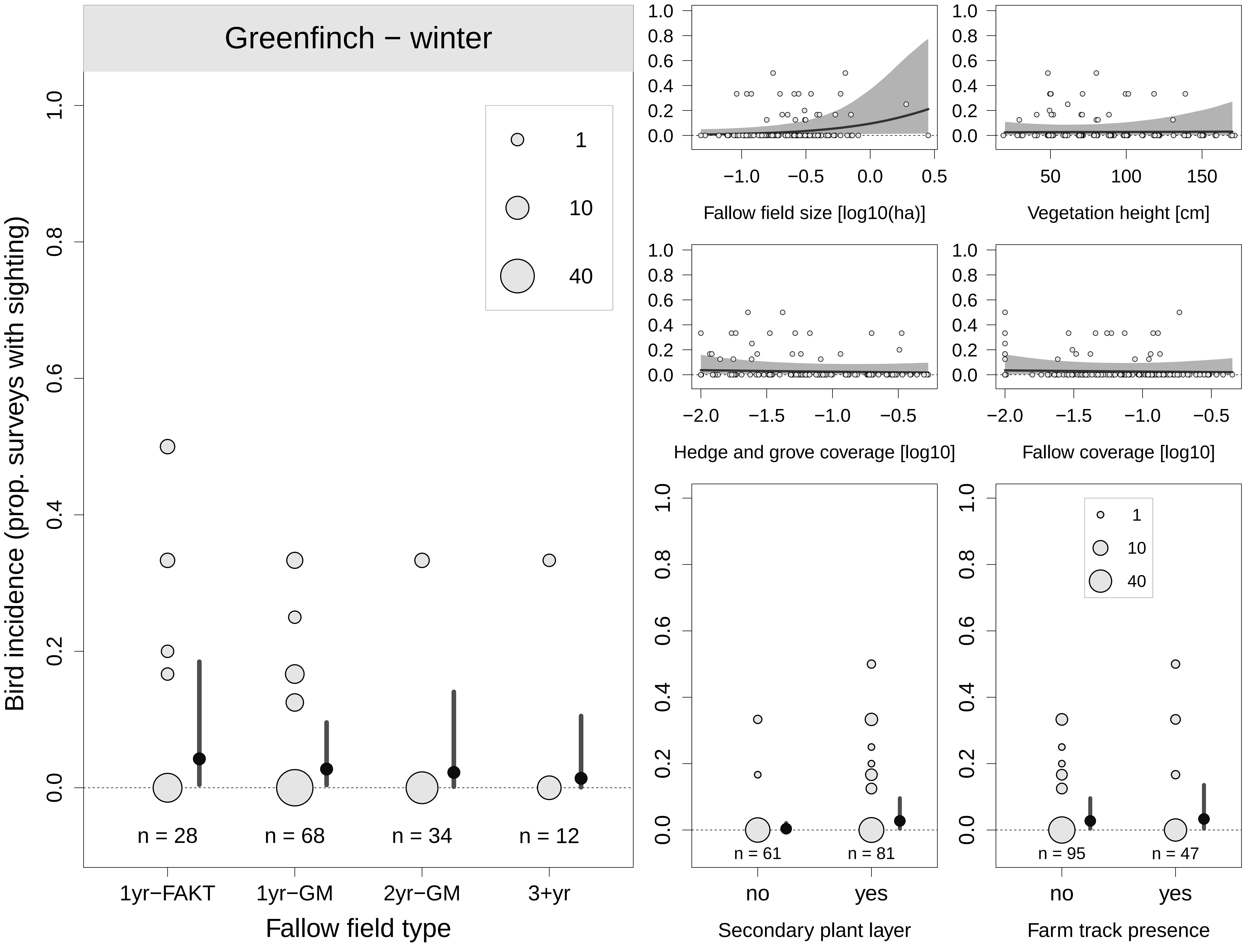
Greenfinch − winter
Bird incidence (prop. surveys with sighting)
1
10
40
n = 28
n = 68
n = 34
n = 12
1yr−FAKT
1yr−GM
2yr−GM
3+yr
Fallow field type
Fallow field size [log10(ha)]
Vegetation height [cm]
Hedge and grove coverage [log10]
Fallow coverage [log10]
n = 61
n = 81
no
yes
Secondary plant layer
n = 95
n = 47
no
yes
Farm track presence

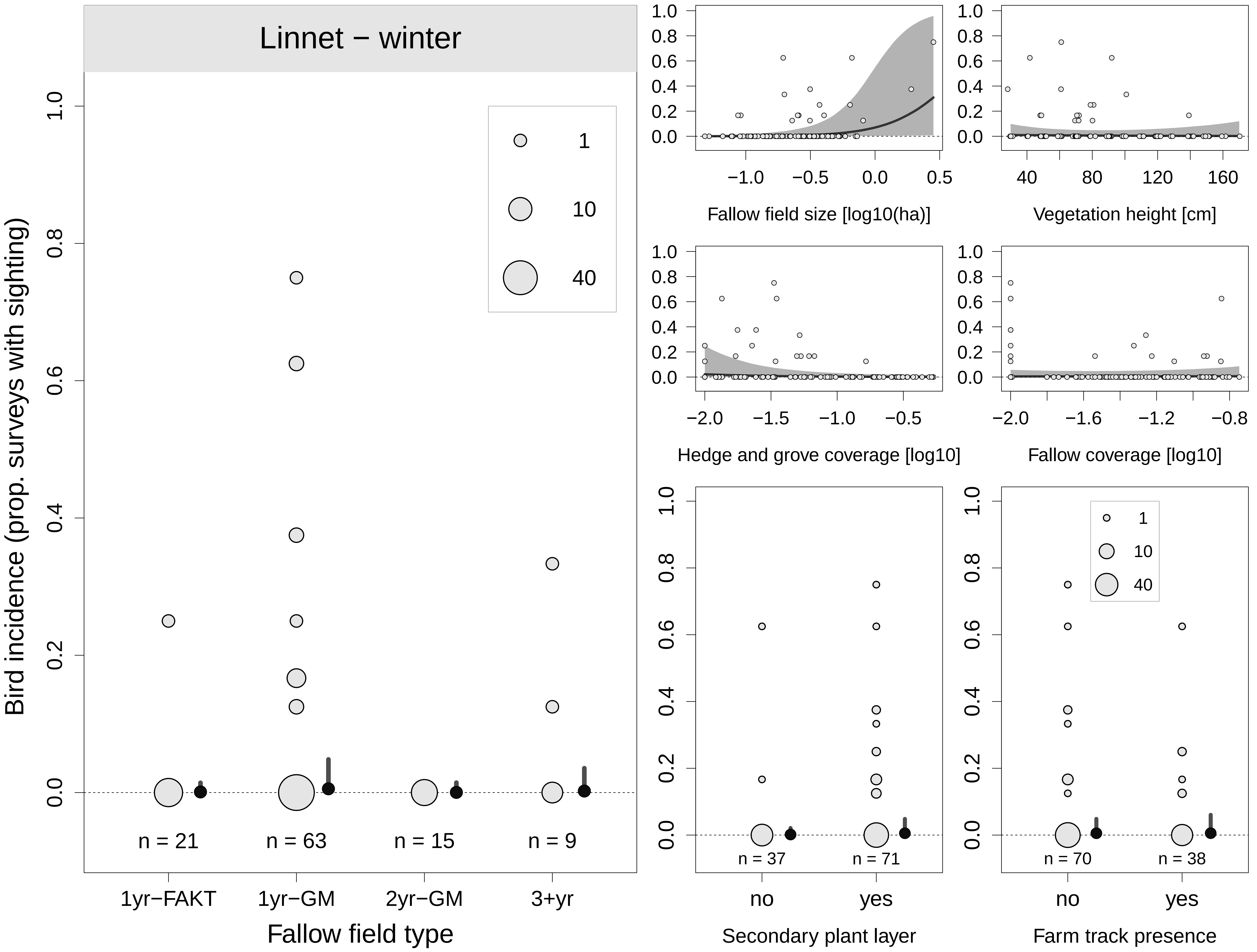

Linnet − winter
Bird incidence (prop. surveys with sighting)
1
10
40
n = 21
n = 63
n = 15
n = 9
1yr−FAKT
1yr−GM
2yr−GM
3+yr
Fallow field type
Fallow field size [log10(ha)]
Vegetation height [cm]
Hedge and grove coverage [log10]
Fallow coverage [log10]
n = 37
n = 71
no
yes
Secondary plant layer
n = 70
n = 38
no
yes
Farm track presence

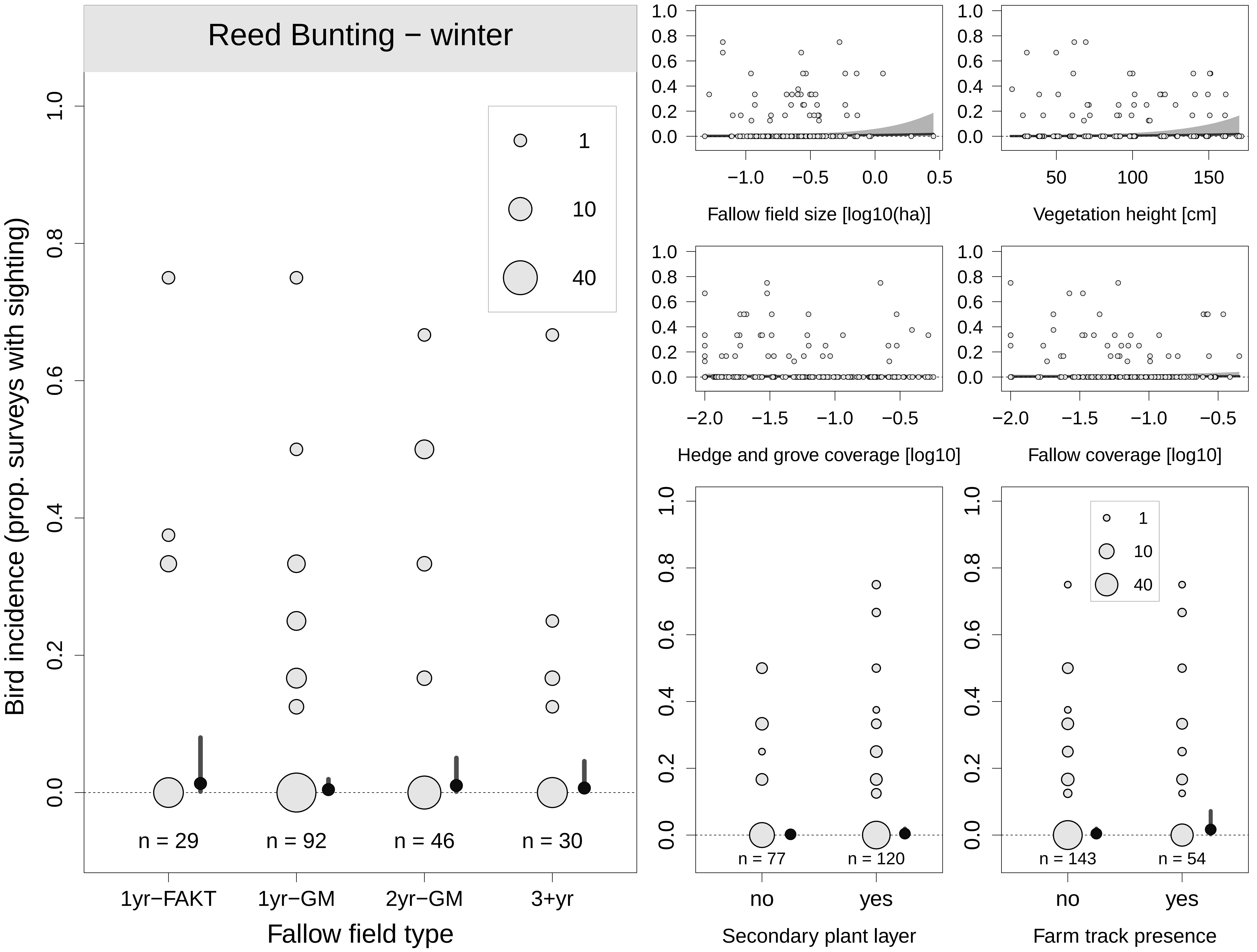

Reed Bunting − winter
Bird incidence (prop. surveys with sighting)
1
10
40
n = 29
n = 92
n = 46
n = 30
1yr−FAKT
1yr−GM
2yr−GM
3+yr
Fallow field type
Fallow field size [log10(ha)]
Vegetation height [cm]
Hedge and grove coverage [log10]
Fallow coverage [log10]
n = 77
n = 120
no
yes
Secondary plant layer
n = 143
n = 54
Farm track presence

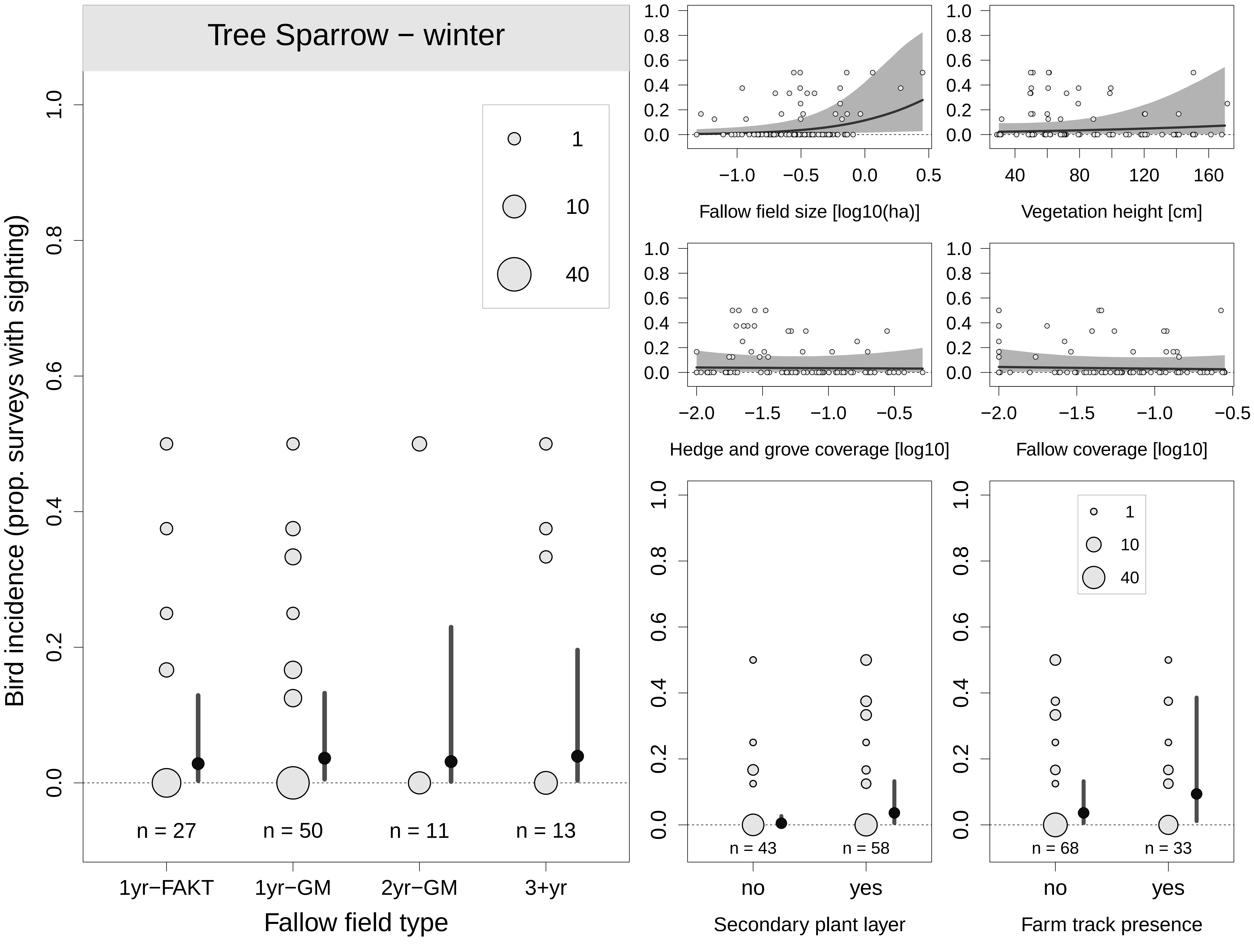

Tree Sparrow − winter
Bird incidence (prop. surveys with sighting)
1
10
40
n = 27
n = 50
n = 11
n = 13
1yr−FAKT
1yr−GM
2yr−GM
3+yr
Fallow field type
Fallow field size [log10(ha)]
Vegetation height [cm]
Hedge and grove coverage [log10]
Fallow coverage [log10]
n = 43
n = 58
no
yes
Secondary plant layer
n = 68
n = 33
no
yes
Farm track presence

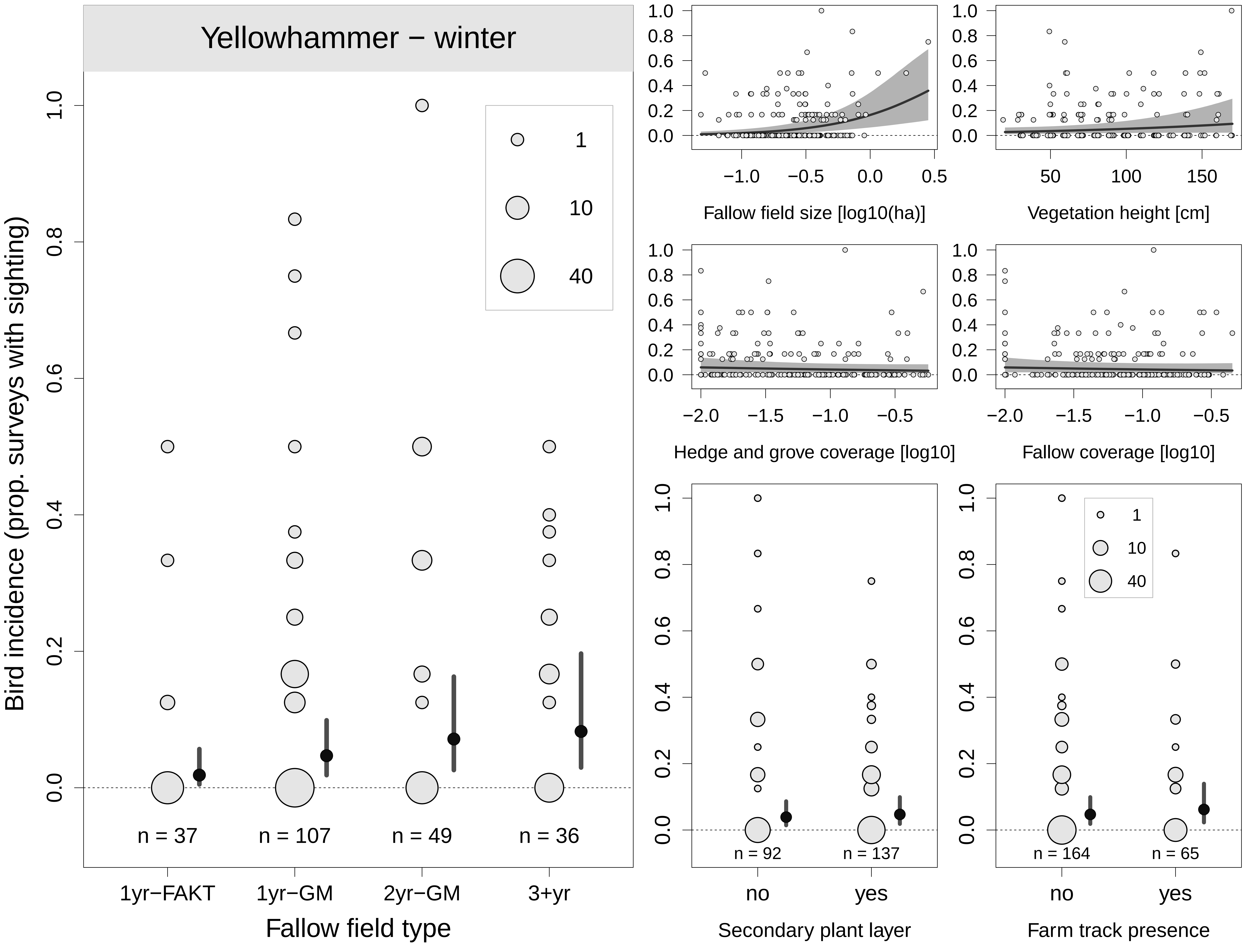

Yellowhammer − winter
Bird incidence (prop. surveys with sighting)
1
10
40
n = 37
n = 107
n = 49
n = 36
1yr−FAKT
1yr−GM
2yr−GM
3+yr
Fallow field type
Fallow field size [log10(ha)]
Vegetation height [cm]
Hedge and grove coverage [log10]
Fallow coverage [log10]
n = 92
n = 137
no
yes
Secondary plant layer
n = 164
n = 65
no
yes
Farm track presence

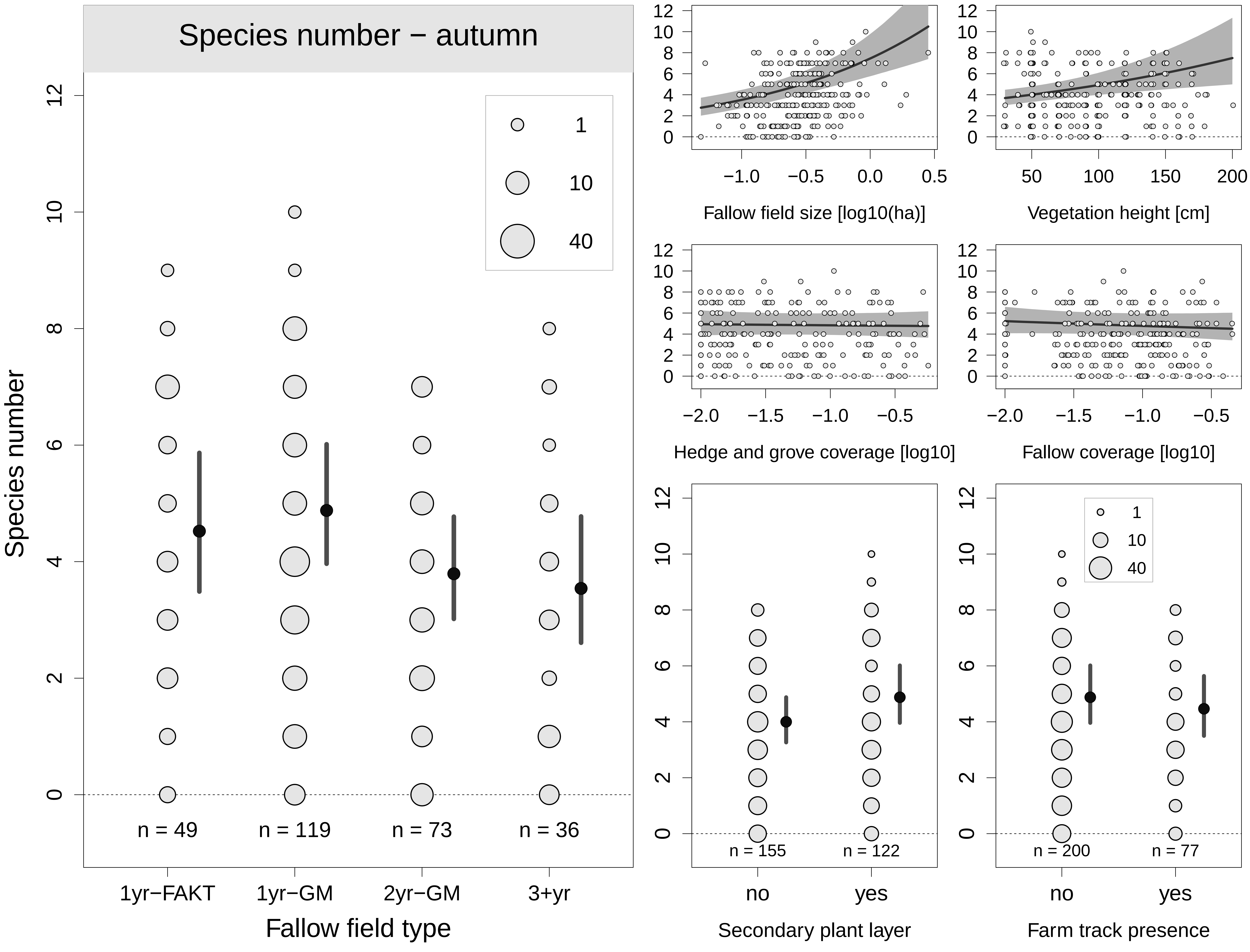
Species number − autumn
Species number
Fallow field type
1yr−FAKT
1yr−GM
2yr−GM
3+yr
n = 49
n = 119
n = 73
n = 36
1
10
40
Fallow field size [log10(ha)]
Vegetation height [cm]
Hedge and grove coverage [log10]
Fallow coverage [log10]
Secondary plant layer
Farm track presence
no
yes
n = 155
n = 122
n = 200
n = 77

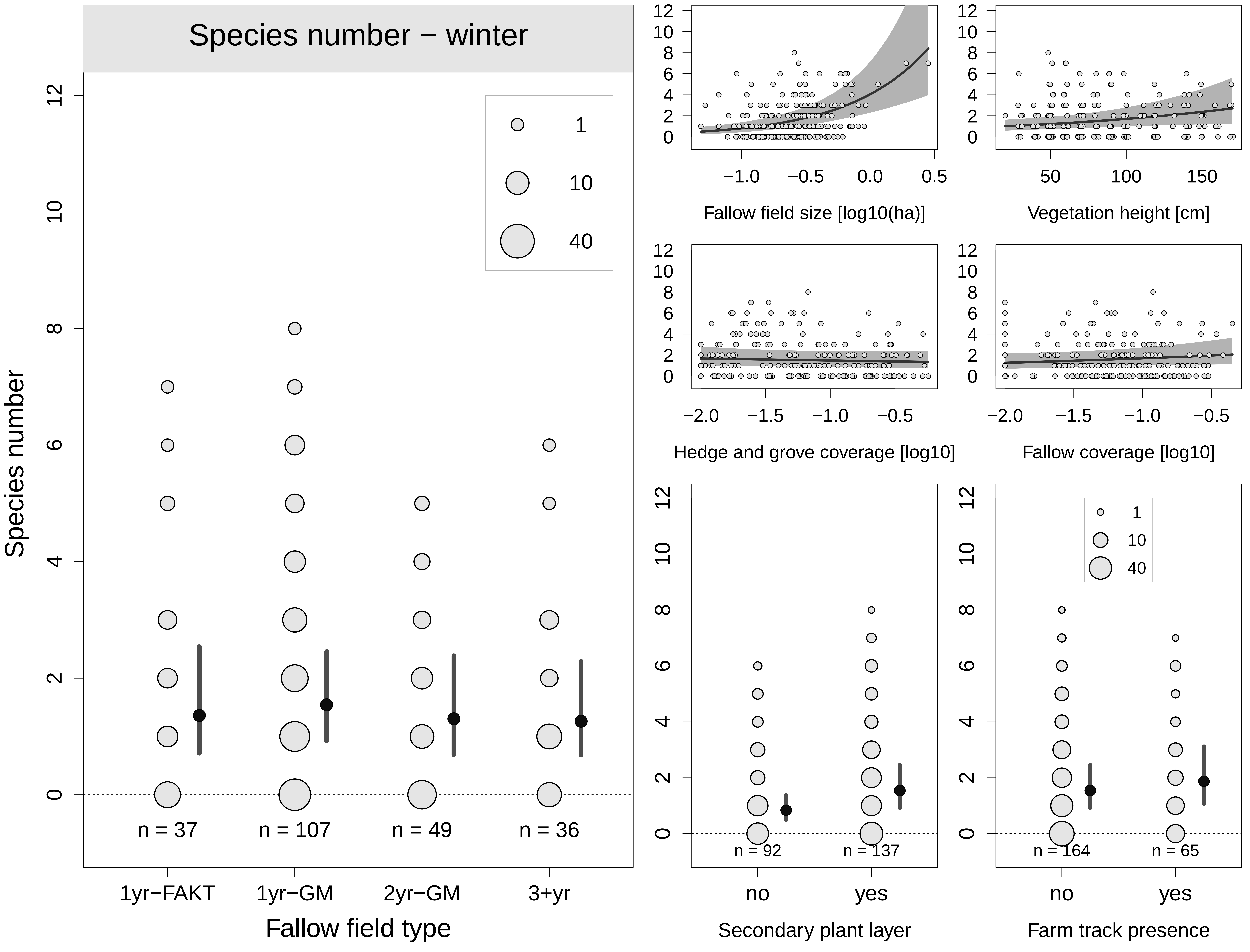
Species number − winter
Species number
1
10
40
n = 37
n = 107
n = 49
n = 36
1yr−FAKT
1yr−GM
2yr−GM
3+yr
Fallow field type
Fallow field size [log10(ha)]
Vegetation height [cm]
Hedge and grove coverage [log10]
Fallow coverage [log10]
n = 92
n = 137
no
yes
Secondary plant layer
n = 164
n = 65
no
yes
Farm track presence
